# Functional independence of entorhinal grid cell modules enables remapping in hippocampal place cells

**DOI:** 10.1101/2025.09.24.677985

**Authors:** Christine M. Lykken, Benjamin R. Kanter, Anne Nagelhus, Jordan Carpenter, Matteo Guardamagna, Edvard I. Moser, May-Britt Moser

**Author notes:** Equal contribution (Edvard I. Moser, May-Britt Moser).

## Abstract

A systems-level understanding of cortical computation requires insight into how neural codes are transformed across distinct brain circuits. In the mammalian cortex, one of the few systems where such transformations are tractable is the spatial mapping circuit. This circuit comprises interconnected regions of medial entorhinal cortex (MEC) and hippocampus, which encode location using fundamentally different neural codes. A key distinction is that neural activity in MEC, including that of directionally tuned cells and grid cells, evolves along low-dimensional manifolds, preserving stable phase relationships across different environments and behaviors^1–8^. In contrast, hippocampal place cells frequently undergo global remapping: their collective firing patterns reorganize randomly across different environments^9–12^, revealing an apparently limitless repertoire of orthogonal spatial representations^12–14^. The mechanisms by which spatial maps are transformed between the two coding schemes remain unresolved. Here, we used large-scale multi-area Neuropixels recordings to show that when rats were transferred from one familiar environment to another, each module of grid cells underwent a unique change in phase on its low-dimensional manifold, at the same time as simultaneously recorded place cells exhibited global remapping. In contrast, training conditions that produced smaller differences in the phase shifts of simultaneously recorded grid modules resulted in incomplete place cell remapping, mirroring previous reports of ‘partial remapping’^15–19^. Hippocampal remapping was not associated with rotational differences between grid modules under any condition. Taken together, these findings suggest that differential phase shifts across grid cell modules form the basis for the orthogonalization of downstream hippocampal spatial codes during remapping. The transformation from low-dimensional spatial representations in the MEC to high-dimensional codes in the hippocampus may underlie the hippocampus’ ability to support high-capacity memory storage^3,13,14,20–22^.

## Main

A central objective in neuroscience is to understand how neural codes are generated and transformed as signals propagate through interconnected neural circuits. Among the relatively few studies that have examined such transformations directly, most have focused on sensory systems, where conceptual frameworks exist for understanding the progression from peripheral receptor activation to cortical representation^23–27^. In contrast, far less is known about the representational transformations that support cognitive functions such as spatial navigation and memory, likely because these involve large populations of neurons distributed across higher-order cortical areas. While recent work in simple circuits of *Drosophila* has shed light on potential elementary mechanisms of code transformation in circuits for space and direction^28–33^, efforts to identify corresponding mechanisms in higher-order mammalian cortex have been hampered by a lack of population-level recordings from interconnected components of the relevant brain systems. With the advent of high-density Neuropixels probes, such data can now be acquired from behaving mammals^34,35^. In the present study, we leverage this technology to uncover the mechanisms that mediate the transformation between two well-characterized neural codes in the rodent spatial navigation circuit.

The positional coding system in rodents offers a unique window into how rigid and low-dimensional neural codes can be converted into sparse and combinatorially rich representations in the brain. Upstream in MEC, grid cells fire in a periodic, hexagonal pattern that tiles the entire environment available to a moving animal^36,37^. Along the dorsoventral axis of MEC, these cells are organized into modules, or sets of cells that share features such as grid spacing and grid orientation^38,39^. Within each module, population activity is confined to a low-dimensional toroidal manifold^8^, with cell-to-cell relationships preserved across behavioral states and experimental contexts^3,4,6–8,40^. These structured, invariant responses stand in contrast to the heterogeneous downstream activity of hippocampal place cells, which ‘remap’ each time the animal encounters a different environment^9,10^. During remapping, individual place cells exhibit pronounced and independent changes in their firing locations and firing rates, most often resulting in a complete orthogonalization of the population activity pattern, referred to as global remapping^11^. This decorrelation, in turn, yields an astronomic expansion in the number of possible neural activity configurations. As a result, the hippocampal code is often described as high-dimensional, with the capacity to store vast amounts of information^20,22^.

How the rigid, low-dimensional code in MEC is transformed into a rich set of orthogonal maps in the hippocampus remains an open question. Computational models have implicated grid cells as a key source of the place cell signal^21,41–47^ and the modular organization of the grid cell system^38,39^ has been identified as a possible basis for remapping^39^. One prominent hypothesis suggests that remapping arises when different grid modules realign in distinct ways with reference to environmental boundaries^3,21,48,49^. Such differential shifts in grid phase or orientation would alter patterns of coactivity between modules, thereby reshaping the input combinations received by downstream place cells, which would lead to remapping. In the present study, we set out to test this hypothesis.

To capture neural activity on both sides of the code transformation, we implanted Neuropixels probes in both the MEC-parasubiculum (PaS) and the hippocampus of 10 rats (Supplementary Table 1). An additional four rats were implanted with probes in MEC-PaS only. Between 488 and 1,531 cells were recorded per probe over a length of 2,880-3,440 µm (Extended Data Fig. 1). Across all 14 rats, we recorded a total of 13,785 cells in layers II-III of MEC-PaS, of which 2,107 were identified as grid cells. These grid cells were classified into 2-4 discrete modules per rat (Supplementary Table 1). Module classification was performed using the nonlinear dimensionality reduction algorithm UMAP^50^ and the DBSCAN clustering algorithm, as described previously^8,40^ (Extended Data Fig. 2).

Neural activity was recorded while rats foraged for food rewards in 150 or 200 cm-wide square open field arenas. In the first set of experiments, rats performed a ‘different-rooms’ task, exploring two highly familiar arenas located in rooms with distinct distal landmarks (rooms A and B; see Supplementary Table 1). In a second set of experiments, rats either explored a novel arena (‘novel-rooms task’) or experienced a rearrangement of local versus distal cues within a single arena (‘double-rotation task’; see Methods).

### Grid modules undergo distinct shifts in spatial phase

We began by analyzing grid cell activity during the ‘different-rooms’ task (11 rats; one experiment per rat). In this paradigm, hippocampal place cells typically undergo complete orthogonalization of their firing patterns (‘global remapping’)^3,11^. As expected^3^, grid cells shifted their firing locations relative to arena boundaries when animals transitioned between rooms A and B (Fig. 1a-b and Supplementary Table 2). Within each grid module, the phase shifts were coherent (Extended Data Fig. 3 and 4), in agreement with previous studies that recorded grid cells from anatomically confined regions of MEC^3,38,39^. Here, however, the use of Neuropixels probes enabled us to sample a broader dorsoventral extent of MEC-PaS, yielding simultaneous recordings from multiple grid modules.

**Fig. 1.**
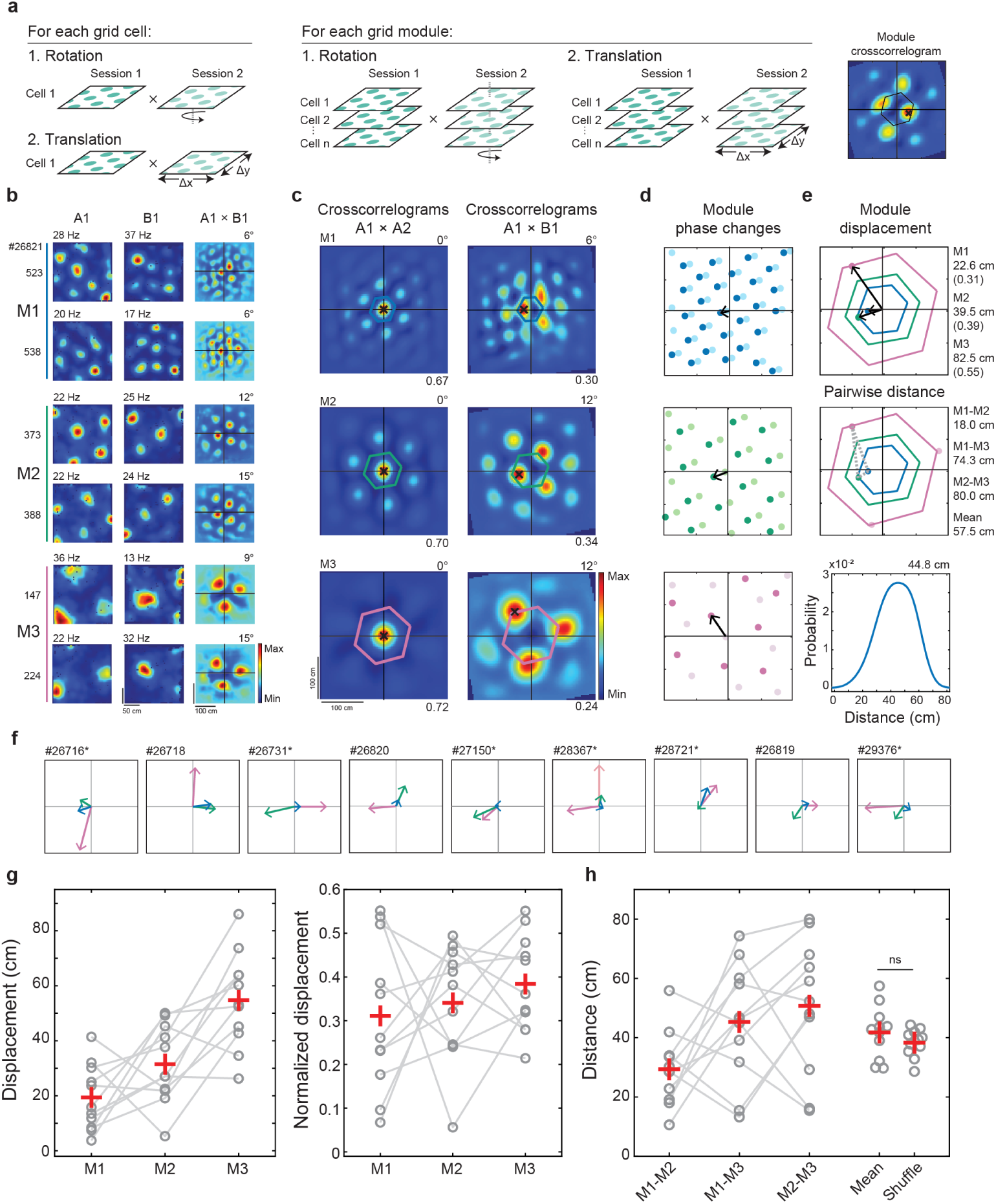
Grid modules undergo distinct changes in spatial phase between familiar rooms. **a**, Schematic of crosscorrelation procedure. Rate maps from each grid cell (left) or from all grid cells belonging to the same module (middle) were crosscorrelated between sessions in the same room (A1×A2) or different rooms (A1×B1). Rate maps from one session were rotated with respect to the other session in 3° increments and, for each rotation step, shifted along the entire x and y axis to generate a spatial crosscorrelogram (right). Note that the crosscorrelogram for a pair of grid maps is itself a grid pattern but it may be shifted from the origin. The rotation of each module was defined as the rotation with the highest correlation after shifting in x or y. We used the spatial crosscorrelogram at this rotation to determine the phase change of each grid cell or module within the inner unit tile of the crosscorrelogram (right panel, indicated by a colored hexagon). Black cross indicates location of peak correlation (see Extended Data Fig. 3). The spatial crosscorrelogram for an entire module was referred to as the module crosscorrelogram. **b**, Rate maps of six simultaneously recorded grid cells (one cell per row) from three grid modules in rooms A (left) and B (middle) in rat #26821. Color indicates firing rate (scale bar). Cell ID (3 digits) and module number (M1-M3) are noted on the left. Peak firing rate is noted above each panel. Crosscorrelograms for each grid cell (A1×B1) are shown in the rightmost column. Color indicates Pearson correlation (scale bar). Module rotation is noted above each panel. Note similar rotation and phase change for cells belonging to the same grid module, but different phase changes for cells from different modules. **c**, Panels show module crosscorrelograms for A1×A2 and A1×B1 in three modules. Black crosses indicate phase change of each module between rooms (i.e., the peak correlation within the unit tile at the rotation with the highest correlation; Extended Data Fig. 3b-c). Unit tile is indicated by a colored hexagon. Module rotation is noted above each panel. Maximum correlation is noted below each panel. Note that the phase shift differs between grid modules. **d**, Phase changes differ between grid modules (black arrows). Colored circles show locations of peaks in module crosscorrelograms comparing sessions in the same room (A1×A2, light colors) or different rooms (A1×B1, dark colors) at the rotation with the highest correlation. The phase change is represented by a vector (black arrow) that points from the origin to the location of the highest correlation within the unit tile of the module crosscorrelogram for A1×B1. **e**, Top, phase displacement is calculated for each grid module as the distance from the origin of the crosscorrelogram to the endpoint of each phase-change vector (black arrows, same as in **d**). Colored hexagons represent the central unit tile of the crosscorrelogram (shown also in **c**). Right, absolute displacement in cm, with normalized displacement (relative to module spacing) in parentheses. Middle, pairwise distance between endpoints of phase change vectors (dashed gray lines; from here on referred to as the pairwise distance between module phase changes) was used to quantify the difference in translation between modules. Distance (cm) between each module pair is noted at the right; mean distance between module pairs at the bottom. Note that the shortest distance between M2 and M3 is measured by wrapping around the axis of the hexagon to the pink point at the bottom of the unit hexagon for M3 (dashed line not shown; see Extended Data Fig. 3d). Bottom, grid patterns for each module were shifted randomly with respect to one another 1,000 times. For each iteration, the mean distance between the endpoints of the phase-change vectors was calculated for each pair of modules. Blue line shows distribution of pairwise distances for rat #26821. Peak of the distribution is noted above plot. **f**, Panels show phase-change vectors for simultaneously recorded grid modules in each animal (as in **d**, **e**). Animal ID is noted at top. Asterisks indicate animals with simultaneous recordings of hippocampal place cells. Nine animals are shown; the remaining two animals included in Fig. 1 are shown in Figs. 1e and 2b. Three modules were recorded from all animals, except rat #28367, which had four. **g**, Left, phase displacement (as in top panel of **e**) of simultaneously recorded grid modules in all 11 rats. Right, phase displacement of simultaneously recorded modules normalized by grid spacing in all 11 rats. Lines connect data from the same animal. Red crosses indicate means. **h**, Pairwise distance (cm) between module phase vectors (as in middle panel of **e**) for each pair of simultaneously recorded modules (all 11 rats). Mean across module pairs is shown for each animal next to the mean of the shuffled distribution for each animal (as in bottom panel of **e**). Red crosses indicate means. Four modules were recorded in rat #28367 (see panel **f**). Pairwise distances between M4 and the remaining modules are not shown here, but were included in the mean for this animal and are shown in Extended Data Fig. 5f. ns, not significant.

To compare spatial shifts across modules following the room change, we computed spatial crosscorrelograms^3^ (Fig. 1a and Extended Data Fig. 3a; see Methods). For each grid module, two-dimensional rate maps from individual grid cells were stacked into a three-dimensional (3D) matrix, with cell identity along the z-axis. This was done separately for each session. To compare a pair of sessions, we rotated one of the stacks in 3-degree increments from 0° to 360° (Fig. 1a; Extended Data Fig. 3a and 4a). At each rotation angle, we calculated pixel-wise population vector correlations^3^ of the two environments by shifting one stack in 3.75 cm steps along both the x and y axes. Note that the resulting crosscorrelogram for a pair of grid patterns is itself a grid pattern. The rotation angle that yielded the highest crosscorrelation across all spatial shifts was defined as the module’s rotation (Extended Data Fig. 3a). The matrix of correlation values across the xy space obtained at this rotation was termed the ‘module crosscorrelogram’. The phase shift (or ‘displacement’) of each module between rooms was defined as the location in the central hexagonal tile of the crosscorrelogram that showed the highest correlation (Fig. 1c and Extended Data Fig. 3b-c). If this location was offset from the origin, the grid map was considered shifted, and the distance and direction of the offset were noted (Extended Data Fig. 3d-e).

In support of our hypothesis, we found that simultaneously recorded grid modules exhibited markedly different phase shifts between environments (A1×B1; Fig. 1c-e; quantified in following paragraph). The magnitude and direction of module displacement varied substantially between simultaneously recorded modules as well as across animals (Fig. 1f-g). The average shift corresponded to approximately one-third of the module’s grid spacing (0.35 ± 0.06, mean ± s.e.m., 11 rats; mean displacement for each module: M1, 19.4 ± 3.6 cm; M2, 31.5 ± 4.3; M3, 55.5 ± 5.0; n = 11 rats; difference normalized relative to each module’s grid spacing: (F(2,30) = 0.88, p = 0.43, one-way ANOVA). Grid spacing and grid ellipticity did not change between rooms (spacing: F(1,30) = 1.3, p = 0.27; ellipticity: F(1,30) = 3.2, p = 0.08; one-within, one-between repeated measures ANOVAs). There was no change in module phase between repeated exposures to the same room (A1×A2: M1, 1.4 ± 0.76 cm; M2, 0.68 ± 0.45 cm; M3, 0.68 ± 0.45 cm; mean ± s.e.m., 11 rats).

To evaluate whether simultaneously recorded grid modules underwent distinct changes in grid phase, we calculated, within the central hexagonal tile, the Euclidean distance between the endpoints of the vectors that represented the change in phase for each module (from here on referred to as the pairwise distance; Fig. 1e, middle, and Extended Data Fig. 3d-e). Between sessions in different rooms, this distance ranged from 10.6 cm to 80.0 cm (n = 33 module pairs), with a mean of 41.8 cm (± s.d. of 8.9 cm, n = 11 rats; Fig. 1h). To determine how these differences in module translation compared to chance, we created a shuffled distribution for each pair of modules in each rat by randomly shifting the grid patterns of each module with respect to one another 1,000 times and calculating the distance between modules in the same manner as described above (Fig. 1e and Supplementary Methods). The mean pairwise distance between modules in our experiments was not significantly different than the mean distance between modules in this shuffled dataset (37.7 cm ± 1.4 cm; mean ± s.e.m.; t(10) = 1.7, p = 0.11, two-sided unpaired t-test; Fig. 1h). When we compared the distance between phase change vectors for each module pair to the shuffle for that pair, the pairwise distance between was smaller than expected by chance in only 1 of 33 module pairs (Extended Data Fig. 5a-b).

We validated this result using several complementary approaches that differed in how we defined the phase change for each module or in the rotation angle at which differences in phase change were measured (Extended Data Fig. 5). Estimates of module phase remained consistent across variations in the spatial bin size or the number of neurons per module (Extended Data Fig. 6). The grid pattern in the module crosscorrelogram was highly robust to contamination caused by the inclusion of cells with or without high spatial information content that did not meet the criteria for grid cells, or the inclusion of grid cells from other modules (Extended Data Fig. 6). Furthermore, grid modules exhibited similar phase changes upon re-exposure to the same environments (Extended Data Fig. 7a-d). The differential phase changes were stable across repeated exposures to the same room pairs within each day of recording (median difference in module phase change A1×B1 vs. A2×B2 = 3.75 cm, 95% CI, 3.75 - 5.30 cm, n = 30 modules) as well as across days of recording (median difference in module phase change A1×B1 on Day 1 vs. Day 2 = 5.30 cm, 95% CI, 3.75 - 8.39 cm, n = 21 modules; Extended Data Fig. 7a-d), even when one of the environments was initially novel to the animal (Extended Data Fig. 7e). Between different pairs of rooms, grid modules underwent unique realignments (Extended Data Fig. 7f). When the same grid modules were recorded simultaneously in both hemispheres, phase changes were coherent across hemispheres (Extended Data Fig. 7g). Changes in module phase across rooms were not accompanied by substantial changes in grid score, grid spacing, ellipticity of the grid pattern, or firing rate of grid cells (Supplementary Table 2). Taken together, these results demonstrate that simultaneously recorded grid modules undergo distinct changes in spatial phase between environments, without altering the metric structure of the grid pattern.

### Differential realignment of grid modules coincides with global remapping of place cells

We next asked whether the differential realignment of grid modules observed between familiar rooms consistently occurred in conjunction with global remapping of hippocampal place cells. In seven rats with a second Neuropixels probe targeting the hippocampus (Extended Data Fig. 1), we recorded a total of 1,617 cells in the hippocampus, including 416 place cells with distinct firing fields (Supplementary Table 1; see Methods for criteria). Simultaneous recordings were made from multiple grid modules and from place cells in CA1 and/or CA3 as the rats successively explored two familiar open-field arenas. As in the complete dataset with 11 rats, simultaneously recorded grid modules exhibited substantially different shifts in grid phase between the two rooms (pairwise distance in experiments with vs. without co-recorded place cells: t(10) = 0.54, p = 0.60; pairwise distance in experiments with co-recorded place cells vs. pairwise distance between shuffled module phases: t(14) = 0.08, p = 0.94; two-sided unpaired t-tests; Fig. 2a-b and Extended Data Fig. 8a).

**Fig. 2.**
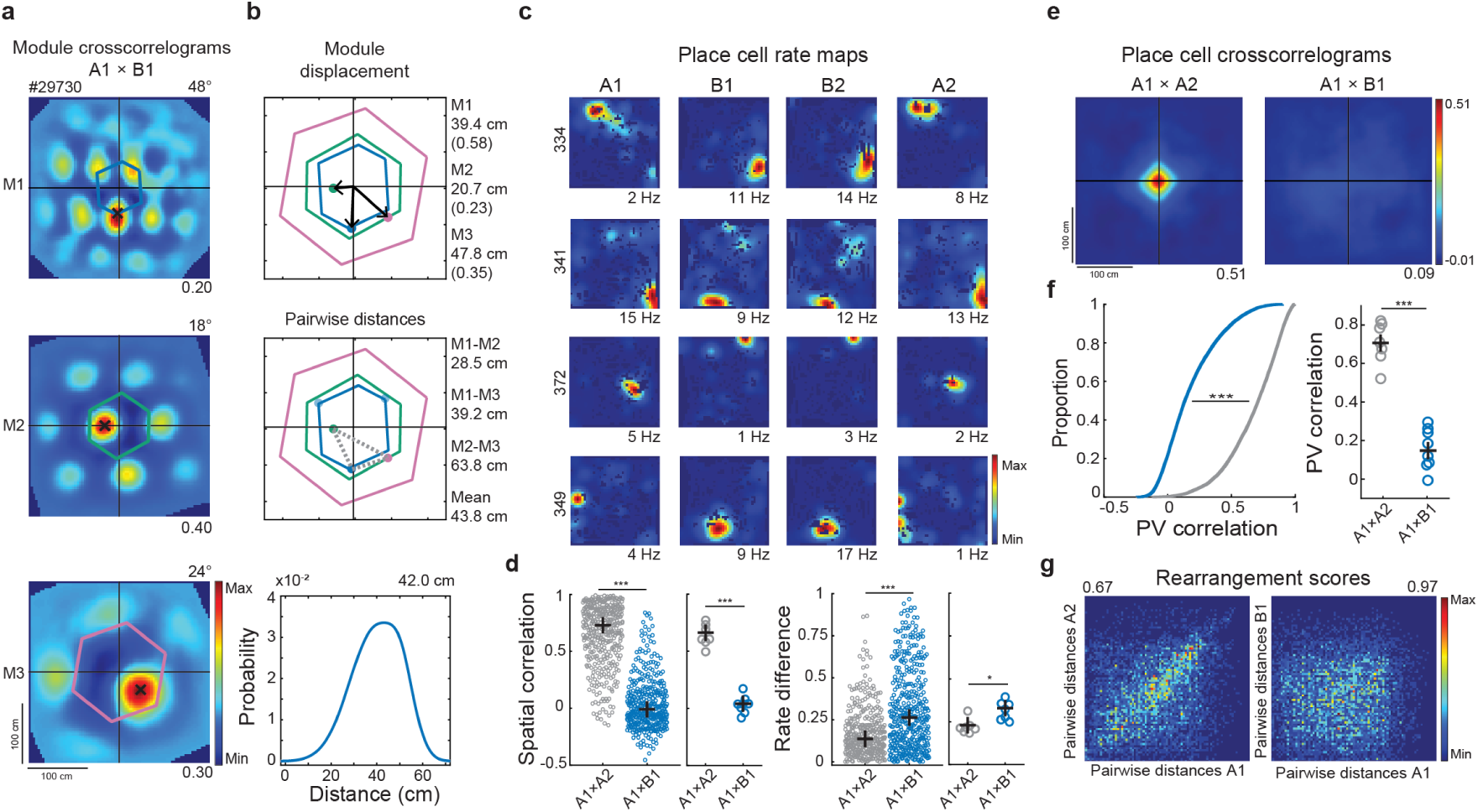
Differential changes in module phase coincide with global remapping of place cells. **a**, Crosscorrelograms comparing sessions in different rooms (A1×B1) shown for three simultaneously recorded grid modules in rat #29730 (as in Fig. 1c). Unit tile is indicated by a colored hexagon. Black crosses indicate the phase change of each module between rooms. **b**, Comparison of phase changes across modules shown in **a**, using the same measures as in Fig. 1e. Top: phase displacement for each grid module (black arrows); middle: pairwise distance between module phase change vectors (dashed grey lines); bottom: distribution of mean pairwise distances in shuffled data. Note that in in the middle panel, the shortest pairwise distance between M1 and M2 is achieved by wrapping around the axis of the hexagon to the blue point in the upper left corner of the unit hexagon for M1. **c**, Example place cell rate maps in rooms A and B (same session as in **a** and **b**). Firing rate is color-coded (scale bar). Cell ID (3 digits) is noted on the left. Peak firing rate is noted below each panel. Note remapping of firing locations between rooms A and B. **d**, Spatial correlation and rate difference (absolute change in mean firing rate) between sessions in the same room (A1×A2, gray) or in different rooms (A1×B1, blue). First and third panels show all identified place cells (n = 416) from 8 recordings with simultaneously recorded place cells and multiple (3 or 4) grid modules (7 rats). Black crosses indicate medians. Second and fourth panels show mean values across all place cells for each pair of sessions. Black crosses indicate means. ***p < 0.001, *p < 0.05. **e**, Place cell crosscorrelograms at the rotation with the highest correlation for sessions in the same room (left, A1×A2) or in different rooms (right, A1×B1) in rat #29730 (as in Fig. 1a, right). The central peak in the left crosscorrelogram indicates stable rate maps between sessions. The lack of a peak in the right correlogram is consistent with a complete change in the coactivity pattern (global remapping). **f**, Left, cumulative distribution function (CDF) shows the population vector (PV) correlation of place cells between sessions in the same room (gray, A1×A2) or different rooms (blue, A1×B1). Right, median PV correlation across all spatial bins for each pair of sessions. Black crosses indicate means. **g**, Place field rearrangement scores comparing sessions in the same room (left, A1×A2) or different rooms (right, A1×B1) in rat #29730. For all cells with activity in both sessions, the pairwise distance between place field locations in the first session is plotted against the pairwise distance between place field locations in the second session. Data are plotted as a 2D histogram; color indicates number of place field pairs per bin (from blue to red, scale bar). Rearrangement score is noted above each panel. Note the strong correlation along the diagonal in the left panel, indicating that place cells were stable between sessions in the same room. No clear relationship is evident when comparing sessions in different rooms (right panel), indicating that place cells randomly reorganized the locations of their place fields.

In each rat with simultaneous probes in MEC-PaS and hippocampus, differential translation (i.e., phase shift) of the grid modules invariably coincided with global remapping of hippocampal place cells in CA1 and/or CA3 (Figure 2c-g). Between environments, place cells exhibited pronounced changes in both firing location and firing rate that were consistent with an orthogonalization of hippocampal population activity (Fig. 2c-d). To quantify the extent of remapping, we applied several complementary measures. At the level of individual cells, the spatial correlation of place cell rate maps was significantly lower between sessions in different rooms (A1×B1) than between sessions in the same room (A1×A2) (all hippocampal place cells; Z = 16.6, p = 4.9 × 10^−62^, d = 2.4, one-sided Wilcoxon sign-rank test; Fig. 2d, left; see Supplementary Table 2 for medians and confidence intervals). This reduction in spatial correlation was accompanied by significant changes in the mean firing rates of individual cells, with some cells exhibiting activity in only one of the environments (Z = 8.9, p = 1.9 × 10^−19^, d = 0.69, one-sided Wilcoxon sign-rank test; Fig. 2d, right; see Supplementary Table 2). For cells that were active in both rooms, shifts in place field location were significantly greater between A1 and B1 than between A1 and A2 (Z = 12.3, p = 3.2 × 10^−35^, d = 0.85, one-sided Wilcoxon sign-rank test; see Supplementary Table 2).

This apparent orthogonalization of the spatial firing pattern was supported by measures based on joint population activity. We first computed spatial population vector (PV) crosscorrelograms using the same approach as for grid cells (Fig. 1a and Extended Data Fig. 3a; see Methods). When we compared sessions in the same room, a clear peak was observed at 0° at the origin of the crosscorrelogram (maximum PV correlation across all rotations and shifts ranging from 0.43 to 0.68; Fig. 2e, left and Extended Data Fig. 8b-c), indicating stable ensemble activity over time. In contrast, comparisons between different rooms yielded nearly flat crosscorrelograms, with substantially lower maximum PV correlations (ranging from 0.07 to 0.28 across all rotations and shifts; Fig. 2e, right and Extended Data Fig. 8b-c). The lack of a distinct peak confirms that place cells remapped heterogeneously between rooms with little coherence in shifts or rotations (PV correlation: A1×A2 vs. A1×B1, t(6) = 7.0, p = 2.1 × 10^−4^, one-sided paired t-test; see Supplementary Table 2). In a second pixel-based PV measure, place cell rate maps were again stacked by session, and population vectors of firing rates were defined for each spatial bin based on the distribution of mean firing rates across cells^11^. Comparing the entire set of population vectors between two trials (without rotating or shifting either stack) revealed high PV correlations within the same room (A1×A2) and significantly reduced correlations across rooms (A1×B1) (A1×A2 vs. A1×B1: t(6) = 7.5, p = 1.5 × 10^−4^, one-sided paired t-test; Fig. 2f; see Supplementary Table 2). Finally, a third population-level remapping metric – the rearrangement score^21^ – quantified changes in pairwise distances between place fields across sessions (see Methods). Rearrangement scores were significantly higher between A1 and B1 than between A1 and A2 (A1×A2 vs. A1×B1, t(6) = 8.5, p = 7.3 × 10^−5^, one-sided paired t-test; Fig. 2g and Extended Data Fig. 8b-c; see Supplementary Table 2), indicating substantial reorganization of place field locations between environments. Taken together, all metrics of remapping showed a clear association between the functional independence of grid modules and the global remapping of place cells.

Finally, the extent of remapping did not markedly differ between subregions of the hippocampus. Comparable changes were observed in both CA3 and CA1 across individual-cell measures (spatial correlations, firing rate changes, changes in place field location; Extended Data Fig. 8d-f) as well as in population-level metrics (Extended Data Fig. 8d-g).

### Hippocampal remapping weakens with reduced disparity between grid modules

We next asked whether the degree of independence between grid modules was linked to the orthogonalization of hippocampal place representations. If differential translation of grid modules contributes to global remapping, we predicted that when only some of the pairs of grid modules undergo distinct phase shifts, the resulting reorganization of the hippocampal place cell code would be incomplete.

To test this hypothesis, we examined all session pairs in our dataset (38 pairs from 10 rats, with 2-4 modules per recording) for evidence of incomplete remapping (see Supplementary Table 1). Beyond comparisons of distinct, familiar rooms (‘different-rooms task’, A×B; n = 19 session pairs), invariably associated with global remapping^3,11,12^ (Fig. 2c-g and Extended Data Fig. 8), we now included sessions in which one environment was novel (‘novel-room task’, familiar vs. novel, F×N; n = 6 session pairs) as well as cue rearrangement sessions within a single arena^16–18^ (‘double-rotation task’, standard vs. rotated, S×R; n = 13 session pairs; see Methods and Supplementary Table 1 for full description of experimental conditions). To quantify similarity in phase changes across simultaneously recorded grid modules, we measured the smallest pairwise distance between their phase-change vectors (Fig. 3a-b and Extended Data Fig. 3e). These values were normalized to the maximum possible phase distance to permit direct comparison across modules of different scales (Extended Data Fig. 3e).

**Fig. 3.**
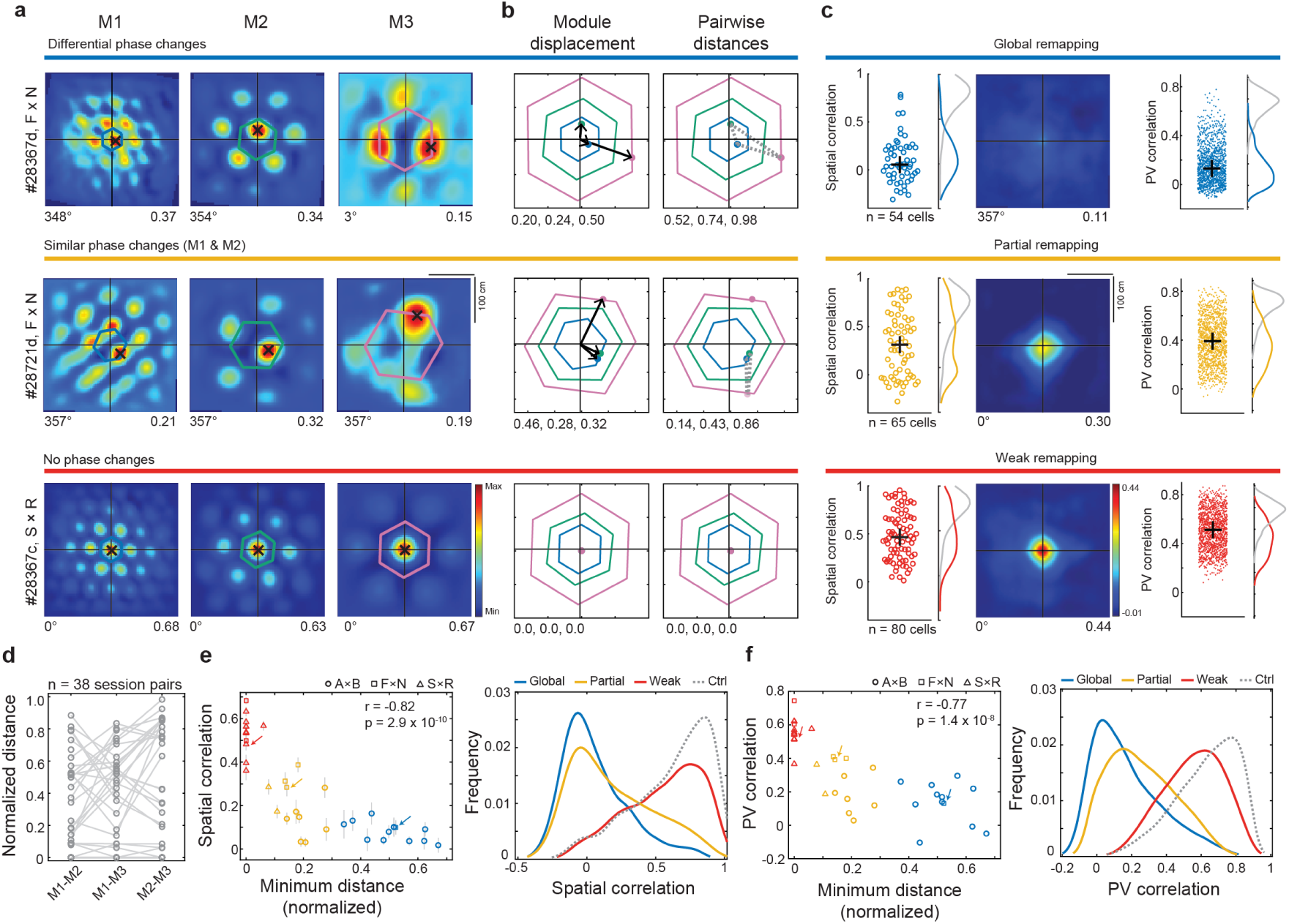
Strength of place cell remapping depends on degree of phase disparity across modules. **a**, Each row shows crosscorrelograms from simultaneously recorded grid modules between sessions when various degrees of remapping were observed in simultaneously recorded hippocampal place cells: global remapping in #28367d (top), partial remapping in #28721d (middle), and weak remapping in #28367c (bottom). Symbols as in Fig. 1c, right. **b**, Phase displacement (left column) and pairwise distance between phase change vectors (right column) for simultaneously recorded grid modules, as in Fig. 1e. Normalized displacement of each module (M1, M2, M3) is noted below panels in left column. Pairwise distances (M1-M2, M1-M3, M2-M3; normalized by the maximum possible distance) are noted below panels in right column. Note small pairwise distance for one module pair (M1-M2) in the recording with partial remapping. **c**, Remapping assessed using spatial correlation of rate maps and PV correlation in the three conditions (global remapping, partial remapping, and weak remapping). Left, circles show spatial correlation between sessions for each place cell in rats #28367d (n = 54 cells), #28721d (n = 65 cells), #28367c (n = 80 cells) (same rats as in **a** and **b**). Black crosses show medians. Curves at right show kernel smoothed density estimate of spatial correlation values between sessions in the same room (gray lines) versus different rooms (colored lines). Middle, each row shows PV crosscorrelograms at the rotation with the highest correlation between sessions for place cells recorded in the three examples. Color indicates Pearson correlation (scale bar). Rotation with highest correlation and maximum correlation are noted below each panel. Note the presence of a clear (but reduced) peak at the origin of the crosscorrelogram in the recording with partial remapping. Right, dots show PV correlation for each spatial bin in the three examples. PV correlations were calculated by stacking all place cell rate maps into a three-dimensional matrix with cell identity on the z-axis. The distribution of mean rates along the z-axis for a given x-y location (i.e., one spatial bin) represents the population vector for that location. Comparing the entire set of population vectors between two trials provides an estimate of how much the ensemble code changed between sessions. Black crosses show medians. Curves at the right show kernel smoothed density estimate of PV correlation between sessions in the same room (gray lines) versus different rooms (colored lines). **d**, Pairwise distance between phase change vectors for simultaneously recorded module pairs (n = 38 session pairs, as in Fig. 1e, middle). Values are normalized by the maximum possible distance between each module pair (see Extended Data Fig. 3e). Note that the distance values vary substantially, covering the full range from no difference to the maximum possible distance between modules (see Extended Data Fig. 3c). **e**, Left, scatter plot showing significant correlation between the mean spatial correlation of place cells (a measure of remapping) and the minimum distance between phase change vectors for simultaneously recorded modules (a measure of module coordination) (n = 38, r = −0.82, p = 2.9 × 10^−10^, Pearson correlation). Vertical lines show standard error of the mean. The data were divided into groups using *k*-means clustering. The number of clusters (*k* = 3) was determined based on criteria that maximized the separation of the clusters (see Extended Data Fig. 9a; Global, blue, n = 13; Partial, yellow, n = 12; Weak, orange, n = 13). Symbols indicate experimental conditions: Circle, different-rooms task, A×B; square, novel-rooms task, familiar (F) vs. novel (N); triangle, double-rotation task, standard (S) vs. rotated (R). Colored arrows indicate sessions shown in **a-c**. Right, panel shows frequency distribution of spatial correlation values of place cells in each group defined using k-means clustering (Global, blue, n = 672 cells; Partial, yellow, n = 816 cells; Weak, orange, n = 1682 cells) compared to a distribution of spatial correlation values between repeated sessions in the same room (Ctrl, stippled gray line, n = 2161 cells). The spatial correlation of place cells between sessions was significantly different between all four groups (Global vs. Partial, D = 0.18, p = 2.2 × 10^−11^; Partial vs. Weak, D = 0.47, p = 1.1 × 10^−105^; Weak vs. Ctrl, D = 0.16, p = 2.5 × 10^−35^; two-sided Kolmogorov-Smirnov tests) **f**, Left, scatter plot shows significant correlation between PV correlation (a measure of remapping) and the minimum distance between phase change vectors across sessions for simultaneously recorded modules (a measure of module coordination) (n = 38 session pairs, r = −0.77, p = 1.4 × 10^−8^, Pearson correlation). Session pairs are separated into groups as in **e**. Symbols indicate experimental conditions, as in **e**. Colored arrows indicate sessions shown in **a-c**. Right, panel shows frequency distribution of PV correlation between place cell rate maps for cells in each group defined using k-means clustering compared to a distribution of PV correlation values between repeated sessions in the same room (Ctrl, stippled gray line). The distributions of PV correlations were significantly different between all four groups (Global vs. Partial, D = 0.16, p = 2.4 × 10^−171^; Partial vs. Weak, D = 0.51, p = 4.4 × 10^−2497^; Weak vs. Ctrl, D = 0.26, p = 1.1 × 10^−4000^; two-sided Kolmogorov-Smirnov tests).

Consistent with our hypothesis, the extent of place-cell remapping scaled with the minimum pairwise distance between module phase changes (Fig. 3). To quantify this, we applied *k*-means clustering to the distribution of spatial rate-map correlations between environments A and B, as a function of the minimum distance between module phase changes (Extended Data Fig. 9a). The clustering separated the data into three groups (Fig. 3e). We refer to these groups using terms that qualitatively describe the form of remapping that was most common among the sessions in that group (see Extended Data Fig. 9b for experimental conditions in each group). When all simultaneously recorded grid modules shifted differentially (minimum distance 0.34 - 0.67), place cells showed robust remapping (‘global remapping’; Fig. 3a-c, top row; Fig. 3e-f, Extended Data Fig. 10, and Supplementary Table 3). When at least one pair of grid modules remained coordinated (minimum distance = 0.08 - 0.28), the remapping in place cells was typically incomplete, mirroring previous accounts of ‘partial remapping,’ where some cells showed strong changes in both firing locations and rates while the fields of other cells remained stable^15–19^ (‘partial remapping’; Fig. 3a-c, middle row; Fig. 3e-f, Extended Data Fig. 10, and Supplementary Table 3). Finally, when phase changes were near-coherent across all modules (minimum distance 0 - 0.06; Extended Data Fig. 9c), place cells underwent only small changes in spatial correlation between sessions (‘weak remapping’; Fig. 3a-c, bottom row; Fig. 3e-f, Extended Data Fig. 10, and Supplementary Table 3), reflecting changes in relative grid-cell firing rates^51,52^ rather than grid phase (Extended Data Fig. 9d). In all three groups, spatial correlation values were significantly reduced compared to repeated visits to the same room (Global vs. Ctrl, Z = 32.4, p = 2.3 × 10^−230^, d = 2.0; Partial vs. Ctrl, Z = 28.8, p = 1.7 × 10^−182^, d = 1.5; Weak vs. Ctrl, Z = 7.7, p = 6.3 × 10^−15^, d = 0.22; one-sided Wilcoxon rank-sum tests with Bonferroni correction).

The sensitivity to maintained phase relationships among subsets of grid modules was evident for every metric of place cell remapping that we employed, regardless of whether the estimates were based on individual cell or population measures (spatial correlation of rate maps: r = −0.82, p = 2.9 × 10^−10^; PV correlation (without rotation or translation): r = −0.77, p = 1.4 × 10^−8^; change in place field locations: r = 0.67, p = 8.5 × 10^−6^; PV correlation: r = −0.82, p = 3.6 × 10^−10^; rearrangement scores: r = 0.71, p = 7.7 × 10^−7^; Pearson correlations; Fig. 3e-f and Extended Data Fig. 10). There was no difference among the groups in the spatial correlation between the first and second half of either session (F(2,33) = 1.5, p = 0.12; one-within, one-between repeated measures ANOVA; Extended Data Fig. 10b), indicating that partial and weak remapping were not simply associated with a decrease in within-session place field stability. There was also no relationship between the extent of remapping and any of the following metrics related to grid cell activity: changes in mean firing rate, peak firing rate, spatial information, grid field size, grid spacing, grid ellipticity, or grid score (Extended Data Fig. 9e). In contrast, every metric that was used to quantify the disparity among the phase changes of grid modules showed a clear relationship with the extent of hippocampal remapping (Extended Data Fig. 9f). Taken together, these results reveal a clear link between phase shifts across grid modules and place cell remapping: the greater the disparity among module phase changes, the more pronounced the remapping of the hippocampal place code.

### Hippocampal remapping is not explained by rotational differences between grid modules

Theoretical work has pointed to both differential phase shifts and differential rotation of grid modules as potential mechanisms of remapping^21^. Our experimental data, however, provided no evidence supporting a role for rotational differences. Grid fields rotated coherently relative to external boundaries not only within individual modules^3,53^ (Fig. 1b and Extended Data Fig. 4), but also between modules (n = 104 module pairs; median difference between modules = 3°, 95% CI, 0 - 3°; Fig. 4a-d and Extended Data Fig. 11a-c). This coherent rotation across environments indicates that grid orientation is not independently determined for each module, but is instead governed by a shared mechanism, such as anchoring to global directional or environmental cues. This shared rotational alignment extended to all other space- and direction-tuned cells in the dataset, whose firing patterns consistently rotated in concert with the grid cells (Fig. 4e; Extended Data Fig. 11d-g).

**Fig. 4.**
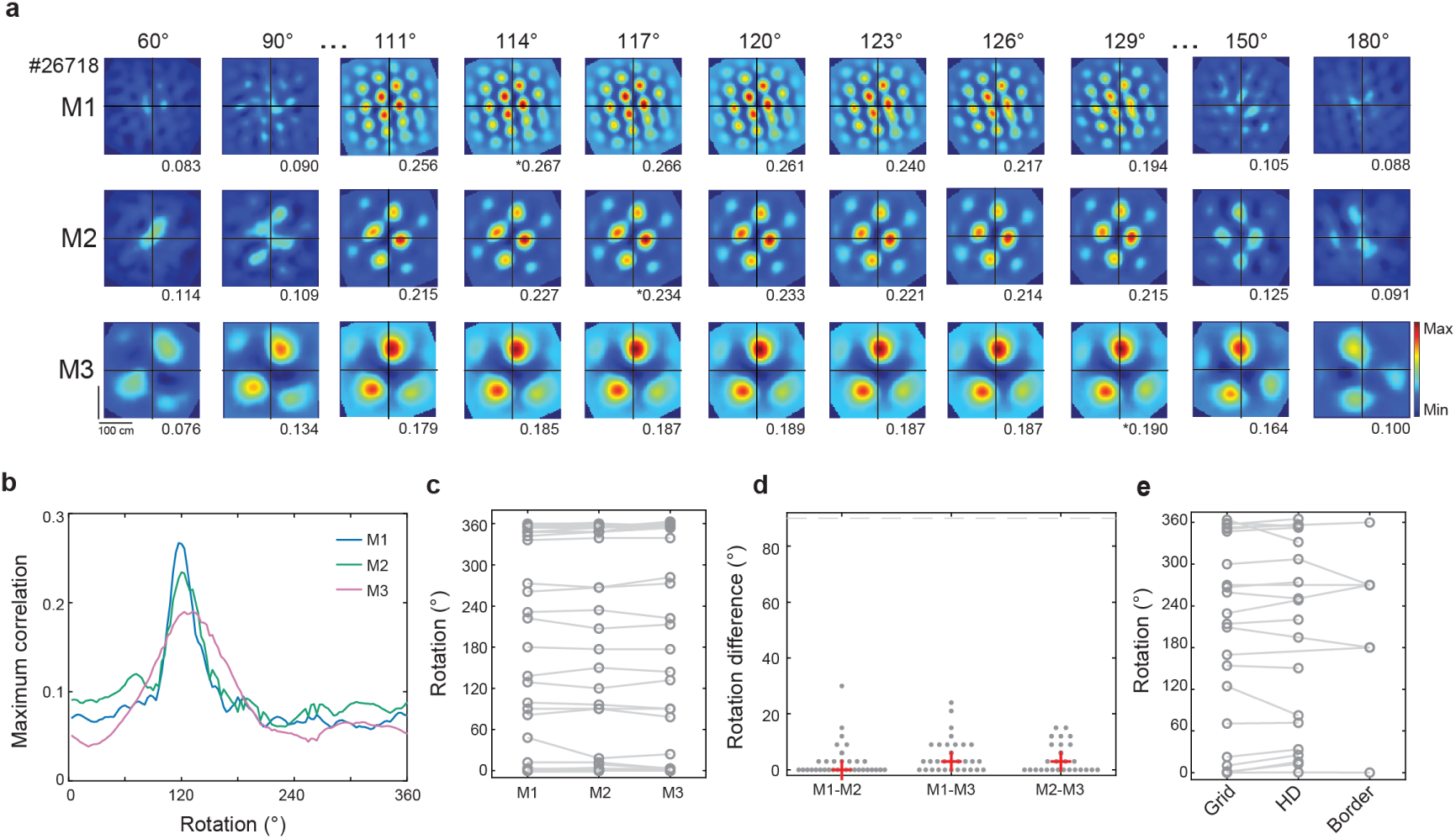
Coherent rotation of grid modules. **a**, Module crosscorrelograms for sessions in different rooms (A1×B1) at incremental degrees of rotation in rat #26718 (select number of rotations shown here for visualization purposes; see Extended Data Fig. 4 for additional rotations) (as in Fig. 1c, right). Asterisks denote overall maximum for each module. **b**, Maximum correlation between 3D stacks of rate maps of sessions A1 and B1 (with cell identity along the z-axis) as a function of rotation for each module in rat #26718. Maximum correlation refers to the highest value within the crosscorrelation matrix at each rotation. Note that the rotation was nearly identical across modules. **c**, Rotation of each simultaneously recorded module between sessions (38 session pairs in 10 rats, as in Fig. 3). Lines connect data from a single recording. A fourth module was recorded in 4 session pairs. Its rotation did not differ from the other simultaneously recorded modules (Extended Data Fig. 11j). **d**, Difference in rotation between simultaneously recorded module pairs (n = 104 module pairs, each pair indicated by a circle). Red crosses indicate medians. Dashed gray line at 90° represents chance. **e**, Gray lines show coherent rotation between sessions for grid, head direction (HD), and border cells (n = 22 experiments). Lines connect data from a single recording.

Although grid maps often rotated when rats were moved between rooms, such rotation relative to external boundaries was not required for global remapping to take place. For example, there was no significant correlation between the degree of grid module rotation and the spatial correlation values of place cells in the ‘global remapping’ condition (n = 13 session pairs; r = −0.38, p = 0.19, Pearson correlation; Extended Data Fig. 11h). Rotation of the grid pattern was less frequent in sessions where place cells underwent partial or weak remapping (Extended Data Fig. 11h-i). Importantly, the extent of rotation did not differ noticeably between grid modules, regardless of the strength of remapping (pairwise differences: Global, median = 6°, 95% CI, 3 - 6°; Partial, median = 6°, 95% CI, 3 - 9°; Weak, median = 0°, 95% CI, 0 - 0°; Extended Data Fig. 11i). Together, these results indicate that rotation differences between grid modules are unlikely to be the driving factor behind the orthogonalization of the hippocampal place code.

### Differential realignment of grid modules in a grid-to-place cell simulation

Finally, we wanted to establish whether differentially shifting and/or rotating grid modules is sufficient to produce hippocampal remapping. To address this question, we adapted a winner-take-all grid-to-place cell model^54^ by separating the 10,000 grid cell inputs to place cells into three modules with spacing values that closely matched our empirical data (Fig. 5a). These simulated grid modules were used to produce a set of simulated hippocampal place cells with realistic place fields (Session A; Extended Data Fig. 12b). During each run of the simulation, we shifted and/or rotated the spatial firing patterns of grid cells in each module and generated a second set of rate maps for the hippocampal place cells using these realigned grid inputs (Session B; Fig. 5b and Extended Data Fig. 12). We applied the same shift and/or rotation to all grid cells within a module, whereas across modules, the realignment could vary. This approach enabled us to compare the extent of modular realignment, the degree of coordination among module pairs, and the amount of remapping among simulated place cells directly with our in vivo data.

**Fig. 5.**
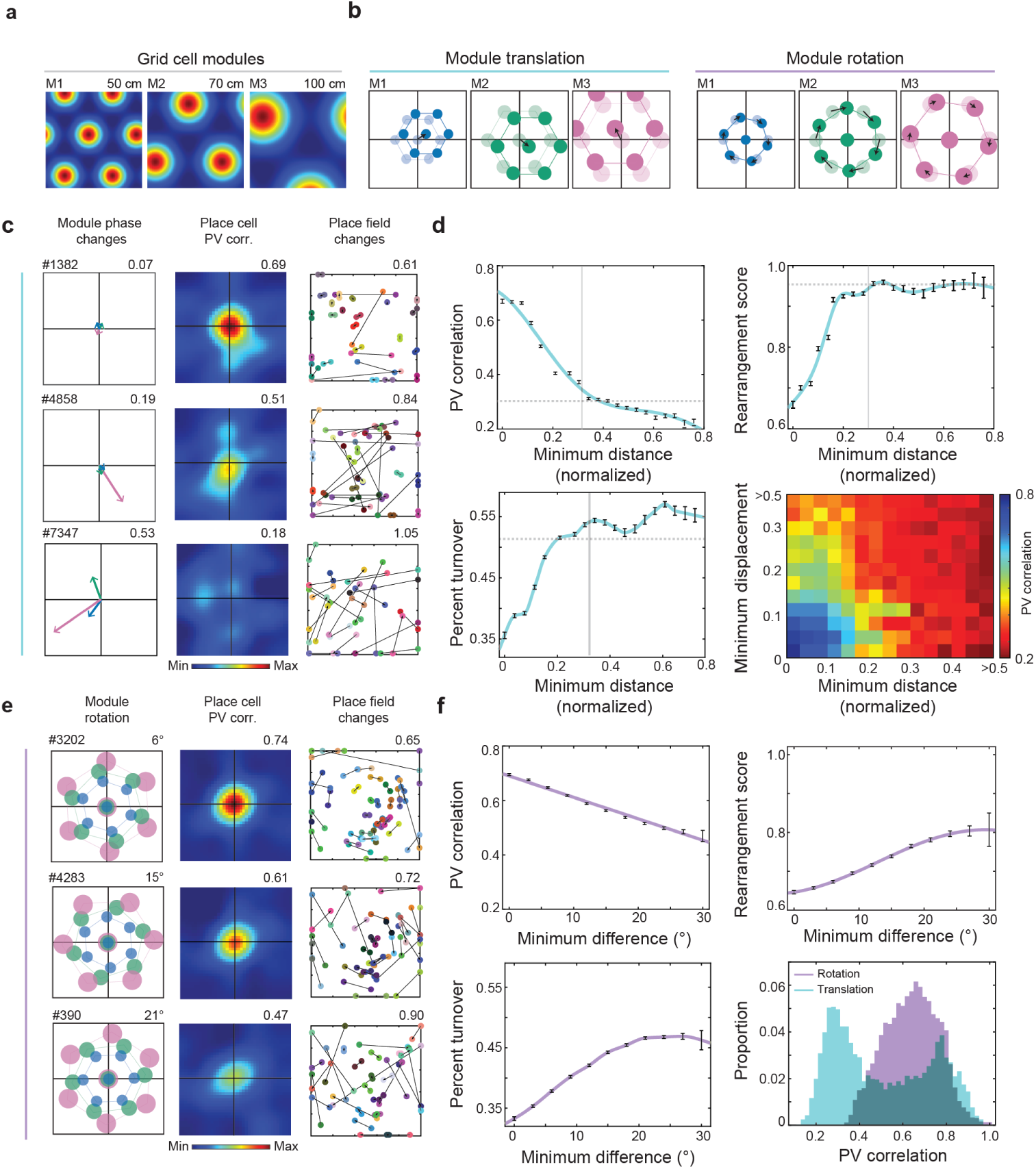
Differential realignment of grid modules produces place cell remapping in a grid-to-place cell simulation. **a**, Rate maps of three simulated grid cells. Color indicates firing rate from blue to red. Module number and grid spacing are noted above each panel. **b**, Schematic representations of module translation (left) and module rotation (right). During each simulation, the spatial pattern of each grid module underwent translation and/or rotation. For simulations of module translation, the grid pattern of each module was shifted separately. Black arrows indicate the change in module phase between sessions. For simulations of module rotation, the grid pattern of each module was rotated separately. Black arrows depict the rotation of each module between sessions. **c**, Left, schematic representation of changes in module phase during simulations of module translation (colored arrows depict the phase-change vector for each module, as in Fig. 1f; from top: simulation #1382, n = 28 cells; simulation #4858, n = 32 cells; simulation #7347, n = 23 cells). Minimum normalized distance between phase change vectors (as in Fig. 3e) is noted above each panel. Note increasing difference in phase change from top to bottom. Middle, population vector (PV) crosscorrelograms at the rotation with the highest correlation for place cells during each simulation of grid module translation. Maximum PV correlation is noted above each panel. Note decreasing PV correlation with increasing difference in phase translation. Right, black lines show the change in field location for each simulated place cell following translation of module phases. Different cells have different colors. Rearrangement score is noted above each panel. **d**, PV correlation (top left), rearrangement score (top right), and percent turnover (bottom left) as a function of the minimum normalized distance between phase change vectors of modules. Note that maximum remapping is obtained even with small differences in phase shifts. Error bars show mean ± s.e.m. Solid vertical lines indicate median normalized distance of a distribution created by randomly shuffling the grid patterns of each module with respect to one another. Dashed horizontal lines indicate medians of the shuffled distributions of PV correlations (top left), rearrangement scores (top right), or percent turnover (bottom left) for simulated place cells. Bottom right, heat map showing that the PV correlation of simulated place cells between sessions varies as a function of the minimum distance (normalized) between module phase changes (x-axis) and the minimum displacement (normalized) relative to the origin (y-axis; see Extended Data 3c). Color indicates PV correlation from blue to red. **e**, Left, schematic representation of module phases during representative simulations of module rotation (from top: simulation #3202, n = 43 cells; simulation #4283, n = 35 cells; simulation #390, n = 33 cells). Minimum rotation difference between all pairs of modules is above each panel. Middle, PV crosscorrelograms at the rotation with the highest correlation for place cells during each simulation of module rotation. Maximum PV correlation is noted above each panel. Right, black lines show the change in field location for each simulated place cell when grid cell rate maps for each module were rotated as indicated in the left column. Rearrangement score is noted above each panel. **f**, PV correlation (top left), rearrangement score (top right), and percent turnover (bottom left) as a function of minimum rotation difference between modules. Note that the strength of remapping increases as the angular difference between modules increases, but that remapping is still less pronounced than after differential phase translation. Error bars show mean ± s.e.m. Note that the minimum difference in rotation across modules in experimental data was 0°. Bottom right, distribution of PV correlations in simulated place cells, shown as the proportion of simulations involving translation (green) or rotation (purple) of grid modules.

In each simulation, we quantified the degree of independence between grid modules following translation or rotation, as well as the extent of remapping in place cells, using the same metrics as in the experimental data. To capture the extent to which modules shifted independently in space and orientation, we calculated, for each pair of grid modules, the minimum distance between changes in module phase, and the minimum difference in module rotation. To assess remapping in the place-cell population, we computed PV correlations and realignment scores, as in the experimental data, as well as the turnover in the active cell population (i.e., proportion of cells that were active in only one session).

There was a strong correspondence between the simulations of module translation and our in vivo data. In the simulations, changes in the phase relationship of grid modules were sufficient to produce hippocampal remapping, and the difference in module phase changes reliably predicted the extent of remapping (Fig. 5c-d and Extended Data Fig. 12b). Place cell remapping (as measured by PV correlations, rearrangement scores, and percent turnover) was most pronounced when modules exhibited differential phase changes (minimum normalized distance > 0.30, n = 2,345 simulations). Under these conditions, the PV correlation was low (median = 0.290, 95% CI, 0.286 - 0.294), while rearrangement scores and turnover rates were high (rearrangement score: median = 0.959, 95% CI, 0.955 - 0.965; percent turnover: median = 0.554, 95% CI, 0.547 - 0.556; Fig. 5c-d and Extended Data Fig. 12b). The values closely matched the threshold for global remapping identified in our in vivo recordings (Fig. 3a,e-f). The extent of remapping was similar to that of a shuffled dataset comparing the rate maps of place cells selected randomly from sessions A and B (Extended Data Fig. 13a). In contrast, when the minimum distance between modules was below 0.30 – but still above the 5^th^ percentile of a shuffled distribution (n = 6,244 simulations) – remapping was more modest. PV correlations were higher (median = 0.558, 95% CI, 0.547 - 0.567), while rearrangement scores and turnover rates were lower (rearrangement score: median = 0.870, 95% CI, 0.866 - 0.876; percent turnover, median = 0.481, 95% CI, 0.474 - 0.482; Fig. 5c-d and Extended Data Fig. 12b), mirroring the partial remapping we observed in vivo. Finally, when the normalized minimum distance between modules was less than expected by chance (< 0.08; n = 2,555 simulations), place cell remapping was minimal (PV correlation, median = 0.737, 95% CI, 0.729 - 0.744; rearrangement score, median = 0.697, 95% CI, 0.682 - 0.710; percent turnover, median = 0.393, 95% CI, 0.387 - 0.404; Fig. 5c-d). In line with these findings, we observed strong and significant correlations between the degree of module independence and the extent of place cell remapping (minimum distance vs. PV correlation: r = −0.93, p = 3.9 × 10^−9^; minimum distance vs. rearrangement score: r = 0.79, p = 2.9 × 10^−5^; minimum distance vs. percent turnover: r = 0.83, p = 4.8 × 10^−6^; Pearson correlations; Fig. 5d).

The degree of remapping depended not only on phase change differences between modules, but also on the magnitude of phase displacement between modules (i.e., how far the spatial grid pattern of each module shifted from the origin, normalized by its spacing; Fig. 5d and Extended Data Fig. 13b). The degree of displacement required to produce remapping increased as the phase distance between modules decreased, indicating an inverse relationship between these two variables (Fig. 5d). To achieve the most complete remapping, all three modules had to be shifted independently (Extended Data Fig. 13c).

Finally, the simulations showed that module translation was significantly more effective at producing remapping than module rotation (n = 9,261 simulations; Fig. 5e-f and Extended Data Fig. 13a), as shown previously^21^. As the angular difference between modules was increased beyond what was ever observed empirically, the extent of decorrelation in the simulated place cell population increased, but the remapping remained incomplete (Fig. 5e-f and Extended Data Figs. 12b, 13a). Combining rotation and translation in a separate set of simulations did not increase remapping beyond what was observed with translation alone (n = 13,500 simulations; Extended Data Fig. 13a, d-e). These results suggest that the primary determinant of hippocampal remapping is the differential translation of grid cell modules.

## Discussion

Using Neuropixels probes to record simultaneously from large numbers of grid cells across multiple modules, along with hippocampal place cells, we provide experimental evidence that differential phase translations among grid cell modules underlie the formation of multiple independent place-cell maps in the hippocampus. Supporting a causal relationship, the degree of place cell remapping – ranging from near-complete orthogonalization of spatial activity (global remapping) to weaker decorrelation of hippocampal representations (partial or weak remapping) – was strongly dependent on the disparity in phase shifts across grid modules. When animals transitioned between highly familiar arenas located in different rooms, we observed global remapping of hippocampal place cells, consistent with previous findings^11,12^. This orthogonalization of spatial activity patterns invariably coincided with differential phase translation across grid modules. In contrast, when phase changes were coordinated across a subset of modules, the hippocampal response was more heterogeneous: some place cells altered their firing locations, while others maintained their firing fields (partial remapping). Even weaker remapping occurred when module phases remained stable and only firing rate relationships changed. Finally, rotations of the grid pattern were always coherent across modules, suggesting that differential rotation is not required for driving remapping in downstream place cells.

Our experimental results were replicated in network simulations of the grid-to-place cell transformation. A simple model connecting idealized modules of grid cells to downstream place cells reproduced the full spectrum of remapping patterns observed in vivo. The results confirm longstanding theoretical predictions proposing that a small number of grid modules – each constrained by low-dimensional continuous attractor dynamics^8,42,43,55,56^ – can be flexibly recombined to a high-dimensional multimap spatial code in the hippocampus^3,21,48,49^. When the disparity between phase translations among subsets of modules was reduced – the model produced partial remapping. The distinctiveness of each hippocampal map likely depends not only on the phase relationship between modules but also on both the underlying connectivity (e.g., the probability of inputs from different modules to each place cell) and the strength of those connections. The balance between these factors remains to be determined.

Together, our results offer a mechanistic framework for understanding not only global remapping, but also partial remapping, a phenomenon observed under a variety of experimental conditions, including double rotations of local and distal cues, rearrangements of cue configurations, and changes in the task demands or reward contingencies^16–19,57^. The extent of remapping also varies across animals^58^. Our observations suggest that while the activity within individual grid modules remains rigid and low-dimensional, differential coordination between module pairs enables the emergence of intermediate representational states in the hippocampus. These intermediate states may be stable – incorporating elements from multiple previously experienced environments^16–19,57,59–61^ – or transient, marking transitions between distinct spatial representations^60,62,63^. The stability of these states likely depends on whether the disparities between modules are persistent or short-lived.

The findings point to a potential organizing principle of cortical computation: the construction of high-capacity, adaptive representations from the combination of a limited set of structured, low-dimensional representations. We identify a potential mechanism by which grid-like representations in the entorhinal cortex can be recombined in a context-dependent manner to generate novel patterns of coactivity in the hippocampus, dramatically expanding the system’s representational capacity, in line with its established role in episodic memory formation^64–67^. This reconfiguration allows the hippocampal network to encode a vast array of distinct spatial or non-spatial contexts while preserving the universal metric structure of the grid code through the interconnected architecture of entorhinal grid cells. In this integrated system of low and high-dimensional neural codes, the grid cell network may serve as a stable, reusable scaffold that – through independent phase alignment among grid modules – supports the flexible formation and recall of episodic memories^22,67^.

## Data availability

The datasets generated during the current study are available at EBRAINS, DOI: https://doi.org/ placeholder (will-be-provided-at-proofs-stage).

## Code availability

Code for reproducing the analyses in this article are available at Zenodo, DOI: placeholder (will-be-provided-at-proofs-stage).

## Acknowledgements

We thank B.A. Dunn, Y. Burak, Y. Gronich, and M. Witter for discussion; S. Ball, K. Haugen, E. Holmberg, K. Jenssen, E. Kråkvik, and H. Waade for technical assistance; the veterinary staff for animal care; R. Gardner and V.A. Normand for initial training with Neuropixels; and Torgeir Waaga for performing the Neuropixels implantation in one animal. The work was supported by a Synergy Grant to E.I.M. from the European Research Council ERC (‘KILONEURONS’, Grant Agreement No. 951319); two Centre of Excellence grants to M.-B.M. and E.I.M. (Centre of Neural Computation, grant number 223262; Centre for Algorithms in the Cortex, grant number 332640); a National Infrastructure grant to E.I.M. and M.-B.M. from the Research Council of Norway (NORBRAIN, grant number 295721); as well as a grant from the Kavli Foundation (M.-B.M. and E.I.M.), a direct contribution to M.-B.M. and E.I.M. from the Ministry of Education and Research of Norway.

## Author Contributions

C.M.L., B.R.K., A.N., E.I.M. and M.-B.M. planned and designed experiments, conceptualized and planned analyses, and interpreted data; C.M.L., B.R.K., A.N., J.C., and M.G. performed experiments; C.M.L. and B.R.K. visualized and analyzed data; C.M.L. curated the data; C.M.L. wrote the first draft of the paper with periodic input from B.R.K.; C.M.L. and E.I.M. edited the paper with periodic input from B.R.K. and M.-B.M. All authors discussed and interpreted data. M.-B.M. and E.I.M. supervised and funded the project.

**Supplementary Information** is available for this paper.

## Methods

### Subjects

Data were obtained from 17 adult male Long Evans rats. The rats were 11-16 weeks old and weighed 350-450 g at the time of implantation. Prior to surgery, the rats were group-housed with their siblings; thereafter they were singly housed in large, two-story enriched cages (95 x 63 x 31 cm). The rats were handled daily. They were kept on a 12-h light / 12-h dark schedule with strict control of humidity and temperature and free access to food and water. Experiments were performed during the dark phase of the schedule. All procedures were performed in accordance with the Norwegian Animal Welfare Act and the European Convention for the Protection of Vertebrate Animals used for Experimental and Other Scientific Purposes. Protocols were approved by the Norwegian Food Safety Authority (FOTS ID 18011 and 29893).

### Surgery and electrode implantation

The rats were implanted with one to three single-shank (phase 3A or version 1.0) or multi-shank (version 2.0) 384-channel Neuropixels probes^34,35^ (Supplementary Table 1). Probes were targeted to MEC-PaS in all 17 animals. Two of the rats (#26820 and #26821) were implanted bilaterally in MEC-PaS. Three of the rats (#26880, #26823, and #27207) were included only in Fig. 4 because the probe tracks were outside MEC layer II/III (Extended Data Fig. 1; see Methods). In 10 of the 14 rats with probes inside MEC layer II/III, one or two additional probes were targeted to the hippocampus (both CA3 and CA1). Four of the 10 rats (#28739, #29376, #29730, and #29731) had a third probe in the subiculum (data not included). The type of Neuropixels probe and the hemisphere of implantation are noted in Supplementary Table 1.

Before implantation, the rats were anaesthetized with 5% isoflurane in a pre-filled induction chamber. After subcutaneous injections of buprenorphine (Temegesic, 0.03 mg/ml) and Meloxicam (Metacam, 2.0 mg/ml), the rats were fixed in a Kopf stereotaxic frame. Continuous administration of 1.0-2.5% isoflurane was delivered through a nose cone. A subcutaneous injection of bupivacaine (Marcain, 2.5 mg/ml) was administered prior to an incision through the scalp. Craniotomies were made above MEC and/or the hippocampus and the subiculum. To target the MEC, the probes were centered at 4.5 mm lateral to bregma, positioned 0.1 mm anterior to the transverse sinus, and inserted at an angle of 26° from posterior to anterior in the sagittal plane to a maximum depth of 4.5 – 5.0 mm relative to the brain surface. To target the hippocampus, the probes were centered at 3.8 mm posterior to bregma and 3.2 mm lateral to bregma, and lowered vertically to a depth of 6.0 – 6.2 mm relative to the brain surface. The orientation of probe shanks relative to the midline is noted in Supplementary Table 1. In one animal (#29376), the probe was implanted at an oblique angle (40°) relative to the midline (medial shank was most anterior) and inserted an angle of 30° from anterior to posterior in the sagittal plane. A single stainless steel screw was secured to the skull above the cerebellum and connected to the probe ground with an insulated silver wire. See previously described procedures^8,35^ for further details of probe implantation. After surgery, the rats were placed in a heating chamber to recover until resuming normal locomotor behavior. Postoperative analgesia (Meloxicam and buprenorphine) was administered subcutaneously during the surgical recovery period.

### Recording procedures

The details of the Neuropixels hardware system and procedures for recording in freely moving animals have been described previously^8,35^. In brief, electrophysiological signals were amplified with a gain of 500 (for phase 3A probes) or 80 (for 1.0 and 2.0 probes), low-pass-filtered at 300 kHz (phase 3A) or 0.5 kHz (1.0 & 2.0), high-pass-filtered at 10 kHz, and then digitized at 30 kHz on the probe circuit board. The signal was multiplexed by an implant-mounted headstage circuit board and transmitted to a Xilinx Kintex 7 FPGA board (phase 3A) or a Neuropixels PXIe acquisition module (1.0 & 2.0) along a lightweight 5-m tether cable made using either micro-coaxial (phase 3A) or twisted pair (1.0 & 2.0) wiring. The signal was then streamed via ethernet connection to the local computer.

### Behavioral tracking

To track the rat’s head position and orientation during recording, we attached a removable rigid body with five retroreflective markers to the implant. These markers were tracked at 120 Hz with a 3D motion capture system (six overhead OptiTrack Flex 13 cameras and Motive recording software). The 3D marker positions were projected onto the horizontal plane to yield the rat’s 2D position and head direction. ‘South was defined based on a fixed point common to all recording rooms, which was where the animal was released into the box and was the point closest to the door of the recording room. Digital pulses generated by an Arduino microcontroller were used to synchronize the timestamps of the Neuropixels acquisition system (via direct transistor-transistor logic input) and the OptiTrack system (via infrared light-emitting diodes).

### Behavioral procedures

Over a period of five to ten days prior to surgery, the rats were familiarized with the experimenter(s) and one or more of the recording rooms. During each day of familiarization, the rats were gently handled and placed on a towel within a terra cotta flowerpot on top of a 1-m pedestal with a clear view of the open field arena, the distal cues surrounding the arena, and the remainder of the recording room. Afterwards, for 10 to 40 min, each animal foraged for scattered food crumbs (corn puffs) in a square open-field (OF) box with a floor size of 150 × 150 cm or 200 × 200 cm and a height of 50 cm. The arena had dark blue or black wax/vinyl flooring. The box was placed on the floor centrally in a large room (16 or 21 m^2^) with full visual access to background cues. The length of the foraging session was determined by how quickly the rat explored the entire open field arena. Sessions were not contiguous in all rats. If the open field arena was not covered completely during any session, the rat was returned to its home cage for several hours.

Following surgery, data were obtained from 2 to 21 recording sessions per animal. Most recordings were performed within the first week after surgery, with the first recordings performed when normal behavior was resumed a minimum of 12 hours after the completion of surgery (full range of recordings: 0-30 days post-operatively; median: 4 days).

Three different protocols were used to induce remapping in the hippocampus (Supplementary Table 1). In the first protocol (‘different-rooms’, defined as Room A and Room B; Figures 1-4), the rats were familiarized with open field arenas in two separate rooms with different distal cues^3^. In the second protocol (‘novel-room’; Figures 3-4), the rats were familiarized with the open field arena and distal cue configuration in one room (familiar room, F) and introduced to a novel room with an open field arena and a unique distal cue configuration during recording (novel room, N). In the third protocol (‘double rotation’; Figures 3-4), rats were familiarized with a standard (S) configuration of local and distal cues in one room.

During recording, the rat was introduced to a rotated (R) configuration of those same cues for the first time. Local cues were rotated by 90° clockwise and distal cues were rotated by 90° counterclockwise, resulting in a 180° mismatch between these sets of cues. This type of double rotation paradigm has been shown previously to induce partial remapping on circular and plus-shaped mazes^16–18,68^.

All animals participated in more than one experiment. Only one experiment per animal is included in Figure 1. When experiments were repeated with roughly the same ensembles of cells in the same set of recording rooms, only the recording with the largest number of grid modules or the largest number of grid cells per module, was analyzed (Supplementary Data Table 1). In Figure 2, only one experiment is included per animal, except in one animal (#28367) that was familiarized with three rooms prior to recording and therefore participated in two ‘different-rooms’ recordings involving exploration of familiar rooms (ABBA and ACCA).

### Different-rooms task

In the different-rooms task, the rats explored a 150 cm^2^ or 200 cm^2^ square black open field arena in two rooms (A and B, see above) with prominent distal cues that differed between the rooms. Before the start of each session, the rat was placed on a 1 m-high pedestal in order to connect the recording cables. During this time, the rat was able to inspect the distal cues in the recording room. Once connected, the rat was picked up and gently placed in the open field arena. The rat was tested first in room A (session A1) and then in room B (session B1). Before the second session in each room (A2 and B2), the floor of the open field was cleaned. The minimum intertrial duration was 5 min. In Figure 2, we only included data from rats exposed to the ABBA sequence. One additional rat (#29731) was exposed to a different room sequence as part of a separate study. Sessions in rooms A and B from this recording are included in Figs. 3 and 4, which display data from the full range of experimental conditions.

### Novel-room task

In the novel-room task, the rats explored a familiar 150 cm^2^ square black open field arena in a room with prominent distal cues (familiar session, F1). Following the first session, the rat was moved to an unfamiliar room (novel session, N1). The sequence of recording was FNNF (unless the open field arena was not covered completely within a session). All other procedures were conducted as in the different-room task with familiar arenas.

### Double-rotation task

In the double-rotation task, the rats explored a 150 cm^2^ or 200 cm^2^ square black open field arena with a standard (S) configuration of local and distal cues. The local cues consisted of four quadrants of textured flooring that were equal in area: (1) blue wax/vinyl, (2) gray stripes, (3) black hashmarks, (4) gray dots. One white paper cue was also placed on each wall of the open field halfway between the corners of the arena as additional local cues. The following four shapes of paper cues were used: (1) circle, (2) triangle, (3) square, and (4) diamond. The distal cues were large objects placed just outside the walls of the open field arena in clear view of the rat. Each of the following distal cues was placed outside the perimeter of the arena, halfway between the corners: (1) black-and-white striped card (30 cm wide x 1 m high), (2) white card (30 cm wide x 1 m high), (3) circular tube (10 cm diameter x 1 m high), (4) cardboard box affixed to broom handle (15 cm wide x 1 m high). A desk lamp on top of a pedestal was placed at one corner of the open field and it was rotated along with the distal cues. The open field was surrounded by dark curtains. When tossing food crumbs into the open field, the experimenter briefly opened the curtains at one corner of the open field. The location of the corner used to toss food crumbs was also rotated along with the distal cues.

Before the start of the first session, the rat was placed on a 1 m-high pedestal in order to connect the recording cables. During this time, the rat was able to inspect the distal cues of the recording room that were outside the curtains. Once connected, the rat was picked up and gently placed in the open field area to explore the standard configuration of cues (session S1). After this session, the rat was placed back on the pedestal and the recording cables were disconnected. The rat was placed in a holding chamber on the floor of the room (30 × 30 cm floor, 80 cm height) so that the remainder of the recording room was not visible. The local cues were then rotated by 90 degrees CW and the distal cues were rotated by 90 degrees CCW, resulting in a 180° mismatch between the sets of cues. The rat was then placed back onto the pedestal and the recording cables were connected. The rat subsequently explored the rotated configuration of cues for the first time (session R1). Before a second session in each configuration (sessions R2 and S2), the floor of the open field was cleaned. The rat was then tested for a second time in the rotated configuration and in the standard configuration (unless the arena was not covered completely within any session). The minimum intertrial duration was 5 min.

### Spike sorting and unit selection

Spike sorting was performed with KiloSort 2.5 (Ref. 35) with customizations as described previously^8^. Units were discarded if more than 1% of their interspike interval distribution consisted of intervals less than 2 ms. Units were also excluded if their mean spike rate was less than 0.05 Hz or greater than 7 Hz across the full recording duration, or if they were recorded on sites outside the region of interest. One recording was excluded from further analysis because drift traces exceeded 15 µm (i.e., the spacing between adjacent probe contacts).

### General electrophysiological analysis

In order to exclude spiking activity occurring during periods of immobility, a walk filter (5 cm/s) was applied. Rate maps for each unit were generated for each recording session by binning the location of each spike for each session, dividing the number of spikes in each bin by the time spent in that bin, and smoothing with a Gaussian kernel (σ = 2 spatial bins). For 150 × 150 cm^2^ environments, 3.75 cm^2^ bins were used. For 200 × 200 cm^2^ environments, 5 cm^2^ bins were used.

Mean firing rate was defined as the total number of spikes divided by the duration of the recording session. Peak firing rate was defined as the maximal firing rate of all spatial bins. To assess spatial correlation, pairs of rate maps were reshaped into a single vector and the correlation coefficient (Pearson correlation) between these vectors was calculated. Pixels of incongruity between the two vectors, resulting from unvisited pixels in either epoch, were excluded from the calculation. Difference-over-sum scores were calculated as:

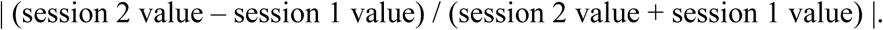

The spatial information rate^69^ of each unit was computed as:

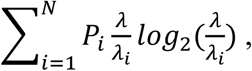

where *N* is the total number of bins, *λ_i_* is the unit’s mean firing rate in the *i*-th bin of the rate map, *λ* is the overall mean firing rate, and *p_i_* is the probability of the animal being in the *i*-th bin (time spent in the *i*-th bin divided by the duration of the session). Spatial information content (bits/spike) was obtained for each session by dividing the spatial information rate by the mean firing rate of the cell in that trial.

### Identification of grid cells and grid modules

To identify grid cells and split them into modules, we used a previously described method^8,70^. In brief, this method uses a nonlinear dimensionality reduction algorithm (UMAP^50^) and a clustering algorithm (DBSCAN; MATLAB) to find clusters of cells that express similar patterns of spatially periodic activity without making any assumptions about the geometry of the grid pattern. 2D autocorrelograms were calculated from coarse-grained spatial rate maps of each cell (10 × 10 cm bins; no smoothing). Autocorrelogram bins within a central radius of 2 bins or beyond an outer radius equal to the size of the rate map were masked, before vectorizing and concatenating the autocorrelograms in a matrix. Each point has an N-dimensional position in the resulting point could. The Manhattan distances between all points were calculated and the nearest neighbors were identified for each point. We used the following hyperparameters for UMAP and DBSCAN: UMAP, ‘metric’ = ‘manhattan’; ‘n_neighbors’ = 4-10; ‘min dist’ = 0.05; ‘init’ = ‘spectral’; DBSCAN, ‘minimum points’ = 4-10; eta 0.3-0.6. The DBSCAN algorithm was used to partition the point cloud into clusters.

We computed the autocorrelogram of all units in each cluster to obtain the spacing, orientation, and grid score of each identified cluster (Extended Data Fig. 2a). Grid pattern consistency^70^ was measured by computing the Pearson correlation between the average autocorrelogram of the cluster and each individual cell’s autocorrelogram. The median consistency across all cells in the cluster was defined as the cluster’s grid consistency. For a cluster to be classified as grid module, three criteria had to be fulfilled: (1) a clear grid pattern in the autocorrelogram of the cells in the cluster, (2) a grid score greater than zero, and (3) grid pattern consistency greater than 0.40. Of the 142 identified clusters, 98 were classified as grid modules. 44 were excluded from further analysis (i.e., Cluster 4 from #27150, shown in Extended Data Fig. 2a). Grid scores were significantly higher for included versus excluded clusters (included clusters: median = 1.26, 95% CI, 1.16 - 1.35; excluded clusters: median = - 0.08, 95% CI, −0.16 - −0.01).

We assessed the quality of the clustering results by calculating the silhouette value for each cluster, which measures how well each data point fits within its assigned cluster compared to other clusters (Extended Data Fig. 2b). In rare situations, an additional module of grid cells was evident that was not detected by UMAP (due to low numbers of units or large spacing of the grid pattern). In these cases, the module was identified using the depth, spacing, and spatial information of the recorded units.

### Grid scale and grid orientation

For each cell in a cluster identified as a population of grid cells, a spatial autocorrelogram was calculated for the smoothed rate map from each session^71^. The six fields closest to the center of the autocorrelogram (‘the inner ring’) were used to estimate the scale and orientation of the cell. Grid spacing was calculated as the arithmetic mean of the distance of each field relative to the origin. We verified our estimation of grid spacing using a second method, previously described^72^ to isolate individual grid fields without setting an arbitrary threshold for local peak detection. In brief, we transformed autocorrelograms into a greyscale image of connected regions by applying the morphological image processing operations erosion and reconstruction (MATLAB image processing toolbox).

Individual grid fields were defined as areas of at least 160 cm^2^ where the firing rate exceeded 30% of the peak rate. To optimize grid field detection, grid fields with peak rates lower than 1 Hz were ignored. Individual grid field rates were defined as the peak rate of each identified firing field. Changes in grid field rates were assessed using absolute difference scores (Extended Data Fig. 9d).

### Grid ellipticity

To measure how much the grid pattern was distorted between sessions, we calculated ellipticity (ε) for each grid module^39^ as follows:

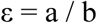

where *a* is the semi-major axis and *b* is the semi-minor ellipse axis. Ellipticity ranges from 1 to ∞, where 1 represents a perfect circle (length of semi-major and semi-minor axes are equal) and ∞ corresponds to the limit at which the ellipse is parabolic.

### Spatial crosscorrelograms

To determine how the grid pattern rotated and/or shifted (i.e., changed phase) between rooms, we computed spatial crosscorrelograms for each module, following previously published methods^3^ (Fig. 1b; Extended Data Fig. 3a). These ‘*module crosscorrelograms*’ were generated by stacking the two-dimensional rate maps of grid cells within the module into a three-dimensional matrix, with cell identity along the z-axis. Two such stacks were constructed, one for each session (e.g. A1 and B1), with cells arranged in the same order. To assess rotational alignment, one of the stacks was rotated in 3° increments from 0° to 360°. At each rotation, we calculated the crosscorrelation dot product between population vectors of activity in corresponding bins. We then shifted one stack relative to the other in steps of 3.75 cm across both the *x* and *y* axes of the environment. These shifts produced a two-dimensional spatial crosscorrelogram: for the 150 × 150 cm^2^ environment, the result was an 80 × 80 matrix of 3.75 cm bins (covering 300 m^2^), and for the 200 × 200 cm^2^ environment, a 80 × 80 matrix of 5.0 cm bins (covering 400 cm^2^). The crosscorrelogram obtained by crosscorrelating two grid patterns was itself a grid pattern. Each bin in this crosscorrelogram contained a correlation value (range −1 to 1; color-coded in the figures), normalized by the number of contributing pixels (number of pixels included for that bin / total number of pixels). From each spatial crosscorrelogram (one per rotation angle), we extracted the peak correlation value. The rotation of each module was defined as the angle at which the maximum correlation occurred. Note that all recording rooms shared the same reference frame (see section for Behavioral tracking). Crosscorrelograms were also computed for individual cells, as in Fig. 1a, although all subsequent analyses were based on population-level (module) data.

Having determined the rotation of each module, we next examined how the phase of the grid pattern changed between sessions (Extended Data Fig. 3). If the grid phase remained stable relative to external boundaries (e.g., the walls of the arena), the central peak of the crosscorrelogram would be at the origin of the crosscorrelogram (as in Fig. 1c, left). In contrast, a shift in grid phase would result in displacement of the central peak away from the origin (as in Fig. 1c, right). Because grid cells within each module shifted coherently, the crosscorrelogram retained its regular grid pattern, but the location with the peak correlation shifted to a new location. By quantifying the displacement of this peak from the origin, we were able to estimate the shift – or phase change – of the grid pattern between sessions, as described in detail below.

We re-computed spatial crosscorrelograms for each module using four additional sizes of spatial bins (20 x 20 matrix of 7.5 cm bins; 30 x 30 matrix of 5 cm bins; 50 x 50 matrix of 3 cm bins; 60 x 60 matrix of 2.5 cm bins; Extended Data Fig. 6f) and two additional bin sizes for rotation (1° and 2° bins; Extended Data Fig. 6g).

### Unit hexagons

Because of the grid’s periodicity, shifts in grid phase produce crosscorrelograms with spatially periodic peaks, making it non-trivial to determine which peak should serve as the reference for estimating displacement (Extended Data Fig. 3a). To handle this challenge, we transformed each crosscorrelogram into a repeating series of hexagonal tiles spanning the entire crosscorrelogram (Extended Data Fig. 3b). We first computed the spatial autocorrelation of the module’s spatial crosscorrelogram at the rotation with the highest correlation (Extended Data Fig. 3b2). From this autocorrelogram, we identified the six peaks surrounding the central peak to determine the grid’s spacing and orientation (Extended Data Fig. 3b3). The orientation of each of the three grid axes (A_X_N) was used to define the corresponding axes (H_A_N) of a central hexagon. For example, if Ax1 = −60°, Ax2 = 0°, and Ax3 = 60°, then H_A_1 = −30°, H_A_2 = 30°, and H_A_3 = 90°. The spacing of the grid pattern was used to calculate the circumradius of the hexagon (*h*; Extended Data Fig. 3b4):

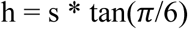

The six vertices surrounding the center (*x_N_,y_N_*) were then computed using:

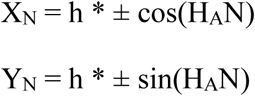

These vertices defined the perimeter of the central unit hexagon (Extended Data Fig. 3b4). The unit hexagon was then overlaid onto the module crosscorrelogram. Identical hexagons, each with the same radius and orientation, were then tiled across the entire crosscorrelogram (Extended Data Fig. 3b5). This ensured that one peak of the crosscorrelogram fell within each tile, with identical relative positions across tiles (assuming regularity of the grid pattern). The displacement of each module was then represented as a single point within the central hexagon (centered at the origin), and its magnitude was calculated as the Euclidean distance (in cm) from the origin to the peak within that hexagon (Extended Data Fig. 3c). Because of the hexagon’s symmetry, points on one edge are equivalent to points on the opposite edge and can be ‘wrapped’ around any of its three axes (Extended Data Fig. 3d).

To compare the displacement of modules with different scales, we computed normalized displacement: the Euclidean distance (in cm) from the origin to the peak within the central hexagon, divided by the modulés grid spacing (Extended Data Fig. 3c). The maximum possible normalized displacement (*d_max_*) for any module is:

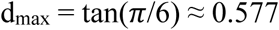

To determine whether grid cells within each module shifted coherently between sessions, we generated crosscorrelograms for each grid cell (Fig.1a). We then calculated the distance (in cm) between the peak within the central unit hexagon and peak within the central unit hexagon for the module to which the cell belonged.

We then tested whether phase changes of intramodular grid cell pairs were more similar than phase changes of transmodular grid cell pairs, as expected if modules aligned differentially. Similarly, to determine whether grid cells within each module rotated coherently between sessions, we calculated the angular difference between the rotation of the module and the rotation of each individual grid cell in that module. We tested whether the rotation of intramodular grid cell pairs was more similar than the rotation of transmodular grid cell pairs.

To determine whether modules shifted coherently between sessions, we compared the phase change of each module between pairs of sessions (Extended Data Fig. 3e). To do this, we calculated the distance between the location of the peak within the central hexagonal tile of the crosscorrelogram for each pair of simultaneously recorded grid modules. For all module pairs, we determined which peak within the central tile of each module were nearest to one another (i.e., had the smallest distance), taking the equivalence of points at the edges of the hexagon into account by wrapping around the edges of the hexagon. We normalized these pairwise distance values to the maximum possible distance between modules to allow for direct comparison of modules with different scales. The maximum possible distance between modules is equal to the spacing of the larger module multiplied by tan(π/6).

### Estimation of phase changes expected by chance

For each animal, we calculated a shuffled distribution in order to obtain an estimate of how different the phase changes of a set of three or more grid modules would be by chance. To generate the distribution, we shifted the grid patterns of the different modules randomly with respect to one another a total of 1,000 times and calculated the displacement from the origin as well as the pairwise distance between modules in the same manner as described for the experimental data above, repeating the procedure 1,000 times for each rat.

### Alternative methods for calculation of pairwise distance between modules

In the Main Figures, the pairwise distance between the endpoints of the phase-shift vectors of different modules was calculated using the spatial crosscorrelogram at the rotation with the highest correlation. We repeated the analysis at the following rotations: (1) the mean rotation of all modules, (2) the rotation of module M1, (3) the rotation of M2, and (3) the rotation of M3.

To take into account any influences of distorted grid patterns^53,73–75^, which might change phase distances differently across the crosscorrelogram, we also tested whether there was any location in the entire crosscorrelogram (beyond the unit hexagon) where the peaks of each module were closer together, indicating that they shifted more coherently (Extended Data Fig. 5d1). This method considers each peak in the crosscorrelogram as a unique location rather than a periodically repeating one, and therefore allows grid modules to shift beyond their maximum displacement (d_max_). When this method was used, the hexagonal tiling of each module was aligned at the center and overlaid to cover the entire correlogram. We then searched for the location, anywhere in the correlogram, where the mean pairwise distance between modules was the lowest, which we refer to as the location with the minimum distance (Extended Data 5c1).

In a second approach, we computed a summed crosscorrelogram for each experiment by adding the spatial crosscorrelograms of all simultaneously recorded modules together at the rotation with the highest correlation to yield one crosscorrelogram for *all* modules (Extended Data Fig. 5d2). Module crosscorrelograms were rescaled from zero to one prior to summation in order to control for the number of cells per module. We identified the location with the highest correlation in the summed crosscorrelogram (referred to as the summed peak), which served as an estimate for the phase change of each module. We then determined which regional maximum in each module crosscorrelogram was closest to the summed peak. The phase change for each module was thus represented by a vector that pointed from the origin to this regional maximum. We then calculated the distance between the endpoints of these phase change vectors to determine whether modules shifted coherently between sessions.

### Varying the selection criteria for grid modules

We computed spatial crosscorrelograms for each grid module using more conservative or more liberal cell selection criteria to show that our measurements of module phase change and rotation between sessions are robust to noise and contamination. We refer to the modules of grid cells shown in Supplementary Table 1 (identified using the criteria described above) as the ‘original’ modules. More conservative selection criteria reduced the number of grid cells per module, while more liberal selection criteria increased the number of cells in each module. To be more conservative, we: (1) used even stricter criteria for inclusion of single units than those described above (Extended Data Fig. 6c1), or (2) used only 25 grid cells with spacing and anatomical depth values closest to the mean depth of the module (Extended Data Fig. 6c2). To be more liberal, we: (3) included additional cells with spatial information above the 95^th^ percentile of a shuffled distribution and anatomical depth that was closest to the mean depth of the module (Extended Data Fig. 6c3), or (4) included additional cells with spatial information below the 95^th^ percentile of a shuffled distribution and depth that was closest to the mean depth of the module (Extended Data Fig. 6c4). When adding neighboring cells with spatial information content above the shuffled criterion (3), an increasing percentage of the original module was added in each run (30%, 50% or 70% of the total number of grid cells in the module was added to the original module). When adding neighboring cells with spatial information content below the shuffled criterion (4), an increasing percentage of the original module was added in each run (10%, 20% or 30% of the total number of grid cells in the module was added to the original module).

To determine if our results were robust to contamination from neighboring modules, we included additional grid cells from anatomically adjacent modules (Extended Data Fig. 6c5). Four runs were completed: (1) grid cells from M2 were added to M1; (2) grid cells from M1 were added to M2; (3) grid cells from M3 were added to M2; (4) grid cells from M2 were added to M3. To maximize the effect of contamination, we added grid cells with spacing and anatomical depth values that were closest to the mean spacing and anatomical depth of the original module. An increasing percentage of the original module was added in each run (10%, 30% or 50% of the total number of grid cells in the module was added to the original module).

### Place cell classification

Place cells were defined as putative excitatory neurons (mean firing rate < 7 Hz) with high spatial stability (spatial correlation between first and second halves of session > 0.5), high spatial information content, peak rate > 1 Hz, and at least one identified place field in at least one session. Place fields were defined as areas with at least 8 contiguous 5 cm^2^ pixels (200 cm^2^) where the firing rate exceeded 30% of the peak rate.

The shift in place field location was determined using rate maps for each cell and was measured as the distance (cm) between the *x,y* location of the peak firing rate for each place cell in each session. Each place cell was required to have an identified field in both sessions, and we computed the change in place field location between fields with the peak firing rate in each session, regardless of the number of identified place fields.

To compute population vectors for the entire ensemble of simultaneously recorded place cells between sessions, rate vectors were constructed by arranging the rate maps for that session into a three-dimensional matrix, where the two spatial dimensions are represented on the x and y axes and cell identity on the z axis. The distribution of mean rates along the z axis for a given x-y location represents the composite population vector for that location. The correlation coefficient was computed for each pair of population vectors corresponding to the same location in the two conditions. The central tendency of the distribution provides a measure of similarity of the hippocampal ensemble activity at corresponding locations in the two conditions.

Spatial crosscorrelograms for ensembles of simultaneously recorded place cells were computed in the same manner as described for grid cell modules above. The strength of the resulting population vector correlation gives an indication of whether simultaneously recorded place cells underwent a coherent rotation or translation between sessions. If place cells remained stable relative to external boundaries (e.g., the walls of the arena) between sessions, the central peak of the crosscorrelogram would remain at the origin of the crosscorrelogram (as in Fig. 1c, left). If place cells rotated coherently between sessions, the central peak would also remain at the origin, and the rotation with the highest PV correlation would indicate the rotation of the ensemble between sessions. If place fields shifted coherently between sessions, the central peak would be displaced from the origin. Finally, if place fields rotated and/or shifted randomly with respect to one another, there would no clear peak in the crosscorrelogram at any rotation.

To quantify the degree of place field reorganization within each experiment at the population level, we used a rearrangement score^21^, which measures the degree of correlation between pairwise place field distances in sessions A and B. For session A and session B, we generated vectors (D_A_, D_B_) containing the Euclidean distance (cm) between the location of peak firing for all pairs of place cells: D_A_ = [d_1,2_; d_1,3_; d_2,3_] and D_B_ = [d_1,2_; d_1,3_; d_2,3_]. The rearrangement score is equal to one minus the Pearson correlation between these two vectors (1 – corr[D_A_,D_B_]). Values near zero indicate that the distances between all pairs of place fields remained unchanged between sessions, either because place fields did not change location or because all place fields moved coherently. Values near 1 indicate that the distances between many place field pairs changed between sessions, reflecting a random reorganization of the place cell code between sessions.

### Classification of grid, head direction (HD), and border cells

To classify HD and border cells, chance levels were estimated using a shuffling procedure^76^. For each cell, its spikes were circularly shifted in time relative to the rat’s position by a random amount between 20 s and 20 s less than the total length of the recorded session. The mean vector length or border score was then calculated using these shuffled spike times, and this procedure was repeated 1,000 times for each cell. A distribution of values was generated from these 1,000 repetitions, and the 95^th^ and/or 99^th^ percentile of that distribution was calculated.

Firing fields were defined as areas with at least 8 contiguous 5 cm^2^ pixels (area > 200 cm^2^) where the firing rate exceeded 30% of the peak rate. Fields with peak rate was less than 1 Hz were ignored. To optimize grid field detection (for the analysis of grid field firing rate change in Extended Data Fig. 9d), individual grid fields were defined as areas of at least 160 cm^2^. Individual grid field rates were defined as the peak rate of each identified firing field. Changes in grid field rates were assessed using absolute difference scores (Extended Data Fig. 9d).

Spatial autocorrelations and grid scores were calculated as described previously^71^, based on the individual cells’ rate maps. Briefly, the grid score was determined for each cell by rotating the autocorrelation map for each cell in steps of 6° and computing the correlation between the rotated map and the original. The correlation was confined to the area defined by a circle around the outermost peak of the six peaks closest to the center of the spatial autocorrelation map. If fewer than 6 peaks were identified, the circle was fitted around the outermost peak. The central peak was not included in the analysis. The grid score was computed as the difference between the lowest correlation at 60° and 120°, and the highest correlation at 30°, 90°, and 150°.

Head direction tuning maps were created by plotting firing rate as a function of the rat’s directional heading. Tuning curves were divided into 6° bins. The length of the mean resultant vector (mean vector length, MVL) was calculated from the head direction tuning curve, as described previously^71^. We also examined the intra-trial tuning stability (Pearson correlation between head direction tuning curves in first and second halves of session). Head direction cells were defined as putative excitatory neurons (mean firing rate < 10 Hz) with mean vector length and intra-trial tuning stability that exceeded the 99^th^ percentile of the shuffled distribution of each measure. When comparing two sessions, a cell had to meet these criteria in both sessions. Head direction cells from two recordings were excluded (#26823 and #28739) due to inaccurate angular, but not positional, tracking data from Motive.

Border scores were calculated by dividing the difference between the maximal length of a wall touching any defined firing field and average distance of that field from the nearest wall by the sum of the same two values^77^. Thus, border scores ranged from −1 to 1; a score of 1 indicates firing exclusively along the entire length of a wall. Border cells were defined as putative excitatory neurons (mean firing rate < 10 Hz) with high spatial stability (spatial correlation between first and second halves of session > 0.5), border scores above the 95^th^ percentile for that cell, and at least one identified field. When comparing two sessions, a cell had to meet these criteria in both sessions.

### Model of grid-to-place cell transformation

To investigate the impact of differential translation and/or rotation of grid modules on hippocampal place cells, we adapted a previously published a competitive linear summation model of the grid-to-place cell transformation by separating the grid cell inputs into three modules with realistic spacing values^54^. In brief, this model is a three-layer network where place fields are created by linear summation of weighted inputs from entorhinal grid cells and dentate gyrus granule cells. Competitive interactions limit the active pool of neurons to only those receiving the most excitation (10% winner-take-all process), which is meant to mimic gamma frequency feedback inhibition.

We generated a library of 10,000 simulated grid cells split into three modules with spacing values closely matched to our empirical data (M1 spacing = 50 cm, M2 spacing = 70 cm, M3 spacing = 100 cm). The number of simulated grid cells in each module was adjusted to match the anatomical distribution of inputs to the hippocampus (i.e., highest density of projections to dorsal hippocampus from dorsal MEC^78–80^; M1 = 5,500 cells, M2 = 3,000 cells; M3 = 1,500 cells). All grid cells had the same orientation and uniform field rates.

Each simulation was run with the following parameters: arena size = 100 × 100 cm, bin size = 1 cm, number of grid cells = 10,000, number of granule cells = 200, excitation (E) = 10%, relative DG-to-MEC input to CA3 (R) = 0.24, place field rate threshold = 20%, minimum place field size = 200 cm^2^. E refers to the ‘E%-max’ rule, in which cells fire if their feedforward excitation is within *E*% of the cell receiving maximal excitation.

The original library of 10,000 simulated grid cells served as input to a set of simulated place cells (Session A). Note that all connections and weights were held constant from Session A to Session B. The only variable that was changed between sessions was the rotation or phase of each grid module.

For simulations of module translation, the possible *x,y* locations to which the phase of each module could be shifted were equally spaced throughout the unit hexagon of that module in steps of 4 (M1, 120 locations; M2, 248 locations; M3, 504 locations). In total, this resulted in 1.5 × 10^7^ combinations. We selected 12,300 combinations for analysis by systematically varying the mean distance between modules and their displacement from the origin. For simulations of module rotation, the orientation of each module was varied from 0° to 60° in 3° steps, resulting in 21 possibilities for each module. In total, this resulted in 9,261 (21^3^) possible combinations. The grid pattern for all grid cells in each module was shifted or rotated coherently. Place cell rate maps were then created using these shifted or rotated grid cells (Session B).

To quantify the effect of module rotation or translation on the simulated population of place cells, we used the rearrangement score^21^ (full description included above). Values near zero indicate that the distances between all pairs of place fields remained unchanged between sessions, either because place fields did not change location or because all place fields moved coherently. Values near 1 indicate that the distances between many place field pairs changed between sessions, reflecting a random reorganization of the place cell code between sessions.

### Histology and recording locations

When recordings were complete, rats were deeply anesthetized with isoflurane, given an overdose of sodium pentobarbital (Euthasol, 100 mg/ml), and perfused intracardially with 0.9% saline followed by 4% formaldehyde. Extracted brains were stored in formaldehyde at room temperature until sectioning. Brains were sectioned on a cryostat at 30 µm and mounted on glass microscope slides. Tissue was sliced in sagittal sections for MEC. For multi-shank probes implanted orthogonal to the midline in the hippocampus, tissue was sliced in coronal sections. For single-shank probes implanted in the hippocampus and for multi-shank probes implanted parallel to the midline in the hippocampus, tissue was sliced sagittal sections. For identification of recording sites, the tissue was Nissl stained with cresyl violet. Photomicrographs were taken through a Zeiss Axio Imager. For identification of recording sites, we used two reference points: (1) the ventral boundary was estimated on the section where the tip of the shank was visible; and (2) the dorsal boundary was estimated on the section where the shank exited the surface of the brain.

### Data analysis and statistics

Data analyses and statistical analyses were performed using custom-written scripts in MATLAB 2020a. Samples included all available cells that matched the classification criteria for the relevant cell type. Power analysis was not used to determine sample sizes. The study did not involve any experimental subject groups; therefore, random allocation and experimenter blinding did not apply and were not performed. Assumptions of parametric tests (i.e., normality, homogeneity of variance) were formally tested. When assumptions of parametric tests were not violated, the error is reported as the standard error of the mean. When assumptions of parametric tests were violated, we used nonparametric tests and report the 95% confidence interval. Exact *p* values were provided unless the computed *p* value was below the numerical limit of machine precision, in which case we report *p* < 2.2 × 10^−16^. When relevant, we also report Cohen’s d as a measure of effect size. ***p < 0.001, **p < 0.01, *p < 0.05.

---

**Extended Data Fig. 1.**
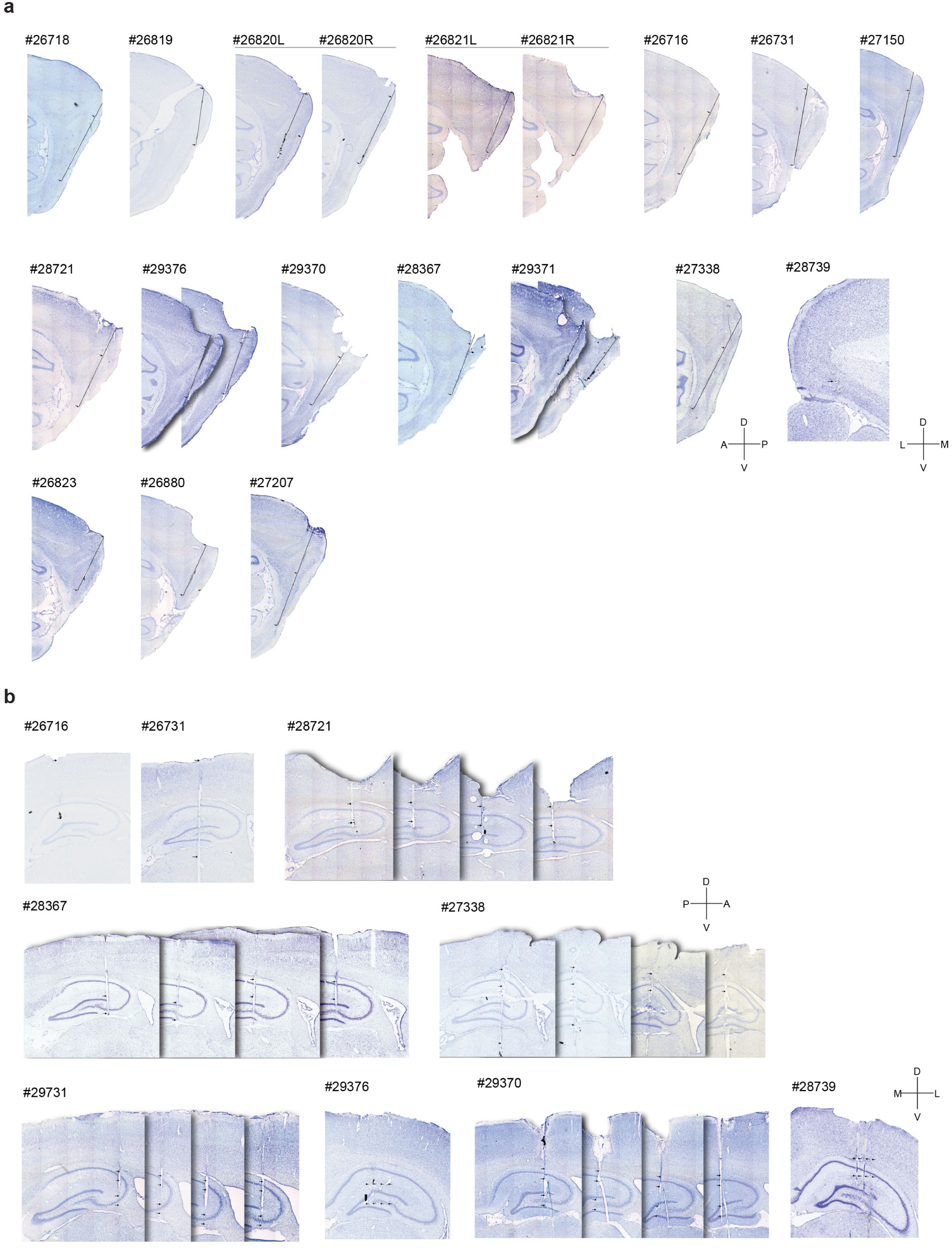
Nissl-stained brain sections showing recording locations in MEC/PaS and hippocampus. **a**, Brain tissue stained with cresyl violet shows probe placement in MEC/PaS in 14 rats (ID noted at top; see Supplementary Table 1). Probes were mostly confined to layers II and III. Sagittal sections are shown except for #28739, which was sectioned coronally. D = dorsal, V = ventral, M = medial, L = lateral, A = anterior, P = posterior. Note that an extended portion of the MEC/PaS was covered by each probe. **b**, Brain tissue stained with cresyl violet shows probe placement in the hippocampus of 9 rats (ID noted at top; see Supplementary Table 1). Sagittal sections are shown except for #29376, #29370, #28739, and #29731, which were sectioned coronally. Histology is not shown for one animal (#27150) whose brain tissue was damaged during processing.

**Extended Data Fig. 2.**
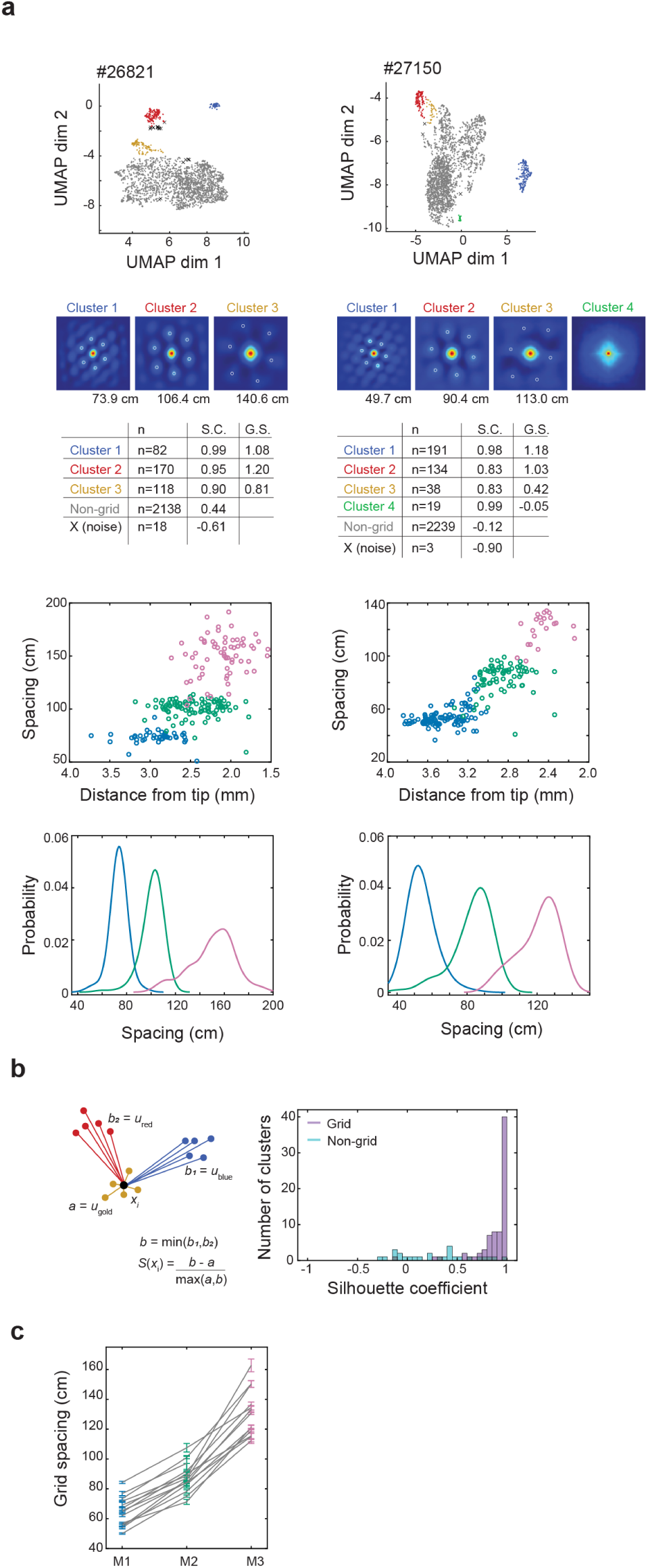
Classification of grid modules. **a**, First row, procedure for classifying grid cells into modules shown for two representative experiments. For each experiment, scatterplots show the two-dimensional UMAP projection of autocorrelograms of all simultaneously recorded units. Each point is colored according to its DBSCAN cluster assignment. Animal ID is noted above each plot. Second row, panels show the autocorrelogram of all units in each cluster during session A1. The grid spacing (cm) calculated from the autocorrelogram of the cluster is noted below each panel. Each table shows the number of units (n), silhouette coefficient (S.C.), and grid score (G.S.) calculated from the autocorrelogram of rate maps from session A1 for each cluster. Third row, points show, for each grid cell, the estimated anatomical depth (distance from the tip of the Neuropixels probe) versus the cell’s grid spacing for experiments in rats #26821 and #27150. Points are colored according to their module assignment (M1 = blue, M2 = green, M3 = pink). Fourth row, kernel smoothed density (KSD) estimate of grid spacing for cells from rats #26821 and #27150. **b**, Left, schematic shows how the silhouette value of each cluster is calculated. The silhouette value of each cluster (x_i_) is given by: S(x_i_) = (b – a) / max(a,b), where a is the mean distance of x_i_ to each point in the cluster, and b is the minimum of the mean distances of x_i_ to points in other clusters. Right, histogram shows silhouette coefficient for all grid (purple) versus non-grid (green) clusters identified using UMAP/DBSCAN (98 grid clusters, 14 rats). **c,** Colored lines show grid spacing (cm) for each module in n = 14 rats shown in Supplementary Table 1. Error bars indicate S.E.M.

**Extended Data Fig. 3.**
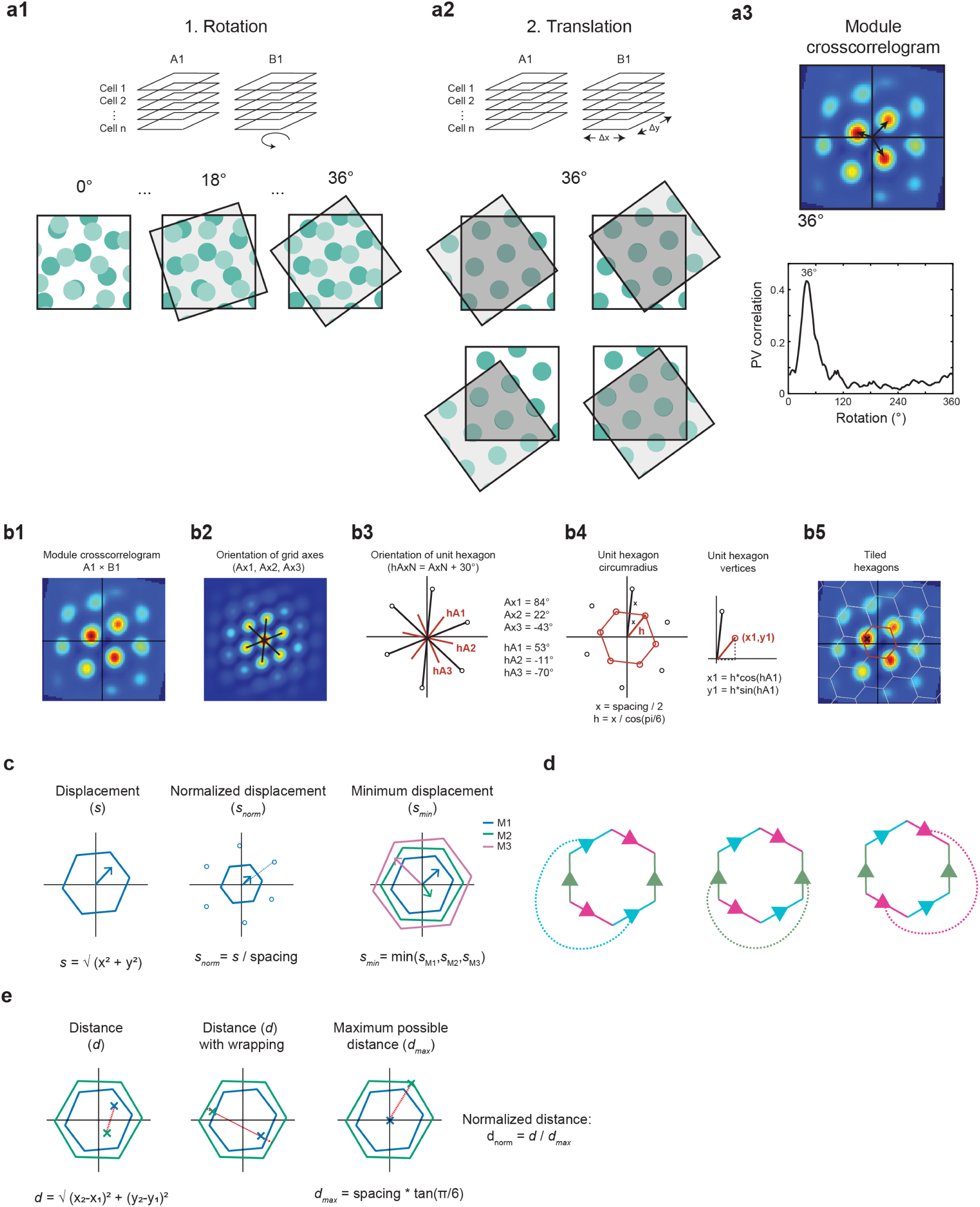
Procedures for calculating rotation and displacement of grid maps. **a1**, Illustration of procedure to determine grid rotation between sessions in different rooms. Schematic illustrates procedure for crosscorrelation of rate maps for ensembles of grid cells from the same module. To compare two recording sessions, one stack of rate maps was rotated counterclockwise in 3° steps from 0° to 360° and the rotation that gave the highest correlation with the original map was selected. **a2**, Illustration of subsequent procedure to determine grid translation between sessions in different rooms. At each rotation, we calculated a crosscorrelation matrix by shifting one stack of maps in 3.75 cm steps along the entire x and y axis of the environment. Dark gray shading depicts overlapping pixels within rate maps of each session. These pixels were included in the calculation of the population vector (PV) crosscorrelation. **a3**, Top, within the crosscorrelogram at each rotation, we identified the maximum correlation. Black arrows illustrate how one stack of maps was shifted with respect to the second stack in order to align (as much as possible) the grid pattern in the rate map from each session (i.e., to the point that gives the highest correlation between the rate maps). Note that the crosscorrelogram of two grid patterns is itself a grid pattern. We refer to the distribution of correlation values across the xy space at the rotation with the highest correlation as the module crosscorrelogram. Bottom, the rotation for each module was defined as the rotation at which the highest correlation from all population crosscorrelograms was found. **b1-b5**, Schematic of procedure used to determine the unit hexagon for each module and to calculate the phase change of the module between sessions. **b1**, Module crosscorrelogram between sessions in different rooms (A1×B1). **b2**, We first calculated the autocorrelation of module crosscorrelogram at the rotation with the highest correlation to find the orientation of each grid axis (Ax_1_, Ax_2_, Ax_3_). **b3**, Orientation of each axis of the unit hexagon (hA_1_, hA_2_, hA_3_) was calculated using the axes of each module (hA_n_ = Ax_n_ + 30°). **b4**, Module spacing (calculated from autocorrelogram in **b2**) was used to calculate the circumradius of the unit hexagon (h) as given by the following equation: h = (spacing/2) * cos(π/6). Each vertex of the unit hexagon was calculated using the circumradius of the unit hexagon and the orientation of its axes as given by the following equations: x_n_ = h * cos(hA_n_) and y_n_ = h * sin(hA_n_). **b5**, Phase change of each module between sessions is defined as the location of the peak (black cross) within the central unit hexagon (red outline). Phase change on all unit tiles is identical when the grid has no local distortions. **c**, Left, phase change (i.e., displacement) of each module between sessions (*s*) was defined as the distance (in cm) from the origin to the location with the highest correlation within the central unit hexagon of the module crosscorrelogram. Middle, the normalized displacement for each module (*s*_norm_) is the displacement divided by the spacing of the module. Right, the minimum displacement (*s*_min_) was defined as the shortest displacement among simultaneously recorded modules. In the example shown, the minimum displacement is that of M2 (green arrow). **d**, Dashed lines depict the three axes around which points on the hexagon can be wrapped for calculations of distance (Ax1, blue; Ax2, green, Ax3, pink). Points that fall on the edges of the hexagon are identical to points on the corresponding edge of the hexagon after wrapping. **e**, Left, for each pair of simultaneously recorded grid modules, we calculated the Euclidean distance (*d*) between the endpoints of the vectors that represented the change in phase for each module (i.e., pairwise distance between module phase changes). Pairwise distance between phase changes for M1 (blue) and M2 (green) is shown in red. Middle, example of how distance is calculated between points representing the phase change of M1 (blue) and M2 (green) in a case where wrapping (red dashed line) yields the shortest distance between the two points. Right, the maximum possible distance between modules (*d*_max_) is calculated as follows: *d*_max_ = spacing of larger module * tan(π/6). The normalized distance (*d*_norm_) between any module pair is defined as the pairwise distance between those modules (*d*) divided by the maximum possible distance between those modules (*d*_max_).

**Extended Data Fig. 4.**
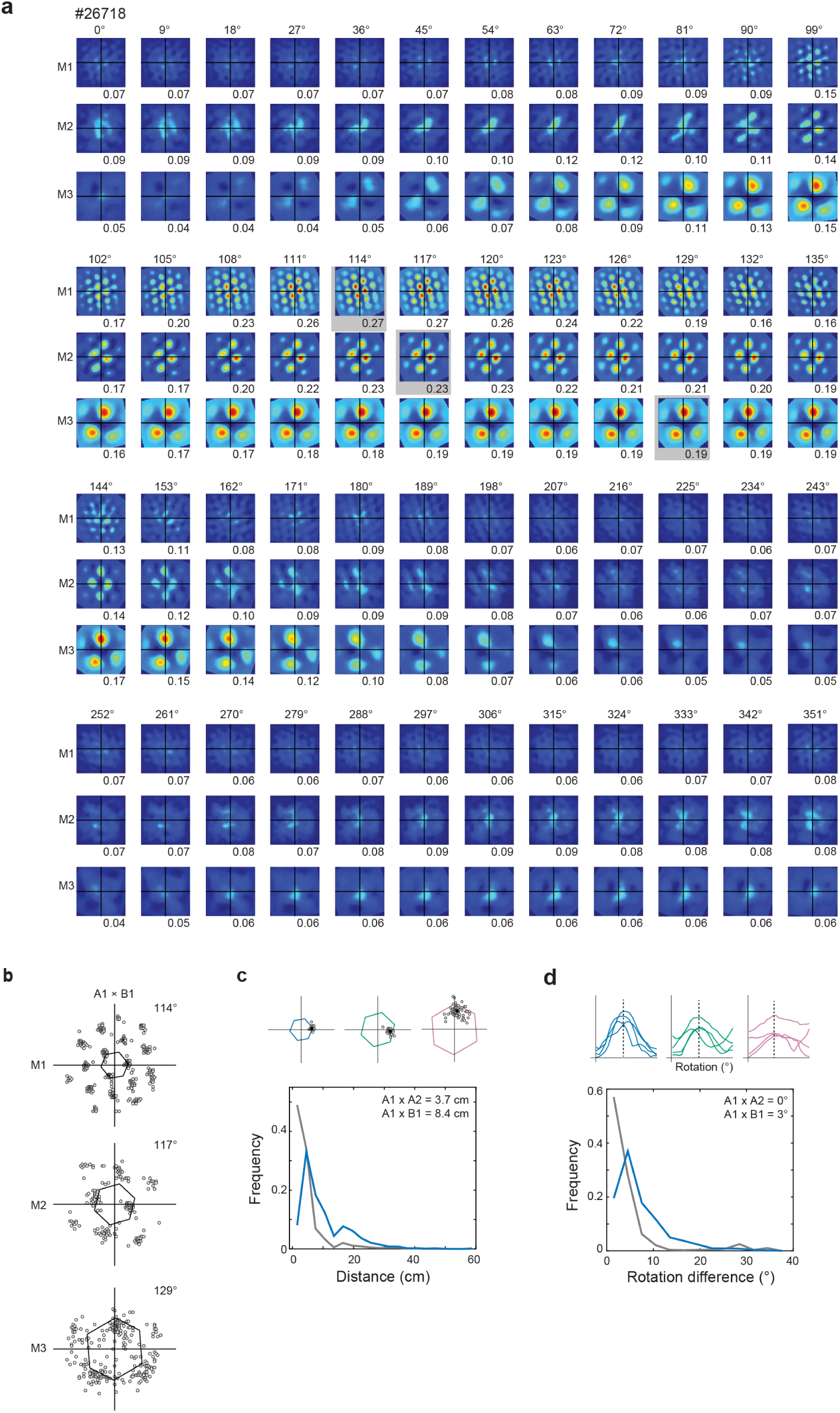
Realignment of grid cells between and within modules. **a**, Crosscorrelograms for each grid module comparing sessions in different rooms (A1×B1; three modules in rat #26718) at incremental amounts of rotation. Crosscorrelograms are depicted every 9° from 0° through 360°. Bin size for realignment analysis was 3°. Maximum correlation value of crosscorrelogram is noted below each panel. Gray box surrounds spatial crosscorrelogram containing the maximum correlation for each module (at 114, 117 and 129°). The location with the highest correlation within the spatial crosscorrelogram at the rotation of that module defined the phase displacement (shift) vector of the grid map. **b**, Within each grid module, grid cells underwent similar changes in phase. In each panel, circles depict the location of the maximum correlation in the crosscorrelograms of each grid cell in the three grid modules recorded from rat #26718 between sessions in different rooms (A1×B1). Intensity of grayscale reflects cells with the same change in phase. **c**, Top, we calculated the distance between the location of the peak within the central unit hexagon for individual grid cells and the location of the peak of the module they belong to (i.e., intramodular differences). Bottom, line graph shows intramodular differences in phase change between sessions (A1×A2, n = 1769 cells, median = 3.7 cm, 95% CI: 0.0 - 3.8 cm; A1×B1, n = 1807 cells, median = 8.4 cm, 95% CI: 7.5 - 8.4 cm). Blue line compares sessions in different rooms (A1×B1); gray line compares sessions in the same room (A1×A2). **d,** Top, we calculated the difference in rotation between individual grid cells and the rotation of the module they belong to (i.e., intramodular differences). Note that only four cells are shown in each panel (for visualization purposes only). Bottom, line graph shows intramodular differences in rotation between sessions (A1×A2, n = 1806 cells, median = 0°, 95% CI: 0 - 0°; A1×B1, n = 1891 cells, median = 3°, 95% CI: 3 - 3°). Blue line compares sessions in different rooms (A1×B1); gray line compares sessions in the same room (A1×A2).

**Extended Data Fig. 5.**
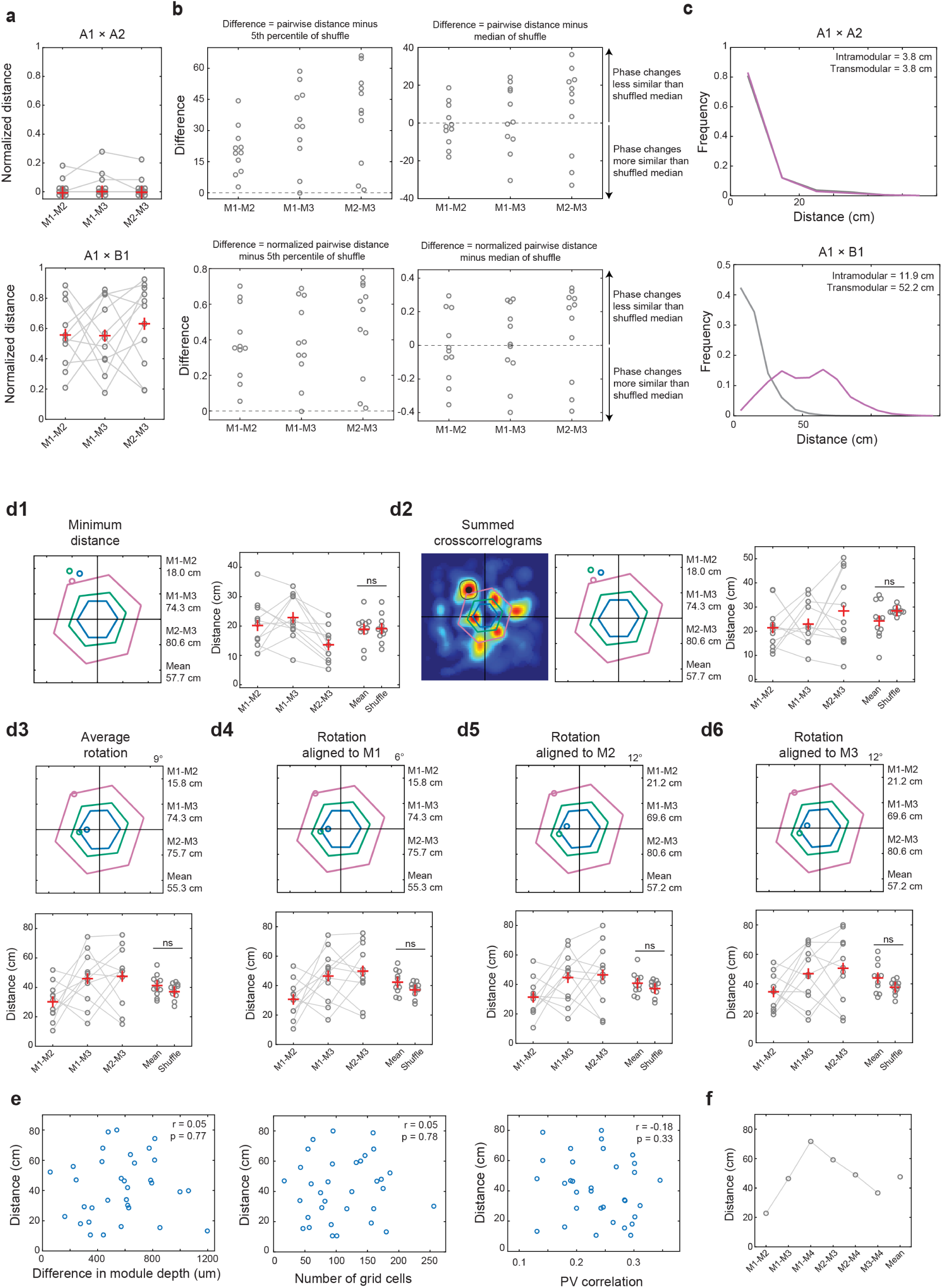
Independent phase displacements across modules are evident across all metrics used to quantify phase change. **a**, Normalized pairwise difference in phase change of grid modules between sessions in the same room (A1×A2, top) or different rooms (A1×B1, bottom) for all 11 rats shown in Figure 1. Difference in phase change of each module pair is normalized by the maximum possible distance between those modules. Lines represent distance between simultaneously recorded module pairs. Red crosses indicate means. **b**, Comparison of the observed pairwise distances between module phase changes and the pairwise distances between module phase changes in a shuffled dataset. Points show the difference between the pairwise distances for each module pair and the 5^th^ percentile (top left) or the median (top right) of the shuffled distribution for that module pair in that animal. Panels in the bottom row show the same comparison, but pairwise distances are normalized by the maximum possible distance between modules. Note that the distance between module phase changes was below the 5^th^ percentile of the shuffled distribution for only 1/33 module pairs (left), i.e. the pairwise distance was smaller than the range of values expected by chance in only one comparison. The pairwise distance between module phase changes was less than the median of the shuffled dataset for just 12/33 module pairs. **c**, As an alternative to crosscorrelations for the entire module as a unit (Extended Data Fig. 3a), we calculated the distance between phase changes of intramodular cell pairs (from the same grid module; gray) and transmodular cell pairs (from different modules; purple) across sessions in the same room (A1×A2, top) or different rooms (A1×B1, bottom) for all 11 rats shown in Figure 1. Line graphs show that the distance between grid cell pairs was significantly larger for transmodular than intramodular cell pairs when comparing sessions in different rooms (A1×B1) (transmodular: n = 105,071 cell pairs, median = 52.2 cm, 95% CI: 52.1 - 52.5; intramodular: n = 142,749 cell pairs, median = 11.9 cm, 95% CI: 11.8 - 11.9; Z = −369.1, p = 2.0 × 10^−300^, Cohen’s *d* = 2.2, two-sided Wilcoxon rank-sum test). When comparing sessions in the same room (A1×A2), the medians of the distributions of distances between intramodular and transmodular grid cell pairs were equal (transmodular, n = 100,932 cell pairs, median = 3.8 cm, 95% CI: 3.8 - 3.8; intramodular: n = 130,334 cell pairs, median = 3.8 cm, 95% CI: 3.8 - 3.8; Z = 5.8, p = 8.2 × 10^−9^, Cohen’s *d* = −0.07; two-sided Wilcoxon rank-sum test). **d1-d6**, Comparison of methods used to measure difference in phase change for the entire ensemble of each module. **d1,** Alternative method for measuring phase displacement. The minimum distance method defines the phase change of each module as the location of the peak within the spatial crosscorrelogram where the distance between all module pairs is lowest, not confining the measurement to the central hexagon. **d2**, In a second alternative approach, we calculated a summed crosscorrelogram by adding the spatial crosscorrelograms of all simultaneously recorded modules together at the rotation with the highest correlation. Crosscorrelograms were normalized to control for the number of cells per module. If grid modules shifted coherently between sessions, we expected to see a clear peak in the summed crosscorrelogram, indicating the phase change of all modules. The phase change of each module would be near the identified peak, and the difference in phase changes across modules would be small. In contrast, if grid modules shifted separately, we expected to see a random distribution of peaks in the summed crosscorrelogram. In this case, the pairwise distance between module phases would still be large. For each experiment, we identified the location of the peak in each module that was closest to the location with the maximum correlation in the summed crosscorrelogram. Note that distances are not significantly different from that of a distribution of shuffled phase changes, supporting the interpretation that grid modules shift independently. **d3-d6**, Central hexagon method defines the phase change of each module as the location of the peak within the central unit hexagon, as illustrated in Extended Data Fig. 3b. **d3**, We estimated the change in phase at the average rotation across modules, rather than using the rotation of each module. **d4-d6**, Using the central hexagon method, we estimated the phase change of each module using spatial crosscorrelograms at the rotation of M1 (**d4**), M2 (**d5**), or M3 (**d6**), instead of using the rotation of each module. For the methods illustrated in the left (**d1-d2**) or top (**d3-d6**) panels, the corresponding right (**d1-d2**) and bottom (**d3-d6**) panels show the pairwise distance between phase change vectors for each module. Mirroring the results with the central hexagon method (reported in the main text), the pairwise distance between modules calculated with the alternative methods was not significantly different than the distance between modules in a shuffled dataset (minimum distance method in **d1**: 18.9 cm ± 1.5 cm vs. 19.1 cm ± 1.4 cm; mean ± s.e.m.; t(20) = 0.11, p = 0.91; summed crosscorrelogram method in **d2**: 24.2 cm ± 2.3 cm vs. 28.4 cm ± 0.50 cm; mean ± s.e.m.; t(20) = 1.8, p = 0.09; two-sided unpaired t-tests). Differences were also not apparent with different choices for rotation alignment in **d3-d6** (p values > 0.05). In experiments with repeated presentation of the same environment (A1×A2), there were no changes in the phase of grid modules (displacement = 0 cm) when using the minimum distance and superimposed crosscorrelogram methods. ns, not significant. **e**, We tested whether there was a relationship between, on one hand, the pairwise distance between the module phase changes, and, on the other hand, the difference in the mean anatomical depth of each module pair (in µm, left), the number of grid cells per module (middle), or the maximum population vector (PV) correlation (right). There was no relationship among any of these variables (n = 33 module pairs; left, r = 0.05, p = 0.77; middle, r = 0.05, p = 0.78; right, r = −0.18, p = 0.33; Pearson correlations). **f**, Pairwise distance (cm) between module phase vectors for each pair of simultaneously recorded modules in rat #28367, in which four modules were recorded. Mean across module pairs is shown at right.

**Extended Data Fig. 6.**
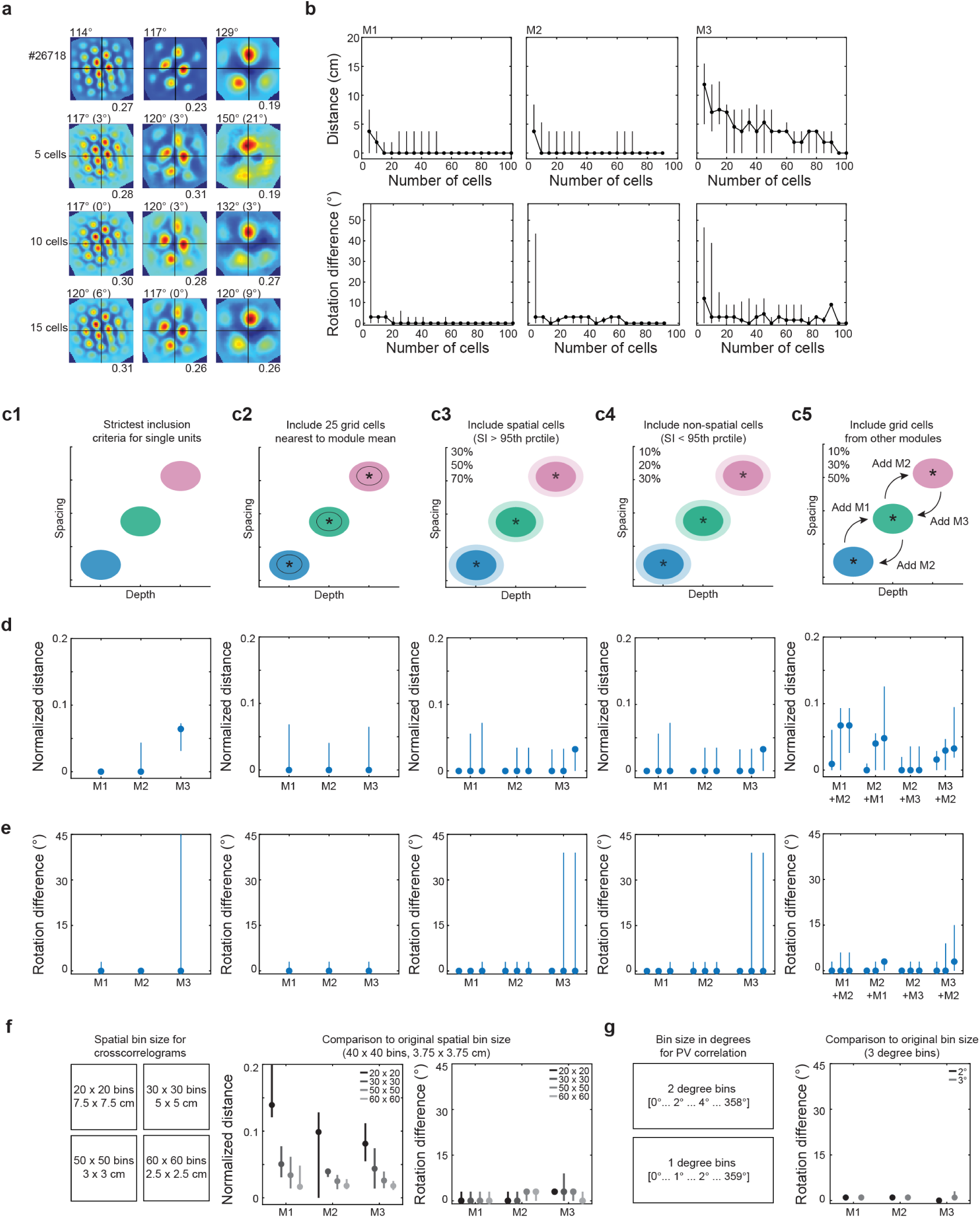
Phase displacement is robust to noise and consistent regardless of bin size or number of cells per module. **a**, Crosscorrelograms comparing sessions in different rooms for each module from rat #26718, based on either the entire module (top row), or samples of five grid cells (second row), ten grid cells (third row), or fifteen grid cells (fourth row) from that module. Grid cells in each module were sorted according to their grid scores; cells with the highest grid scores were used in each sample. Rotation of each module is noted above each panel. Maximum correlation value for each module is noted below each panel. **b**, To determine how many grid cells are needed for stable estimates of rotation and phase change, we measured phase displacement for each grid module using increasing numbers of grid cells (adding 5 cells per run, up to a maximum of 100 cells). Top row, plots show the mean difference in phase change (in cm) between each subsampled set of grid cells and the full module for all 11 rats shown in Figure 1. Panels show medians of subsampled sets of grid cells from M1, M2, and M3, respectively. Stable estimates of module phase were typically observed with just 5-10 grid cells per module (median difference in module phase using full module versus subsampled module with 10 cells: M1 = 1.9 cm, 95% CI, 0.0 - 3.8 cm; M2 = 0.0 cm, 95% CI, 0.0 - 5.3 cm; M3 = 7.1 cm, 95% CI, 3.8 - 11.9 cm). Bottom row, plots show the mean angular difference between rotation values of each subsampled set of grid cells and that of the full module for all 11 rats shown in Figure 1. Stable estimates were observed with only 5-10 grid cells per module (median difference in module rotation using full module versus subsampled module with 10 cells: M1 = 3°, 95% CI, 0 - 3°; M2 = 3°, 95% CI, 0 - 3°; M3 = 3°, 95% CI, 0 - 42°). **c1-c5**, We measured phase displacement for each grid module using more conservative or more liberal cell selection criteria to show that our measurements of module phase change and rotation between sessions are robust to noise and contamination. Criteria were based on the distribution of individual cells’ values for spatial information and anatomical position (depth). **c1**, Strictest criteria for inclusion of single units. **c2**, Inclusion of only the 25 grid cells with spacing and anatomical depth values closest to the mean depth of the module. **c3**, Inclusion of additional cells with spatial information above the 95^th^ percentile of a shuffled distribution and anatomical depth that was closest to the mean depth of the module. An increasing percentage of the original module was added in each run (30%, 50% or 70% of the total number of grid cells in the module was added to the original module). **c4**, Inclusion of additional less spatially tuned cells, defined as those with spatial information below the 95^th^ percentile of a shuffled distribution, still restricting the selection to the depths that were closest to the mean depth of the module. An increasing percentage of the original module was added in each run (10%, 20% or 30% of the total number of grid cells in the module was added to the original module). **c5**, Inclusion of additional grid cells from other modules. An increasing percentage of the original module was added in each run (10%, 30% or 50% of the total number of grid cells in the module was added to the original module). Four runs were completed in **c5**: (1) grid cells from M2 were added to M1; (2) grid cells from M1 were added to M2; (3) grid cells from M3 were added to M2; (4) grid cells from M2 were added to M3. **d**, Normalized distance between the phase change of the original module and the phase change of the adjusted module for each run shown in panels **c1-c5**. Circles show medians of all 11 rats shown in Figure 1; lines show 95% CI. Note that our estimates of module phase change did not differ until 30% or more cells were mixed in from another module, suggesting that any read-out of phase change was robust to noise. Inclusion of additional cells with or without high spatial information also did not affect our estimates of module phase or rotation. **e**, Angular difference between the rotation of the original module and the rotation of the adjusted module for each run shown in panels **c1-c5**. Circles show medians of all 11 rats shown in Figure 1; lines show 95% CI. **f**, Left, four additional sizes of spatial bins were used to compute module crosscorrelograms: (1) 20 x 20 bins (7.5 x 7.5 cm); (2) 30 x 30 bins (5 x 5 cm); (3) 50 x 50 bins (3 x 3 cm); (4) 60 x 60 bins (2.5 x 2.5 cm). Middle, difference in normalized pairwise distance between estimates obtained with the adjusted bin sizes (1-4, from left to right) and the estimate obtained with the original spatial bin size (40 x 40 bins, 3.75 x 3.75 cm). Right, difference between module rotation with the original spatial bin size and rotation values obtained with adjusted spatial bin sizes. Grayscale lines reflect spatial bin sizes (black, 20 x 20 cm; light gray, 60 x 60 cm). Circles show medians of all 11 rats shown in Figure 1; lines show 95% CI. Note that phase and rotation estimates do not deviate much from the original values despite considerable changes in bin size. **g**, Left, finer resolution of the analysis of module rotation was done using two-degree and one-degree bins. Right, difference between module rotation with the original angular bin size (three-degree bins) and rotation values obtained with adjusted angular bin sizes (2°, black; 1°, gray). Circles show medians of n = 11 rats; lines show 95% CI. Note that rotation estimates do not deviate from the original values despite changes in angular bin size.

**Extended Data Fig. 7.**
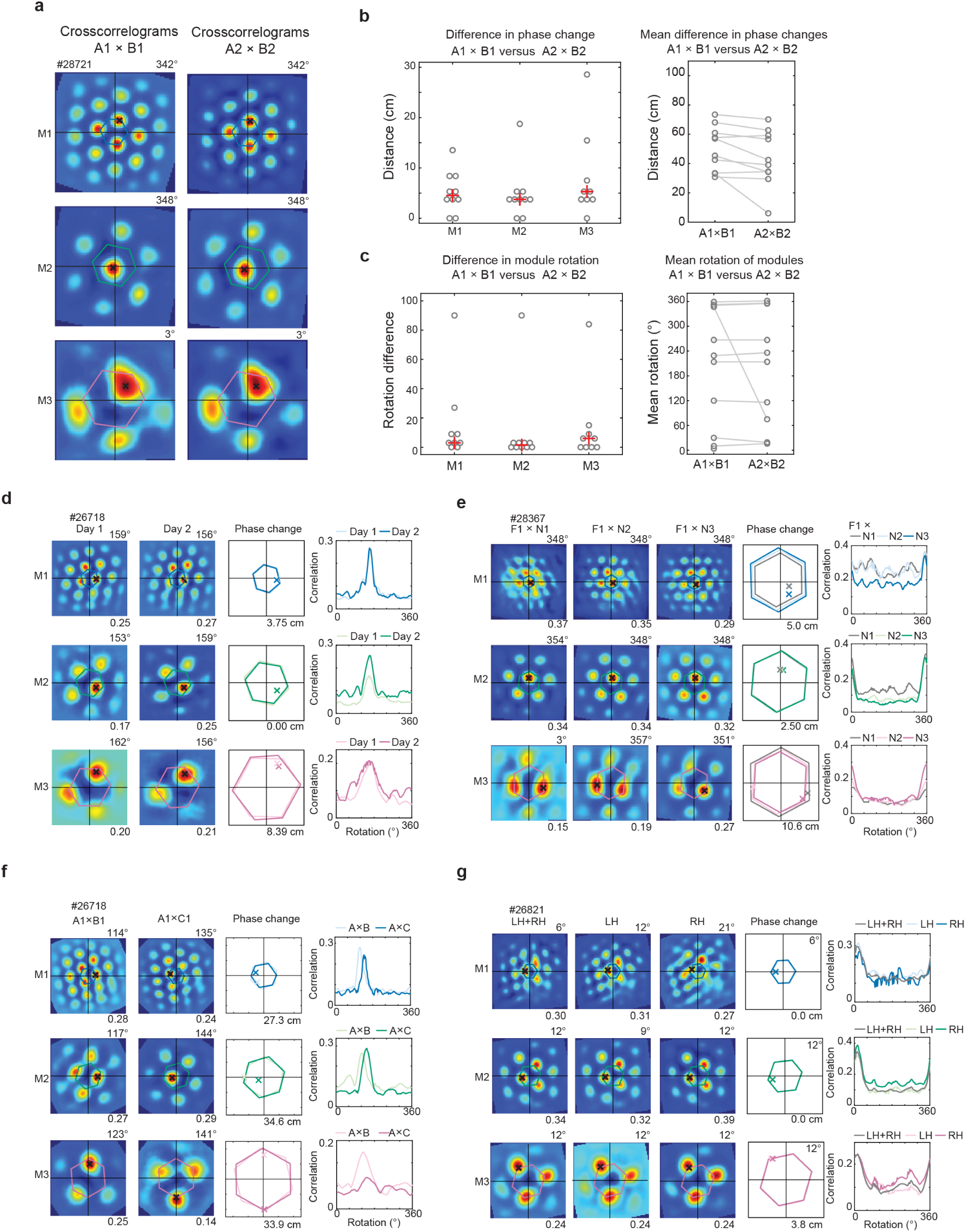
Differential phase displacements are consistent across hemispheres as well as repeated testing in rooms A and B. **a**, Crosscorrelograms for three grid modules comparing repeated sessions (1 and 2) in different rooms (A and B) in rat #28721 (left column, A1×B1; right column, A2×B2). Rotation between sessions is noted above each plot. Colored lines indicate central unit hexagon. Black crosses indicate the phase change of each module between sessions. Note similar rotation and phase change across session pairs. **b**, Left, points show the difference in phase change of each module (in cm) between A1×B1 and A2×B2. Medians are below 5 cm for all modules, indicating that modules typically shift similarly between rooms on separate occasions. Note that in one animal, the grid pattern of all three modules was unstable between sessions in room A, resulting in large distance values for all module pairs. Right, lines show the mean difference in the phase changes of simultaneously recorded grid modules between sessions in different rooms (A1×B1 and A2×B2). Red crosses indicate medians. **c**, Left, points show the difference in the rotation of each module between A1×B1 and A2×B2. Note that in one animal, the grid pattern of all three modules was unstable between sessions in room A, resulting in large rotation differences for all module pairs. Right, lines show the mean rotation of simultaneously recorded grid modules between sessions in different rooms (A1×B1 and A2×B2). Red crosses indicate medians. **d**, Phase realignment is consistent across recording days in familiar rooms. Crosscorrelograms for three simultaneously recorded modules comparing sessions in different rooms (A1×B1) in rat #26718 on two consecutive days of recording. Rotation of each module is noted above each panel. Maximum correlation value is noted below each panel. Colored lines represent the unit hexagon for each module. Black crosses indicate the phase change of each module between sessions. Third column, unit hexagons for each module on each recording day were aligned at the center and overlaid (Day 1, light colored lines; Day 2, dark colored lines). Colored crosses depict the phase change of each module between sessions. Difference in the phase change of each module between days is noted below each panel. Fourth column, plots display PV correlation at each rotation for each module. **e**, Phase realignment between a familiar (F) and an initially novel (N) room is consistent across days. Crosscorrelograms comparing the familiar room and the novel room are shown for three simultaneously recorded modules in rat #28367 on three non-consecutive days of recording (Day 1: F1×N1, gray lines; Day 2: F1×N2, light colored lines; Day 3: F1×N3, dark colored lines). Third and fourth columns as in **d**. **f**, Grid modules recorded in the same animal undergo unique realignments between different pairs of rooms. Module crosscorrelograms in rat #26718 between sessions in rooms A1 and B1 (A1×B1, first column) and between rooms A1 and C1 (A1×C1, second column) for three simultaneously recorded grid modules on two non-consecutive days of recording (A1×B1, light colored lines; A1×C1, dark colored lines). Note that the realignment of each module (in terms of its rotation and phase change) is unique between A1×B1 and A1×C1. Rooms B and C are not compared since recordings took place on different days (which impairs success in matching cell IDs and corresponding rate maps). Third and fourth columns as in **d**. **g**, Grid cells that belong to the same module in the left and right hemispheres undergo similar changes in rotation and phase between rooms. Crosscorrelograms comparing sessions in different rooms (A1×B1) are shown for three modules recorded in left and right hemispheres (LH+RH: left column, gray lines) in rat #26821. Grid cells belonging to the same module recorded in the left hemisphere (LH: middle column, light colored lines) and right hemisphere (RH: right column, dark colored lines) realigned similarly. Third and fourth columns as in **d**.

**Extended Data Fig. 8.**
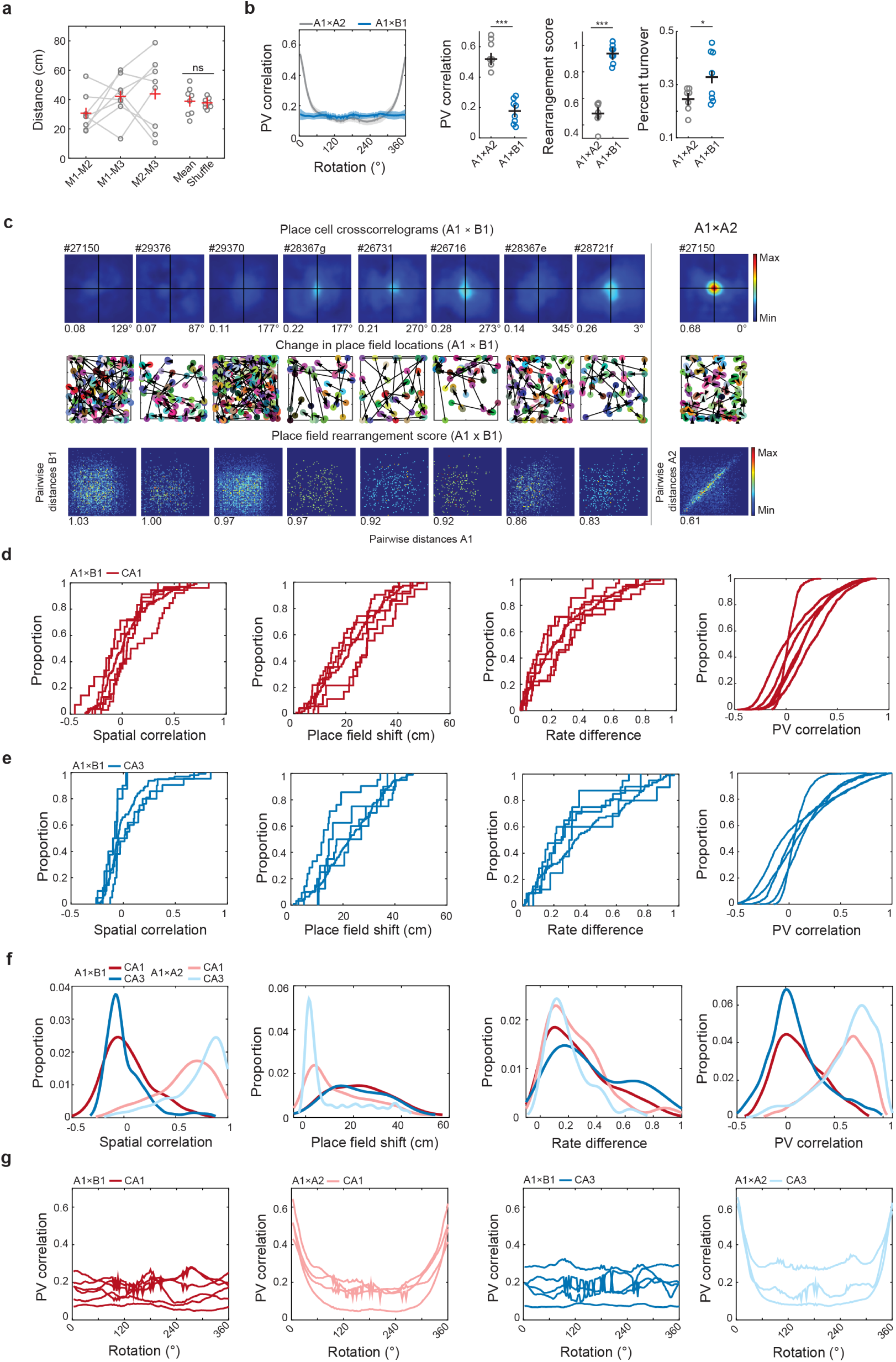
Global remapping in place cells from rats with three or more simultaneously recorded grid modules. **a**, Pairwise distance (in cm) between module phase changes (as in Fig. 1h) for each experiment with simultaneously recorded place cells and multiple grid modules (n = 8 experiments). Mean across module pairs is shown for each experiment next to the mean of the shuffled distribution for that experiment (as in bottom panel of Fig. 1e). Four modules were recorded in rat #28367 (see Fig. 1f). Pairwise distances between M4 and the remaining modules are shown in Extended Data Fig. 5f and were included in the mean for this animal. Red crosses indicate means. ns, not significant. **b**, Left, PV correlation at each rotation for sessions in the same room (A1×A2, gray) or in different rooms (A1×B1, blue; shaded region represents mean ± s.d.) in each experiment with simultaneous recordings of place cells and multiple grid modules. Right, panels show maximum PV correlation, place field rearrangement scores, and percent turnover between sessions in the same room (A1×A2, gray) versus sessions in different rooms (A1×B1, blue) for each experiment with simultaneous recordings of place cells and multiple grid modules. PV correlations were significantly reduced, while place field rearrangement scores and percent turnover were significantly increased between sessions in different rooms (PV correlation: A1×A2, 0.56 ± 0.04; A1×B1, 0.17 ± 0.03; mean ± s.e.m.; A1×A2 vs. A1×B1, t(6) = 8.6, p = 5.1 × 10^−7^; rearrangement scores: A1×A2, 0.49 ± 0.03; A1×B1, 0.94 ± 0.02; mean ± s.e.m.; t(6) = 8.5, p = 7.3 × 10^−5^; percent turnover: A1×A2, 0.25 ± 0.02; A1×B1, 0.34 ± 0.03; mean ± s.e.m.; A1×A2 vs. A1×B1, t(6) = 2.9, p = 0.01; one-sided paired t-tests). Black crosses indicate means. **c**, Population measures of global remapping between sessions in different rooms (A1×B1) in experiments with simultaneous recordings of place cells and multiple grid modules. Each column shows data from one experiment (animal ID noted at top). Top row, place cell crosscorrelograms at the rotation with the highest correlation. Maximum PV correlation and rotation are noted below each panel. Note that in four cases, a light blue peak is visible at the origin of the crosscorrelogram. While this may suggest that remapping was incomplete, the remaining metrics used to quantify remapping (as well as place cell rate maps) indicate that the effect was very weak and that global remapping occurred. Middle row, black lines show the change in field location for each place cell between sessions. Bottom row, pairwise distance between place field locations in session A versus pairwise distance between place field locations in session B (i.e., place field rearrangement score) is shown as a scatter plot. Data are plotted as a 2D histogram; color indicates number of place field pairs per bin (from blue to red, scale bar). Rearrangement score is noted below each panel. Note the absence of a clear peak at the origin of the crosscorrelogram and the random reorganization of place field locations between sessions. Rightmost column compares sessions in the same room (A1×A2) in one animal. Note clear peak at the origin of the crosscorrelogram (top), minimal changes in place field location (middle), and the strong correlation along the diagonal between pairwise place field distances (bottom), indicating that place cells were stable between sessions. **d**, Cumulative distribution function (CDF) plot of spatial correlation (first column), change in place field location (second column), change in mean firing rate (third column), and population vector (PV) correlation (fourth column) for experiments in which CA1 place cells were simultaneously recorded with three or more grid modules. Population vector correlations were calculated by stacking all place cell rate maps (including those of the silent cells) into a three-dimensional matrix with cell identity on the z-axis. The distribution of mean rates along the z-axis for a given xy location (i.e., one spatial bin) represents the population vector for that location. Comparing the entire set of population vectors between two trials provides an estimate of how much the ensemble code changed between sessions. Each curve shows one experiment (7 recordings in 6 animals). The distributions are overlapping in each panel, indicating that the degree of remapping is similar in each experiment. **e**, Same panels as in **d** but for experiments in which CA3 place cells were simultaneously recorded with three or more grid modules Each curve shows one experiment (5 recordings in 4 animals). Note the similarity of the distributions in each panel. **f**, Frequency distribution of spatial correlation (first column), change in place field location (second column), change in mean firing rate (third column), and PV correlation (fourth column) for the entire population of CA1 (red) and CA3 (blue) place cells between sessions in the same room (A1×A2, lighter colors) versus sessions in different rooms (A1×B1, darker colors). Spatial correlation of both CA1 and CA3 place cells is significantly lower between sessions in different rooms (A1×B1) than between sessions in the same room (A1×A2) (A1×B1 vs. A1×A2, CA1: Z = 15.7, p = 1.0 × 10^−55^, d = 2.4; CA3: Z = 13.7, p = 6.4 × 10^−43^, d = 2.9; one-sided Wilcoxon rank-sum tests), but does not differ between subregions (CA1 vs. CA3: A1×B1, Z = 1.6, p = 0.12; two-sided Wilcoxon rank-sum test). Rate changes in both CA1 and CA3 were significantly greater between sessions in different rooms than between sessions in the same room (A1×B1 vs. A1×A2, CA1: Z = 3.7, p = 1.2 × 10^−4^, d = 0.38; CA3: Z = 6.4, p = 8.6 × 10^−11^, d = 0.92; one-sided Wilcoxon rank-sum tests) and were slightly (but significantly) greater in CA3 than in CA1 (A1×B1, CA1 vs. CA3: Z = 3.2, p = 1.2 × 10^−3^, d = 0.38; two-sided Wilcoxon rank sum test). Changes in place field location were significantly larger between sessions in different rooms than between session in the same room in CA1 and CA3 (A1×B1 vs. A1×A2, CA1: Z = 7.9, p = 1.8 × 10^−15^, d = 0.76; CA3: Z = 8.5, p = 1.2 × 10^−17^, d = 0.98; one-sided Wilcoxon rank-sum tests), but did not differ between subregions (CA1 vs. CA3, A1×B1: Z = 0.49, p = 0.62; two-sided Wilcoxon rank-sum test). PV correlations were significantly lower between sessions in different rooms than between sessions in the same room in CA1 and CA3 (A1×B1 vs. A1×A2, CA1: Z = 73.1, p < 1.0 × 10^−16^, d = 1.7; CA3: Z = 63.3, p < 1.0 × 10^−16^, d = 1.9; one-sided Wilcoxon rank-sum tests) and were slightly (but significantly) lower in CA3 than in CA1 (CA1 vs. CA3, A1×B1: Z = 11.5, p = 9.2× 10^−31^, d = 0.15; two-sided Wilcoxon rank-sum test). **g**, Lines show the PV correlation at each rotation for experiments with CA1 (red) or CA3 (blue) place cells. Each line shows one experiment (all cells from the region). Darker colors show PV correlation between sessions in different rooms (A1×B1, first and third columns). Lighter colors show PV correlation between sessions in the same room (A1×A2, second and fourth columns).

**Extended Data Fig. 9.**
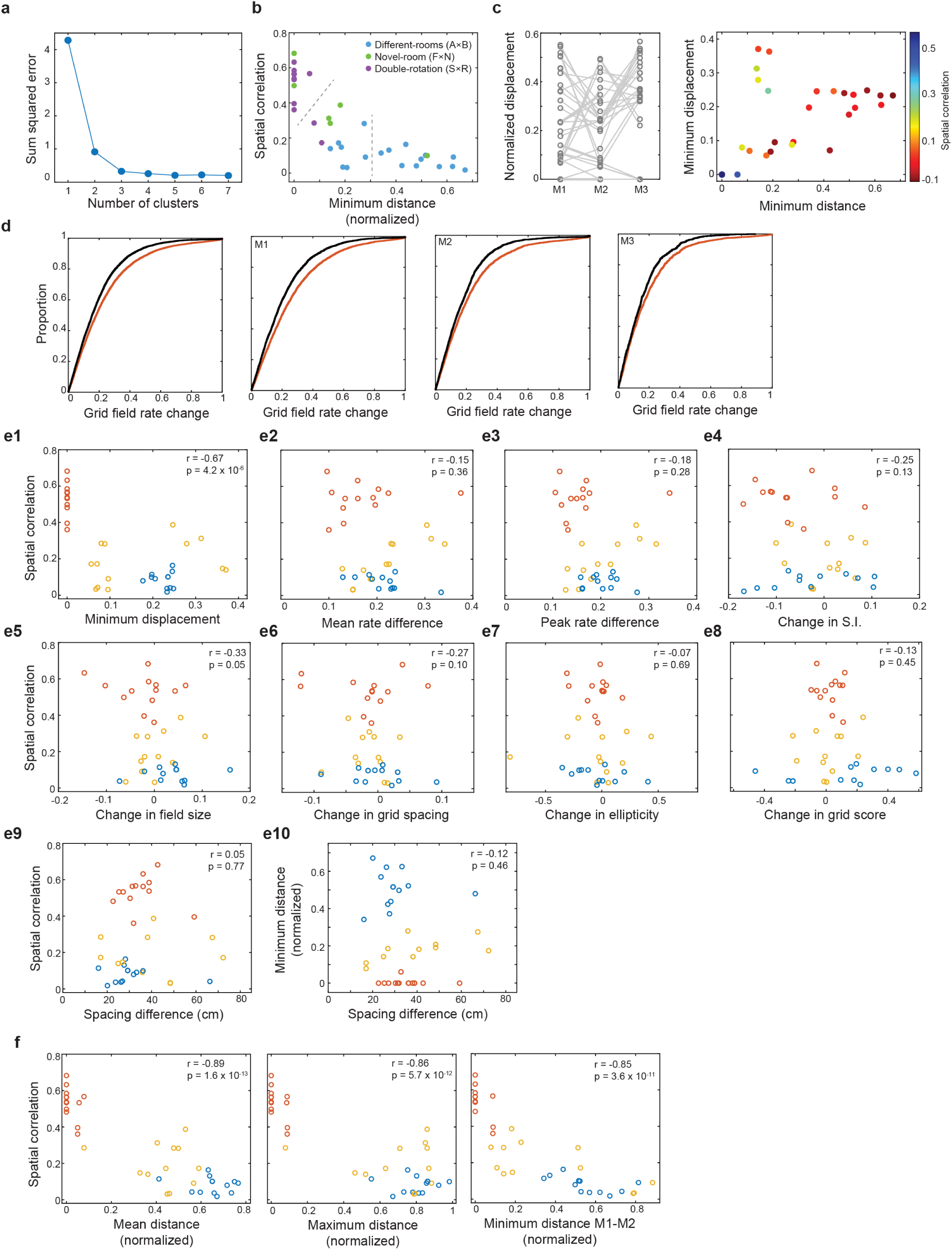
Minimum distance between phase change vectors across module pairs is the clearest predictor of the degree of remapping. **a**, Number of clusters in *k*-means clustering. We applied *k*-means clustering to separate the datapoints in distributions showing the spatial correlation of place cells as a function of the minimum normalized distance between module phase changes. The number of clusters was determined empirically. We calculated the sum squared error to evaluate the quality of the clustering after varying the number of clusters (k) from one to seven. The point at which the sum squared error starts to decrease at a slower rate indicates the optimal number of clusters (‘elbow method’). The empirically-determined *k*-value based on this metric was three. **b**, Experimental conditions included in each group separated using *k*-means clustering. Each point represents one session pair; points are colored according to experimental condition (blue = ‘different-rooms’ task, A×B; green = ‘novel-rooms’ task, F×N; purple = ‘double-rotation’ task, S×R). Dashed gray lines show separation of groups based on *k*-means clustering. The spatial correlation values of place cells were significantly different between conditions (A×B, median = 0.08, 95% CI, 0.05 - 0.11; F×N, median = 0.38, 95% CI, 0.32 - 0.45; S×R, median = 0.57, 95% CI, 0.54 - 0.60; A×B vs. F×N, Z = 11.2, p = 3.6 × 10^−29^; F×N vs. S×R, Z = 8.5, p = 1.3 × 10^−17^; two-sided Wilcoxon rank sum tests). **c**, Left, normalized displacement of simultaneously recorded grid modules (n = 38 session pairs). Modules from the same recording are connected by lines. Normalized displacement is expressed as Euclidean distance from the origin to the location in the central hexagonal tile of the crosscorrelogram with the highest correlation value, divided by the spacing of the module. Right, scatter plot comparing the minimum normalized phase distance with the minimum normalized displacement for all experiments. Points are colored according to the mean spatial correlation of place cells (scale bar). The strength of remapping increased (blue to red) with increasing changes in grid phase and disparity among grid modules. Note that the points for 11 session pairs are overlapping in the bottom left corner (11 data points), where the minimum displacement and minimum distance between modules is zero. Therefore, in these sessions, there were no changes in module phase. This was most frequently observed when local and distal cues were rearranged within the same room and was associated with weak place cell remapping. Although the extent of remapping in response to similar cue manipulations is often stronger than we show here^18,68^, it is conceivable that stronger remapping in those studies was also associated with both rotation and translation of the grid pattern. **d**, For session pairs in which grid modules did not undergo a change in phase, we were able to track the location of each grid field across sessions. We calculated the change in the peak firing rate within each field between sessions in the standard and rotated configuration (S1×R1) (red lines) and between sessions in the standard configuration (S1×S2) (black lines). Panels show that changes in grid field rate are greater between sessions with different cue configurations than between sessions in the same configuration for all modules (first panel), M1 (second panel), M2 (third panel), and M3 (fourth panel). Note that changes in the firing rate of individual grid fields were associated with weak remapping in response to the double rotation of local and distal cues. **e1-e9**, The minimum normalized distance between module phase changes and the minimum normalized displacement were the only variables that were significantly correlated with the extent of place cell remapping. **e1**, Scatter plot shows a significant negative correlation between the minimum phase displacement of grid modules and the spatial correlation of place cells (n = 38 session pairs, r = −0.67, p = 4.2 × 10^−6^, Pearson correlation). Points are colored according to the *k*-means clustering into groups described above. There was no correlation between the spatial correlation of place cells and any of the following measures: changes in the mean firing rate of grid cells (rate difference is equivalent to the absolute change in mean firing rate) (**e2**, r = −0.15, p = 0.36, Pearson correlation), changes in the peak firing rate of grid cells (**e3**, r = −0.18, p = 0.28, Pearson correlation), changes in the spatial information of grid cells (**e4**, r = −0.25, p = 0.13, Pearson correlation), changes in the size of grid fields (**e5**, r = - 0.33, p = 0.05, Pearson correlation), changes in grid spacing (**e6**, r = −0.27, p = 0.10, Pearson correlation), changes in the ellipticity of the grid pattern (**e7**, r = −0.07, p = 0.69, Pearson correlation), changes in grid score (**e8**, r = −0.13, p = 0.45, Pearson correlation), or the difference in the spacing of module pair with the most similar change in phase (**e9**, r = 0.05, p = 0.77, Pearson correlation). **e10**, There was also no correlation between the minimum distance between module phase changes and the spacing difference (in cm) between those modules (r = −0.12, p = 0.46, Pearson correlation). **f**, All metrics used to quantify the disparity among the phase changes of grid modules showed a clear relationship with the extent of hippocampal remapping. Scatter plots shows a significant negative correlation between the mean normalized distance (left) and maximum normalized distance (middle) between module phase changes and the spatial correlation of place cells (n = 38 session pairs; mean distance: r = −0.89, p = 1.6 × 10^−13^; maximum distance: r = −0.86, p = 5.7 × 10^−12^; Pearson correlations). Right, in 24/38 (63%) session pairs, M1 and M2 underwent the most similar changes in phase. Therefore, there was a significant correlation between the minimum distance between the phase changes of M1 and M2 and the spatial correlation of place cells (n = 38 session pairs; r = −0.85, p = 3.6 × 10^−11,^ Pearson correlation). Note that the points are colored according to the *k*-means clustering into groups described above and that the relative grouping of session pairs remains nearly unchanged.

**Extended Data Fig. 10.**
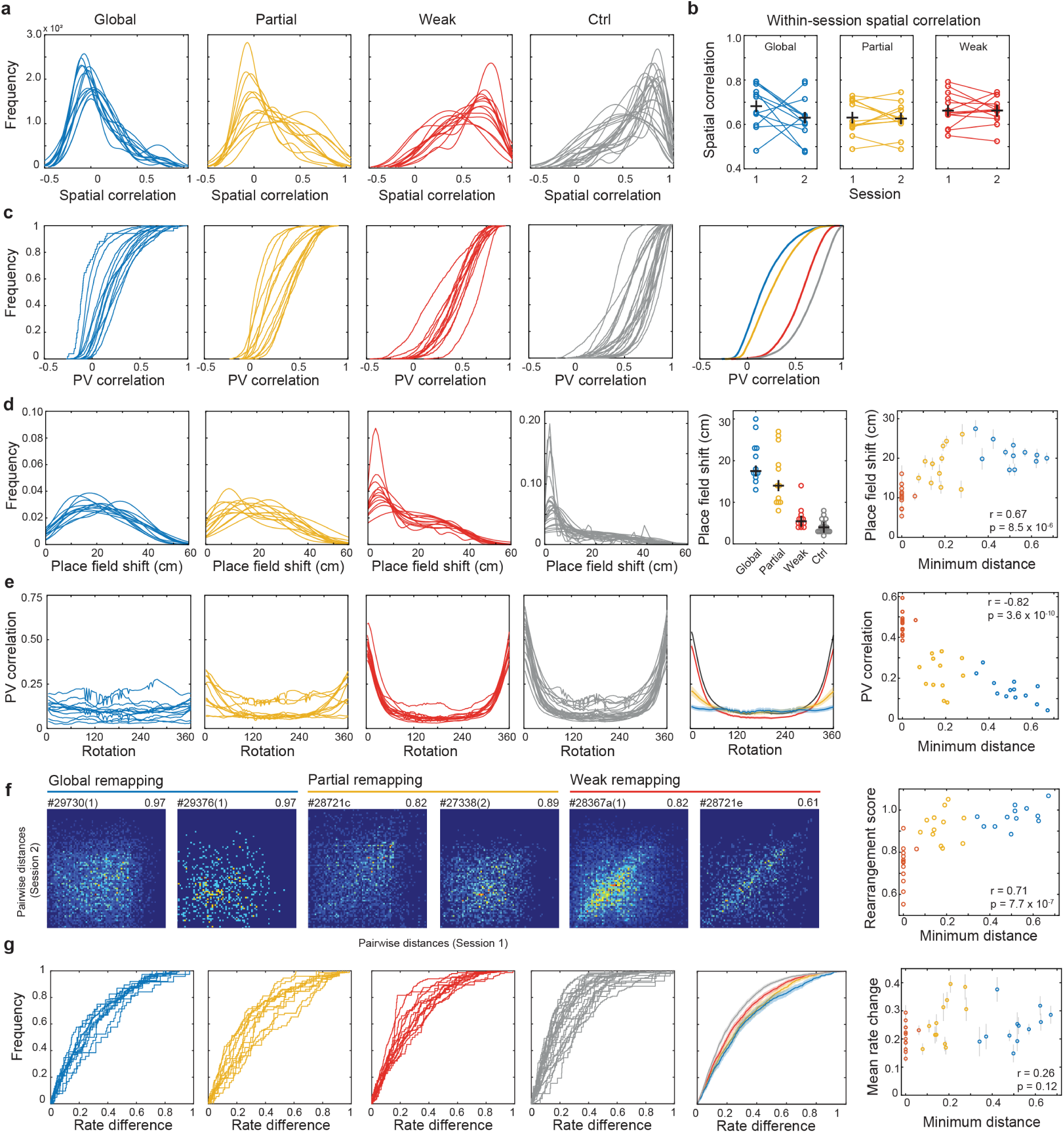
Extent of place cell remapping increases with greater disparity among the phase changes of recorded modules. **a**, Frequency distribution of spatial correlation of place cells for session pairs in each group. Each line shows data from one session pair. Session pairs are separated into groups across panels using the *k*-means clustering described in Fig. 3 and Extended Data Fig. 9a (Global, blue, n = 13; Partial, yellow, n = 12, Weak, orange, n = 13, Ctrl, gray, n = 23). **b**, Panels show the spatial correlation between the first and second halves of each session for session pairs included in the following groups: Global (left, blue), Partial (middle, yellow), Weak (right, orange). Lines connect sessions from the same recording. There was no significant difference in the within-session stability of place cells in any group (F(1,2) = 1.1, p = 0.35), and there was no difference in within-session stability between groups (F(2,33) = 1.5; p = 0.12; one-within, one-between repeated measures ANOVAs). Red crosses indicate means. **c**, CDF plots show the PV correlation of place cells between sessions. PV correlations were calculated by stacking all place cell rate maps (including those of the silent cells) into a three-dimensional matrix with the two spatial dimensions on the x and y axes and cell identity on the z-axis. The distribution of mean rates along the z-axis for a given x-y location (i.e., one spatial bin) represents the population vector for that location. Comparing the entire set of population vectors between two trials provides an estimate of how much the ensemble code changed between sessions. Each line shows data from one session pair. Session pairs are separated into groups as in **a**. Fifth column, CDF plot shows the mean PV correlation of each group (Global, n = 15,293 pixels; Partial, n = 14,448 pixels; Weak, n = 15,024 pixels; Ctrl, n = 20,506 pixels; Global vs. Partial: D = 0.16, p < 2.2 × 10^−16^; Partial vs. Weak: D = 0.51, p < 2.2 × 10^−16^; Weak vs. Ctrl: D = 0.25 p < 2.2 × 10^−16^; two-sided Kolmogorov-Smirnov tests). **d**, Frequency distribution of place field shift (in cm) of all place cells with detectable fields in both sessions. Each line shows data from one session pair. Session pairs are separated into groups as in **a**. Fifth column, mean change in place field location for each experiment separated by group. Changes in place field location in all groups were significantly higher than in the Ctrl group (Global, n = 13; Partial, n = 12; Weak, n = 13; Ctrl, n = 23; Global vs. Ctrl, Z = 4.8, p = 1.4 × 10^−6^; Partial vs. Ctrl, Z = 4.8, p = 1.5 × 10^−6^; Weak vs. Ctrl, Z = 2.6, p = 8.7 × 10^−3^; two-sided Wilcoxon rank-sum tests). Sixth column, scatter plot shows significant correlation between the mean change in place field location and the minimum distance between phase shifts across pairs of modules (n = 38 session pairs, r = 0.67, p = 8.5 × 10^−6^, Pearson correlation). **e**, PV correlation at each rotation for place cells. Since place cells did not consistently rotate between sessions, the strength of the PV correlation at 0° provides a population-level metric to quantify the strength of remapping in each group. Each line shows data from one experiment. Session pairs are separated into groups as in **a**. Fifth column, PV correlation at each rotation for place cells in each group (shaded region represents mean ± s.d.). The maximum PV correlation significantly decreased as the disparity between modules increased (Global vs. Partial, Z = 2.6, p = 8.9 × 10^−3^; Partial vs. Weak, Z = 8.5, p = 7.1 × 10^−9^; Weak vs. Ctrl, Z = 2.8, p = 3.7 × 10^−3^; one-sided Wilcoxon rank-sum tests). Sixth column, scatter plot shows significant correlation between the maximum PV correlation of place cells and the minimum phase-shift distance between modules (n = 38 session pairs, r = −0.82, p = 3.6 × 10^−10^, Pearson correlation). **f**, Heat map of scatter plot showing the pairwise distance between place field locations in each session (for cells with fields identified in both recordings). Two representative experiments are shown per group (Global, blue; Partial, yellow; Weak, orange). Data are plotted as a 2D histogram; color indicates number of place field pairs per bin (from blue to red, scale bar). Animal ID and place field rearrangement score are noted above each plot. Scatter plot shows significant correlation between the rearrangement score and the minimum phase-shift distance between modules (n = 38 session pairs, r = 0.71, p = 7.7 × 10^−7^, Pearson correlation). **g**, CDF plots show the change in mean firing rate of place cells between sessions. Each line shows data from one session pair. Session pairs are separated into groups as in **a**. Fifth column, CDF plot shows the mean change in firing rate for all place cells included in each group. Changes in firing rate in all groups were significantly higher than in the Ctrl group (Global, n = 649; Partial, n = 797; Weak, n = 1682; Ctrl, n = 1926; Global vs. Ctrl: Z = 6.0, p = 1.7 × 10^−9^; Partial vs. Ctrl, Z = 8.2, p = 3.4 × 10^−16^; Weak vs. Ctrl, Z = 4.5, p = 7.2 × 10^−6^; two-sided Wilcoxon rank-sum tests). Shaded regions show 95% CI. Sixth column, scatter plot shows no correlation between the change in mean firing rate and the minimum distance between phase-change vectors of simultaneously recorded modules (n = 38 session pairs, r = 0.26, p = 0.12, Pearson correlation).

**Extended Data Fig. 11.**
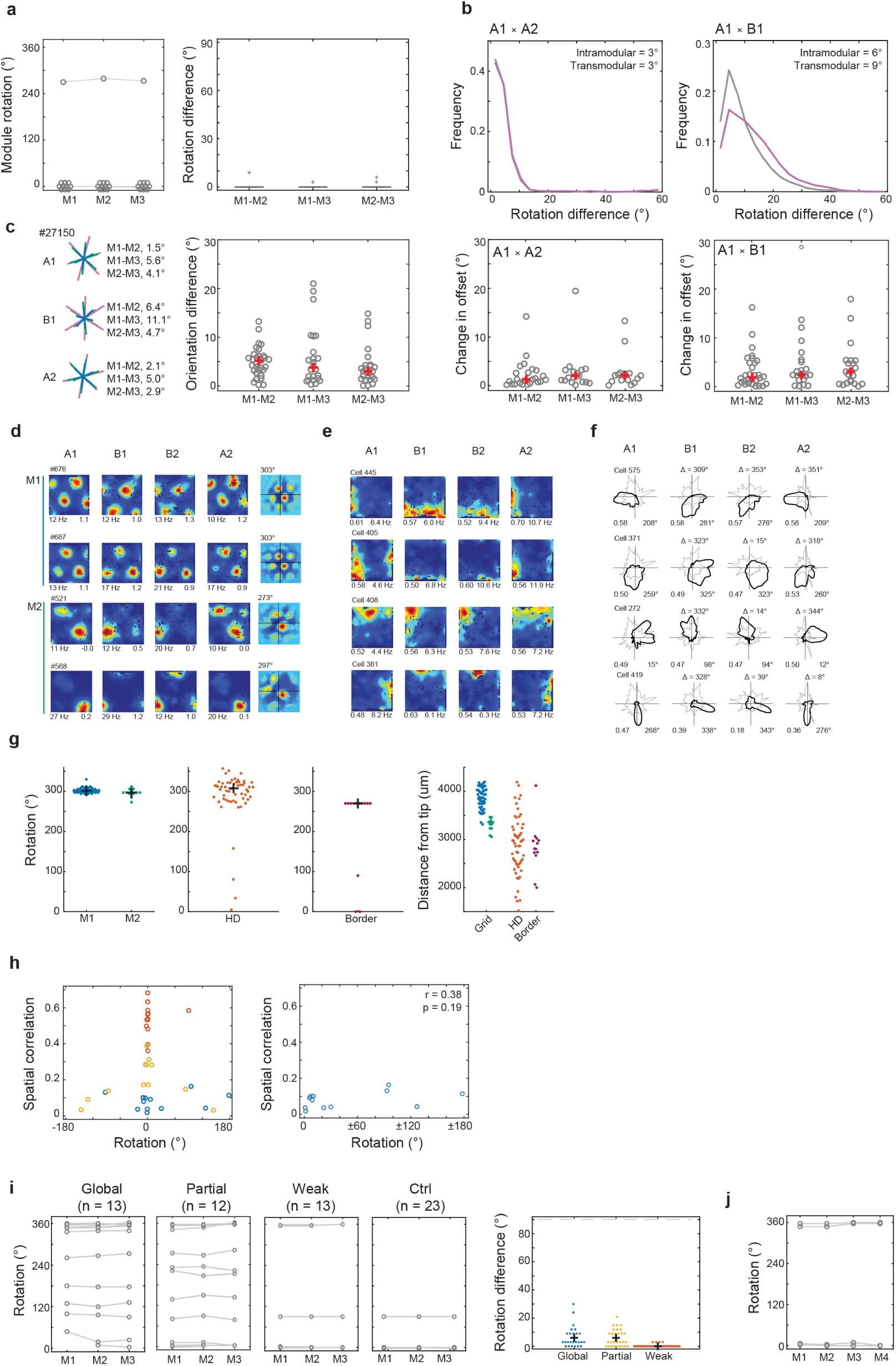
Differential rotation of grid modules does not contribute to remapping in the hippocampus. **a**, Rotation of simultaneously recorded grid modules across sessions in the same room (A1×A2) for all 11 rats included in Figure 1. Left, gray circles indicate rotation with the highest correlation between rate maps from A1 and A2. Gray lines connect simultaneously recorded grid modules. Note that in one animal, the grid pattern of all three modules was unstable between sessions in the same room (rotation = 270°). In the remaining animals, there was no rotation of grid modules between sessions in the same room; circles are offset for visualization only. Right, boxplot shows the difference in rotation between simultaneously recorded module pairs between sessions in the same room (A1×A2). **b**, Histograms show similar intramodular (gray) and transmodular (purple) differences in rotation for all grid cell pairs between sessions in the same room (A1×A2, left) or different rooms (A1×B1, right) for all 11 rats included in Figure 1. Intramodular differences are defined as the difference in rotation for pairs of grid cells in the same module. Transmodular differences are defined as the difference in rotation for pairs of grid cells recorded in different modules. Transmodular differences in rotation were slightly (and significantly) larger than intramodular differences, both when comparing sessions in the same room (medians of 3.0° and 3.0°, respectively; Z = 5.4, p = 7.6 × 10^−8^, d = 0.01) and sessions in different rooms (medians of 9.0° and 6.0°, respectively; Z = 83.6, p < 1.0 × 10^−16^, d = 0.39; two-sided Wilcoxon rank-sum tests). **c**, Orientation differences between modules are generally small and do not change between rooms. First panel, orientation of each grid axis in three simultaneously recorded modules from rat #27150 in sessions A1, B1, and A2. Mean difference in orientation between each module pair is noted at right. Second panel, mean difference in orientation between grid modules in the first session of each recording. All experimental conditions are shown (‘different rooms’, ‘novel-rooms’, and ‘double rotation’; n = 38 session pairs; M1-M2, 5.23°, 95% CI, 3.36 - 5.83°; M1-M3: 3.78°, 95% CI, 1.98 - 5.69°; M2-M3: 3.02°, 95% CI, 1.80 - 4.06°). There was no significant difference between any module pairs (p > 0.05). There was little change in the offset between modules between sessions in the same room (third panel; M1-M2, 1.86°, 95% CI, 1.13 - 3.64°; M1-M3: 2.39°, 95% CI, 2.17 - 5.05°; M2-M3: 3.04°, 95% CI, 1.02 - 4.73°) or different rooms (fourth column; M1-M2, 1.28°, 95% CI, 0.76 - 2.69°; M1-M3: 2.04°, 95% CI, 0.79 - 3.22°; M2-M3: 2.12°, 95% CI, 1.13 - 2.54°; same vs. different room for each module pair, p > 0.05). **d**, Rate maps of 2 grid cells from M1 and 2 grid cells from M2 in rat #27207 in sessions A1, B1, B2, and A2. Peak firing rate and grid score are noted below each panel. Rightmost column shows crosscorrelograms (A1×B1) for individual grid cells. Rotation is noted above each panel. **e**, Rate maps of 4 border cells from rat #27207 in sessions A1, B1, B2, and A2 arranged according to border score in session A1. Border score^77^ and peak firing rate are noted below each panel. Note that the cells’ preferred boundary rotates counterclockwise between rooms (A1×B1), reflecting a 270° rotation. **f**, Polar plots showing firing rate as a function of head direction in 4 head direction cells from rat #27207 recorded in sessions A1, B1, B2 and A2 and arranged according to mean vector length in session A1. Mean vector length and mean angle are noted below each panel. Change in mean angle is noted above each plot. **g**, Space and direction-tuned neurons rotate coherently. First panel, Rotation of grid maps in M1 (blue) and M2 (green) when rat #27027 was moved between rooms (A1×B1). Each point represents the rotation of one grid cell. Second panel, change in mean orientation of head direction cells recorded simultaneously with grid cells as the rat was moved between rooms. Black cross indicates median. The angular difference between grid cells and head direction cells (median = 8.2°, 95% CI, 6.0 - 14.5°) was not significantly different than the angular difference among simultaneously recorded pairs of head direction cells (median = 12.3°, 95% CI, 8.0 - 17.8°; Z = 1.6, p = 0.11, two-tailed Wilcoxon rank-sum test). Third panel, rotation of border cells in rat #27207 between the two rooms. Border cells reorganized between environments by rotating to the nearest cardinal angle to the rotation of co-recorded grid and head direction cells. Given that the rotation of border cells is restricted to one of the cardinal angles, the angular difference between grid and border cells was slightly greater than the angular difference between pairs of border cells (angular difference between border and grid cells: median = 10.5°, 95% CI, 0 - 29°; angular difference between pairs of border cells = 0°, 95% CI, 0 - 0°; Z = 4.0, p = 6.2 × 10^−5^, two-tailed Wilcoxon rank-sum test; Fig. 4e). Each point represents the rotation of one border cell. Black cross indicates median. Note that grid cells, head direction cells and border cells rotated in unison. The remaining spatial and non-spatial cells in the entorhinal cortex (some of which may be grid cells with very large fields) also exhibited a modest reorganization of their firing patterns between environments (Supplementary Table 2). However, given their large field sizes and low spatial information content (Supplementary Table 2), it seems unlikely that these cells contribute to the precise positioning of place fields in each environment, and instead may represent environmental features that differ between contexts^52^ or task demands^52–53,75^. Fourth panel, distance relative to the tip of the recording probe for grid cells in M1 (n = 48, blue) and M2 (n = 15, green), head direction cells (n = 57, orange), and border cells (n = 13, red) in rat #27207. Each dot indicates one cell. **h**, Left, scatter plot shows the mean spatial correlation of place cells (a measure of remapping) as a function of the mean rotation of grid modules between sessions. Each session pair is represented by one point. Points are colored according to group (Global, blue; Partial, yellow; Weak, orange), as in Fig. 3e. Ctrl group is not shown (no rotation between sessions). Rotation is shown as ±180°. Note that non-zero changes in grid orientation were more often associated with global remapping, but that spatial correlation values near zero were still observed in the absence of grid rotation. Right, scatter plot shows mean spatial correlation of place cells as a function of the mean rotation of grid modules between sessions. Rotation is shown as the absolute value of the rotation from 0° (±0-180°). There was no relationship between the extent of remapping and the difference in rotation across modules in these experiments (‘Global’, n = 13 session pairs, r = 0.38, p = 0.19). **i**, Rotation of grid maps across modules does not influence the extent of remapping in place cells (from left: Global, n = 13 session pairs; Partial, n = 12 session pairs; Weak, 13 session pairs; Ctrl, 23 session pairs). Rotation of grid modules (all modules rotated by at least 3°, the 5^th^ percentile of a shuffled distribution) was most frequent between sessions when place cells underwent global remapping (Global, 11/13 session pairs; Partial, 8/12 session pairs; Weak, 1/13 session pairs; Ctrl, 1/23 session pairs). Despite this, rotation differences between modules were very small in all groups (Global: median = 6°, 95% CI, 3 - 6°; Partial: median = 6°, 95% CI, 3 - 9°; Weak: median = 0°, 95% CI, 0 - 0°; Ctrl: median = 0°, 95% CI, 0 - 0°). Right, difference in rotation between simultaneously recorded module pairs in each group (each pair indicated by a circle). Black crosses indicate medians. Dashed gray line at 90° represents chance. **j**, Rotation of all four simultaneously recorded modules between sessions in rat #28367. Lines connect data from a single recording. Note that the rotation of the fourth module did not differ from the remaining modules.

**Extended Data Fig. 12.**
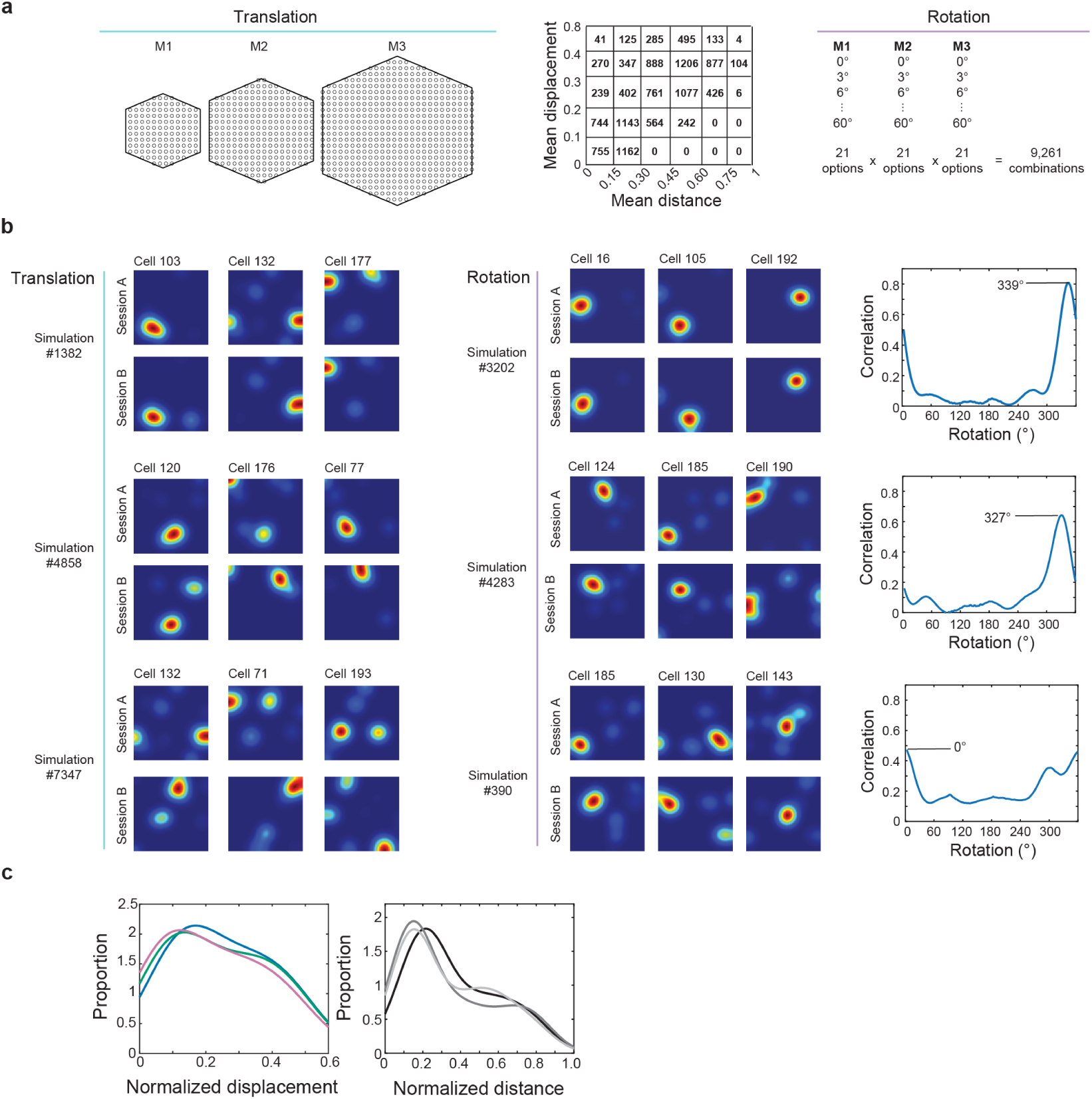
Network model of grid-to-place cell transformation. **a**, Selection of phase and rotation combinations for simulations. Left, Schematic indicates the possible (x,y) locations to which the phase of each module could be shifted during simulations of phase translation in three grid modules (M1, 120 locations; M2, 248 locations; M3, 504 locations). For each module, points were equally spaced throughout the unit hexagon in steps of 4. In total, this resulted in 1.5 × 10^7^ combinations. Middle, we selected 12,300 combinations by varying the mean phase distance between modules and their displacement from the origin. Values indicate the number of simulations in each square. Right, schematic indicates how the 9,261 combinations of module rotation were selected. The rotation of each module varied from 0° to 60° in 3° steps, resulting in 21 possibilities for each module. In total, this resulted in 9,261 (21^3^) possible rotation combinations. **b**, Three representative rate maps of place cells in Session A and Session B from six representative simulations of module translation or rotation. Cell ID is noted above each plot. Left, each pair of rows shows one simulation of module translation (from top: simulations #1382, #4858, and #7347). Minimum pairwise distance between module phase changes increases from top to bottom. Note that as the minimum distance between modules increases (from top to bottom), simulated place cells remap increasingly between sessions. Right, each pair of rows shows one simulation of module rotation (from top: simulation #3202, #4283, and #390). Panels at right show population vector (PV) correlation between sessions for simulated place cells as a function of rotation difference between the modules. Note that as the minimum difference in rotation increases from top to bottom, place cells remap increasingly between sessions, rather than undergoing a coherent rotation. **c**, Left, panel shows the distribution of normalized phase displacement values for each module (M1, blue; M2, green; M3, pink) for 12,300 simulations of module translation. Right, panels shows the distribution of normalized distance between the endpoints of the phase-change vectors for each pair of grid modules (M1-M2, black; M1-M3, dark gray; M2-M3, light gray) for 12,300 simulations of module translation.

**Extended Data Fig. 13.**
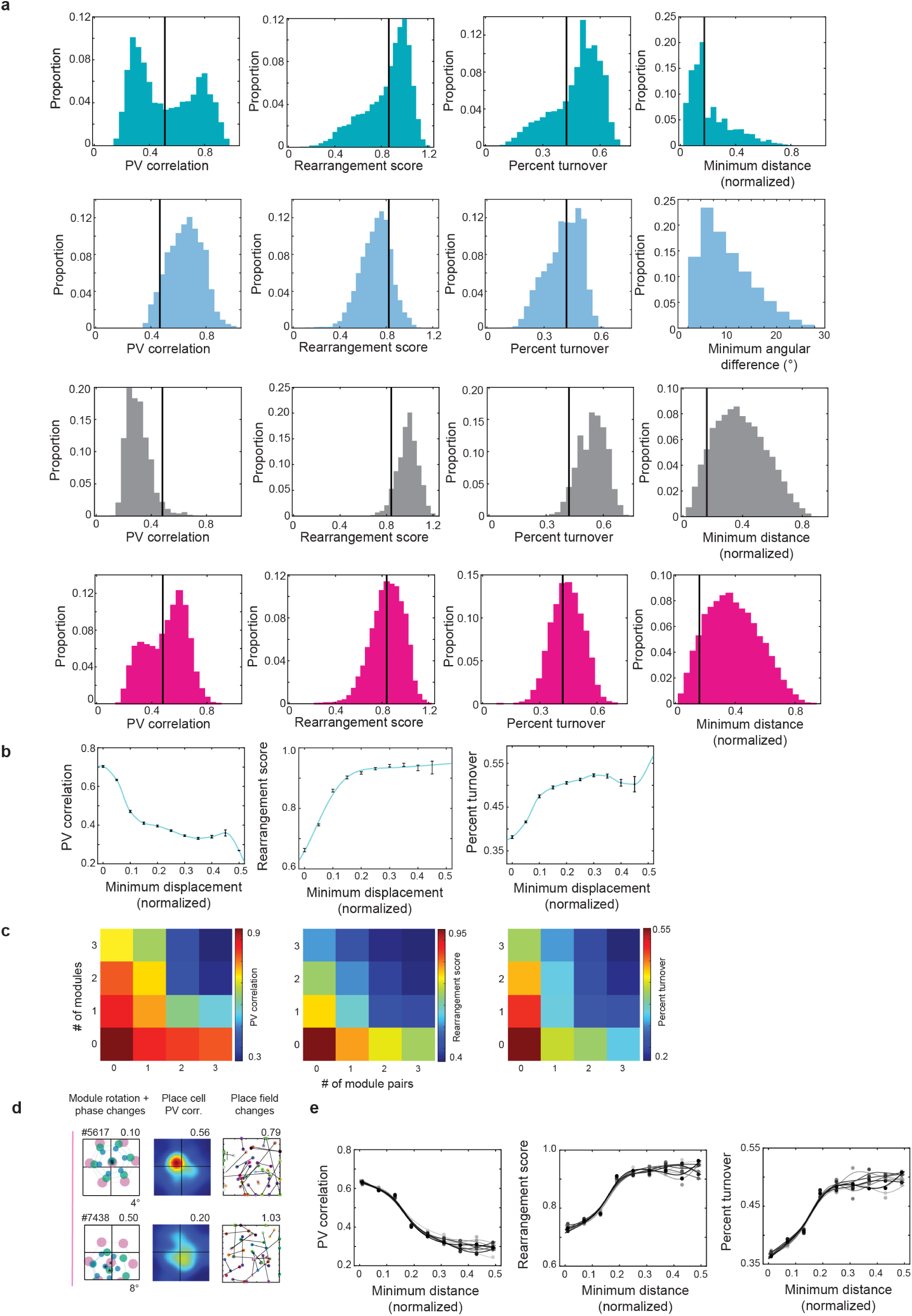
Independent translation, but not rotation, of grid modules is sufficient to produce hippocampal remapping. **a**, Histograms of PV correlations (first column), rearrangement scores (second column), and percent turnover (third column) for all simulations of module translation (top row, green), module rotation (second row, blue), a shuffled distribution (third row, gray), or combined simulations of translation and rotation (fourth row, pink). Shuffled distributions were created by calculating PV correlations, rearrangement scores, and percent turnover between place cells from session A in one simulation and place cells from a randomly selected simulation of session B (n = 1,000 shuffled values for each panel). Black lines indicate the significance threshold for each distribution (95^th^ percentile for PV correlation; 5^th^ percentile for rearrangement score and percent turnover). Fourth column, histogram of normalized minimum distance for simulations of module translation (top row), minimum angular difference between modules for simulations of module rotation (second row), normalized minimum distance for a shuffled distribution (third row) and for simulations of combined module translation and rotation (fourth row). The shuffled distribution was created by randomly shuffling the grid patterns of each module with respect to one another 1,000 times, and calculating the normalized minimum distance between modules. Black lines indicate the significance threshold of this distribution (5^th^ percentile). **b**, PV correlation (left), rearrangement score (middle), and percent turnover (right) as a function of the minimum phase displacement (normalized) of grid modules in the simulations. **c**, Heat maps show PV correlation (left), rearrangement score (middle), and percent turnover (right) as a function of the number of independent module pairs and the number of modules shifted relative to the origin. A module pair was considered to be independent if the pairwise distance between module phase changes was greater than the significance threshold (5^th^ percentile of a shuffled distribution). A module was considered to be shifted if the displacement of the module exceeded the significance threshold (5^th^ percentile of a shuffled distribution). **d**, Left column, schematic representation of module phase changes during representative simulations of combined module rotation and translation (from top: simulation #5617, n = 38 cells; simulation #7438, n = 27 cells). Minimum normalized distance between module phase changes and minimum angular difference between modules are noted above and below each panel, respectively. Middle column, PV crosscorrelograms at the rotation with the highest correlation for simulated place cells during each simulation of combined module rotation and translation. Maximum PV correlation is noted above each panel. Right column, black lines show the change in field location for each simulated place cell when grid cell rate maps for each module were shifted and rotated as indicated in the left column. Rearrangement score is noted above each panel. **e**, PV correlation (left), rearrangement scores (middle), and percent turnover (right) versus normalized minimum distance between module phase changes in the simulations. Color of lines (gray to black) reflects increasing angular difference between modules. Minimum angular difference between modules increased from 2° to 18° in 2° steps.

**Supplementary Table 1.**
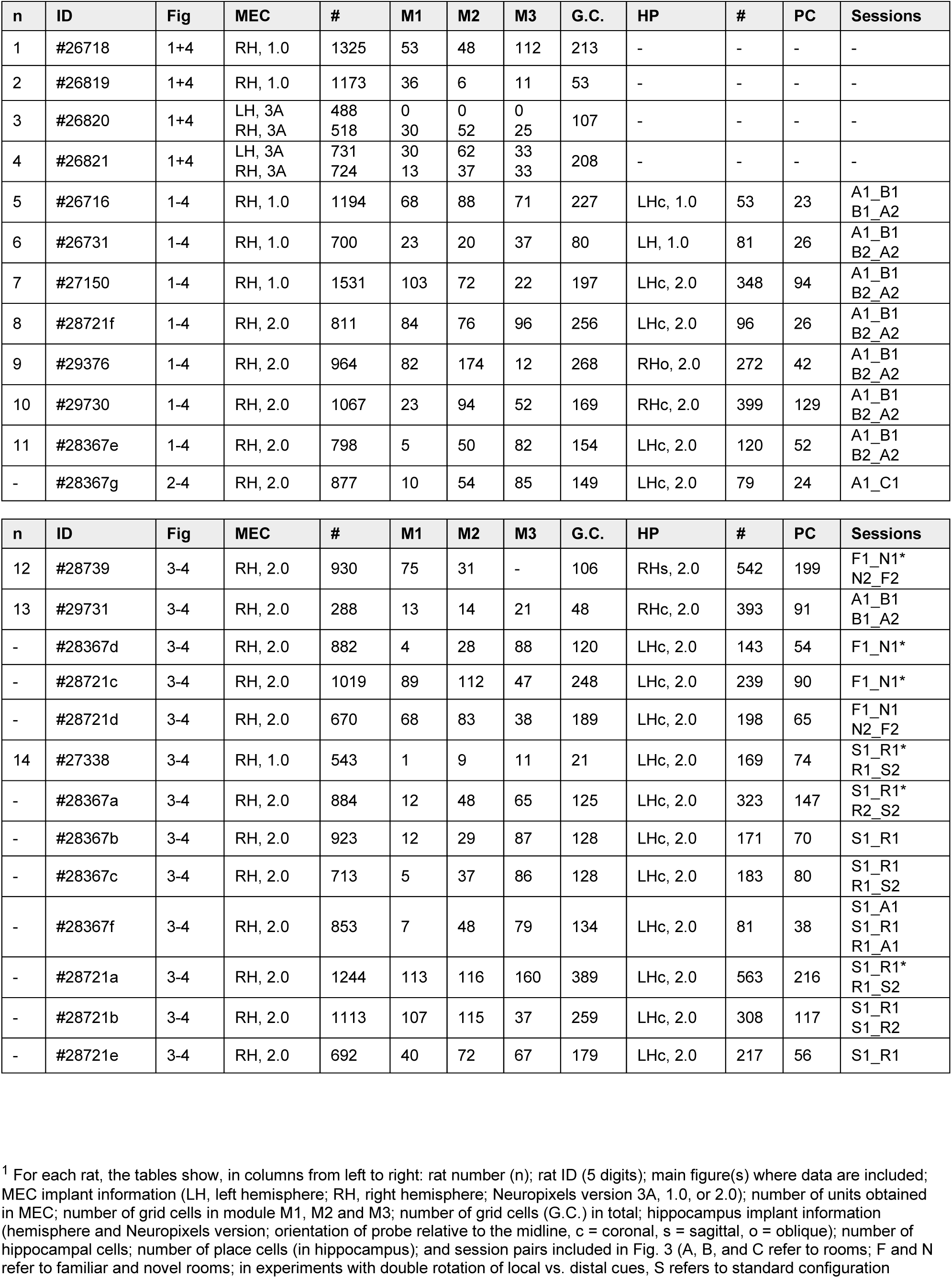
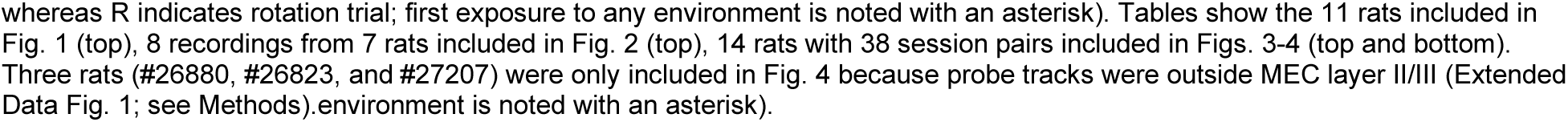
Number of units and sessions for individual rats.^1^.

**Supplementary Table 2.** Changes in the firing properties of cells in MEC/PaS and hippocampus between familiar rooms.^2^.

| MEC/PaS | Grid | Border | HD | Spatial | Other |
| --- | --- | --- | --- | --- | --- |
| # cells | 3,094 | 75 | 1,589 | 2,383 | 15,053 |
| # expts / # rats | 22 expts / 14 rats | 14 expts / 11 rats | 22 expts / 14 rats | 20 expts / 14 rats | 22 expts / 14 rats |
| \| Δ mean rate \| | 0.136; 0.131 - 0.144<br>0.139; 0.132 - 0.147 | 0.13; 0.10 - 0.17<br>0.15; 0.09 - 0.19 | 0.14; 0.13 - 0.15<br>0.18; 0.17 - 0.19 | 0.112; 0.107 - 0.119<br>0.159; 0.149 - 0.166 | 0.162; 0.159 - 0.166<br>0.180; 0.175 - 0.184 |
| \| Δ peak rate \| | 0.132; 0.126 - 0.137<br>0.141; 0.135 - 0.148 | 0.14; 0.10 - 0.18<br>0.17; 0.13 - 0.21 | 0.14; 0.14 - 0.15<br>0.17; 0.16 - 0.18 | 0.135; 0.125 - 0.143<br>0.177; 0.166 - 0.190 | 0.181; 0.177 - 0.184<br>0.209; 0.205 - 0.214 |
| \| Δ size \| | 0.143; 0.135 - 0.151<br>0.196; 0.189 - 0.206 | 0.16; 0.14 - 0.23<br>0.27; 0.23 - 0.31 | 0.20; 0.18 - 0.22<br>0.28; 0.26 - 0.30 | 0.135; 0.127 - 0.142<br>0.171; 0.161 - 0.181 | 0.199; 0.192 - 0.207<br>0.263; 0.256 - 0.272 |
| Spatial correlation | 0.75; 0.74 - 0.76<br>0.002; -0.01 - 0.01 | 0.77; 0.74 - 0.81<br>-0.08; -0.17 - 0.07 | 0.47; 0.45 - 0.49<br>0.07; 0.05 - 0.08 | 0.61; 0.59 - 0.62<br>0.06; 0.05 - 0.08 | 0.209; 0.203 - 0.214<br>0.067; 0.063 - 0.072 |
| Spatial info | 0.77; 0.74 - 0.79<br>0.75; 0.73 - 0.76 | 0.97; 0.81 - 1.03<br>0.99; 0.83 - 1.13 | 0.46; 0.44 - 0.48<br>0.50; 0.48 - 0.52 | 0.51; 0.49 - 0.53<br>0.51; 0.48 - 0.53 | 0.324; 0.312 - 0.331<br>0.397; 0.390 - 0.403 |
| \| Δ functional score \| | 0.04; 0.03 - 0.06<br>0.09; 0.06 - 0.12 | 0.05; 0.01 - 0.10<br>0.01; -0.01 - 0.04 | -0.01; -0.01 - 0.01<br>-0.01; -0.02 - -0.01 |  |  |
| \| Δ grid spacing \| | 0.050; 0.047 - 0.052<br>0.073; 0.069 - 0.075 |  |  |  |  |
| \| Δ ellipticity \| | 0.26; 0.23 - 0.28<br>0.31; 0.29 - 0.33 |  |  |  |  |
| HP | All cells | CA3 | CA1 | DG |  |
| # cells | 416 | 153 | 221 | 42 |  |
| # expts / # rats | 8 expts / 7 rats | 5 expts / 4 rats | 6 expts / 5 rats | 1 expt / 1 rat |  |
| \| Δ mean rate \| | 0.14; 0.12 - 0.15<br>0.26; 0.23 - 0.28 | 0.14; 0.11 - 0.16<br>0.30; 0.26 - 0.34 | 0.14; 0.12 - 0.16<br>0.22; 0.20 - 0.27 | 0.14; 0.11 - 0.26<br>0.32; 0.24 - 0.45 |  |
| \| Δ peak rate \| | 0.16; 0.14 - 0.18<br>0.32; 0.28 - 0.35 | 0.16; 0.14 - 0.20<br>0.40; 0.33 - 0.50 | 0.15; 0.13 - 0.18<br>0.24; 0.20 - 0.28 | 0.18; 0.12 - 0.23<br>0.48; 0.38 - 0.56 |  |
| \| Δ size \| | 0.18; 0.16 - 0.20<br>0.26; 0.23 - 0.30 | 0.16; 0.12 - 0.19<br>0.22; 0.18 - 0.26 | 0.20; 0.16 - 0.25<br>0.30; 0.24 - 0.34 | 0.10; 0.07 - 0.19<br>0.26; 0.18 - 0.46 |  |
| Spatial info | 1.26; 1.16 - 1.30<br>1.07; 0.99 - 1.16 | 1.51; 1.33 - 1.65<br>1.27; 1.11 - 1.47 | 1.14; 1.07 - 1.19<br>0.98; 0.90 - 1.05 | 1.25; 1.11 - 1.52<br>1.12; 0.83 - 1.24 |  |
| Spatial correlation | 0.69; 0.66 - 0.73<br>-0.03; -0.04 - 0.01 | 0.75; 0.69 - 0.78<br>-0.05; -0.06 - -0.03 | 0.66; 0.61 - 0.70<br>-0.01; -0.02 - 0.04 | 0.84; 0.77 - 0.88<br>-0.04; -0.09 - 0.05 |  |
| Place field shift | 5.39; 4.18 - 7.21<br>21.10; 19.42 - 22.80 | 3.16; 2.24 - 4.47<br>20.10; 17.46 - 23.02 | 8.94; 7.04 - 13.00<br>21.54; 19.42 - 24.02 | 2.24; 2.00 - 4.12<br>20.19; 14.27 - 24.63 |  |
| PV correlation<br>(no shift or rotation) | 0.655; 0.649 - 0.663<br>0.076; 0.071 - 0.081 | 0.71; 0.70 - 0.72<br>0.06; 0.05 - 0.06 | 0.62; 0.61 - 0.63<br>0.11; 0.10 - 0.12 | 0.70; 0.69 - 0.71<br>0.04; 0.03 - 0.05 |  |
| PV correlation | 0.52; 0.48 - 0.65<br>0.18; 0.09 - 0.26 | 0.62; 0.60 - 0.65<br>0.26; 0.09 - 0.33 | 0.51; 0.43 - 0.64<br>0.26; 0.13 - 0.28 | 0.62<br>0.07 |  |
| Rearrangement score | 0.49; 0.45 - 0.56<br>0.95; 0.89 - 0.99 | 0.40; 0.21 - 0.57<br>0.99; 0.48 - 1.07 | 0.59; 0.31 - 0.78<br>0.91; 0.76 - 0.99 | 0.56<br>0.92 |  |

<sup>2</sup> Changes in the firing properties of cells in MEC-PaS and hippocampus (HP) between sessions in the same room (A1×A2, top row of each cell) or in different rooms (A1×B1, bottom row of each cell). Table shows medians and 95% confidence intervals. An additional decimal place is shown when necessary for precision. Results of relevant statistical tests are reported in the main text or figure legends. HD, head direction-tuned cells; Spatial, cells with non-periodic spatial firing fields not satisfying criteria for grid cells or border cells. Rows for changes in mean rate, peak rate, field size, and grid spacing show absolute value of difference scores (difference between sessions divided by sum). Size refers to the size of the largest firing field. Spatial information is calculated in sessions A1 (top row of each cell) or B1 (bottom row of each cell). Spatial correlation is between A1×A2 or A1×B1. Changes in functional score refer to (from top to bottom) grid score<sup>71</sup>, border score<sup>77</sup>, and mean vector length for head direction. Changes in functional score and grid ellipticity were calculated as the absolute value of the difference between sessions. Place field shift refers to how far the location of peak firing in the rate map of each cell shifted between sessions (in cm). Note drop in spatial correlation for both grid cells and hippocampal cells, as expected when the grid phase changes and hippocampal cells remap.

**Supplementary Table 3.**
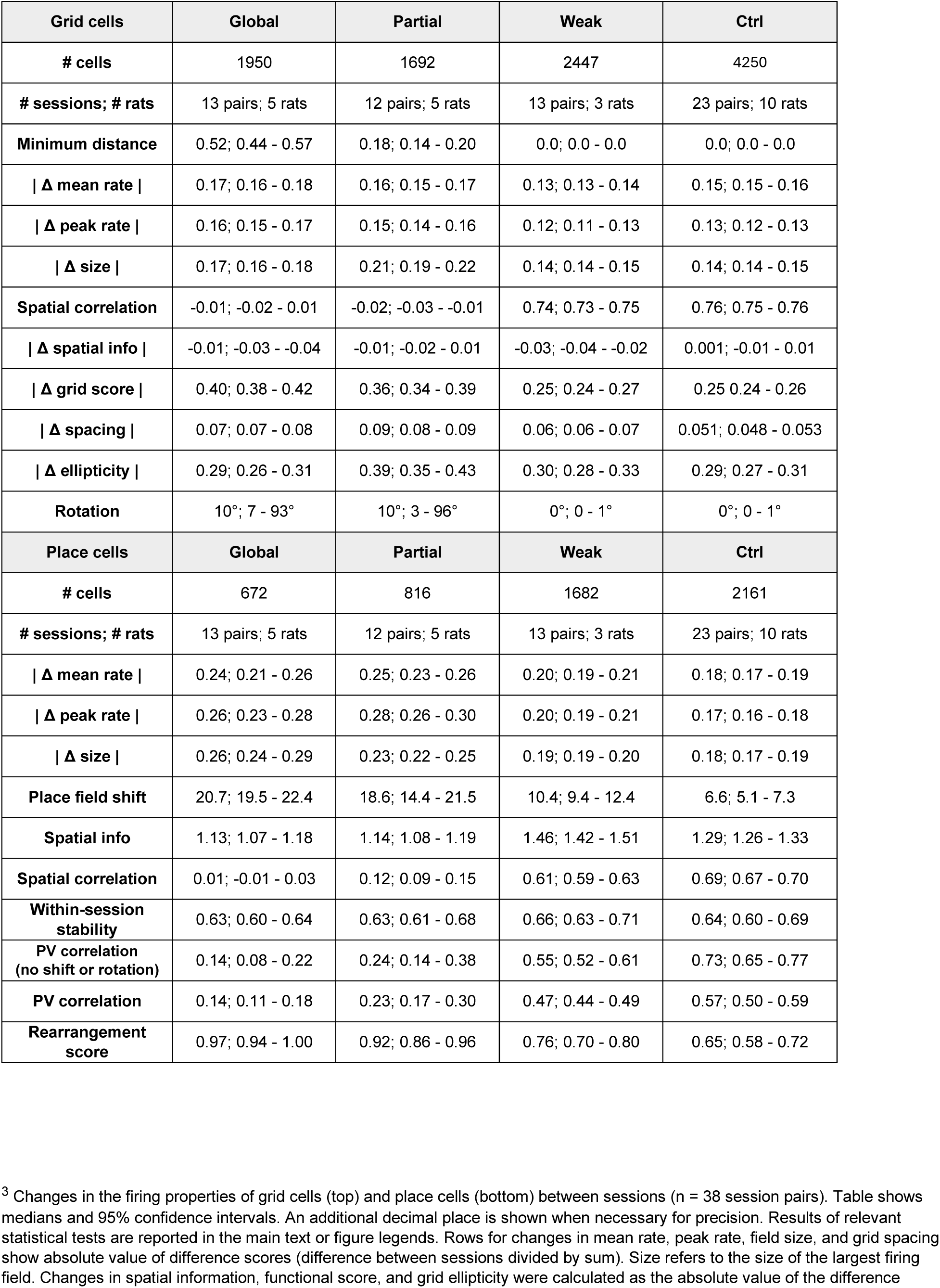
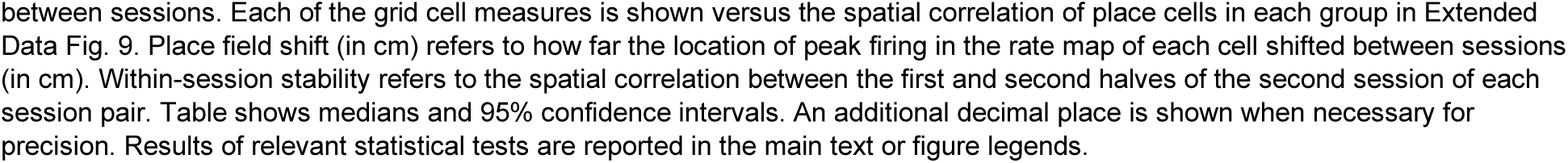
Changes in the firing properties of grid and place cells between sessions.^3^.

## References

1 Taube, J. S., Muller, R. U. & Ranck, J. B., Jr. Head-direction cells recorded from the postsubiculum in freely moving rats. II. Effects of environmental manipulations. J Neurosci 10, 436–447 (1990).

2 Yoganarasimha, D., Yu, X. & Knierim, J. J. Head direction cell representations maintain internal coherence during conflicting proximal and distal cue rotations: comparison with hippocampal place cells. J Neurosci 26, 622–631 (2006). 10.1523/jneurosci.3885-05.2006

3 Fyhn, M., Hafting, T., Treves, A., Moser, M. B. & Moser, E. I. Hippocampal remapping and grid realignment in entorhinal cortex. Nature 446, 190–194 (2007). 10.1038/nature05601

4 Yoon, K. et al. Specific evidence of low-dimensional continuous attractor dynamics in grid cells. Nat Neurosci 16, 1077–1084 (2013). 10.1038/nn.3450

5 Peyrache, A., Lacroix, M. M., Petersen, P. C. & Buzsáki, G. Internally organized mechanisms of the head direction sense. Nature Neuroscience 18, 569–575 (2015). 10.1038/nn.3968

6 Trettel, S. G., Trimper, J. B., Hwaun, E., Fiete, I. R. & Colgin, L. L. Grid cell co-activity patterns during sleep reflect spatial overlap of grid fields during active behaviors. Nat Neurosci 22, 609–617 (2019). 10.1038/s41593-019-0359-6

7 Gardner, R. J., Lu, L., Wernle, T., Moser, M.-B. & Moser, E. I. Correlation structure of grid cells is preserved during sleep. Nat Neurosci 22, 598–608 (2019). 10.1038/s41593-019-0360-0

8 Gardner, R. J. et al. Toroidal topology of population activity in grid cells. Nature 602, 123–128 (2022). 10.1038/s41586-021-04268-7

9 Muller, R. U. & Kubie, J. L. The effects of changes in the environment on the spatial firing of hippocampal complex-spike cells. J Neurosci 7, 1951–1968 (1987).

10 Bostock, E., Muller, R. & Kubie, J. Experience-dependent modifications of hippocampal place cell firing. Hippocampus 1, 193–206 (1991).

11 Leutgeb, S. et al. Independent codes for spatial and episodic memory in hippocampal neuronal ensembles. Science (New York, N.Y.) 309, 619–623 (2005). 10.1126/science.1114037

12 Alme, C. B. et al. Place cells in the hippocampus: Eleven maps for eleven rooms. Proceedings of the National Academy of Sciences of the United States of America 111, 18428–18435 (2014). 10.1073/pnas.1421056111

13 McNaughton, B. L. & Morris, R. G. Hippocampal synaptic enhancement and information storage within a distributed memory system. Trends in Neurosciences 10, 408–415 (1987). 10.1016/0166-2236(87)90011-7

14 Colgin, L. L., Moser, E. I. & Moser, M. B. Understanding memory through hippocampal remapping. Trends Neurosci 31, 469–477 (2008). 10.1016/j.tins.2008.06.008

15 O’Keefe, J. & Burgess, N. Geometric determinants of the place fields of hippocampal neurons. Nature 381, 425–428 (1996). 10.1038/381425a0

16 Shapiro, M., et al. Cues that hippocampal place cells encode: Dynamic and hierarchical representation of local and distal stimuli. Hippocampus 7(**6**), 624–642 (1997).

17 Tanila, H., et al. Discordance of spatial representation in ensembles of hippocampal place cells. Hippocampus 7(**6**), 613–623 (1997).

18 Knierim, J. J. Dynamic interactions between local surface cues, distal landmarks, and intrinsic circuitry in hippocampal place cells. J Neurosci 22, 6254–6264 (2002). 20026608

19 Anderson, M. I. & Jeffery, K. J. Heterogeneous modulation of place cell firing by changes in context. J Neurosci 23, 8827–8835 (2003).

20 Treves, A. & Rolls, E. T. What determines the capacity of autoassociative memories in the brain? Network: Computation in Neural Systems 2, 371–397 (1991). 10.1088/0954-898X_2_4_004

21 Monaco, J. D. & Abbott, L. F. Modular realignment of entorhinal grid cell activity as a basis for hippocampal remapping. J Neurosci 31, 9414–9425 (2011). 10.1523/jneurosci.1433-11.2011

22 Chandra, S., Sharma, S., Chaudhuri, R. & Fiete, I. Episodic and associative memory from spatial scaffolds in the hippocampus. Nature 638, 739–751 (2025). 10.1038/s41586-024-08392-y

23 Hubel, D. H. & Wiesel, T. N. Receptive fields, binocular interaction and functional architecture in the cat’s visual cortex. J Physiol 160, 106–154 (1962). 10.1113/jphysiol.1962.sp006837

24 Barlow, H. B., Blakemore, C. & Pettigrew, J. D. The neural mechanism of binocular depth discrimination. J Physiol 193, 327–342 (1967). 10.1113/jphysiol.1967.sp008360

25 Poggio, G., Gonzalez, F. & Krause, F. Stereoscopic mechanisms in monkey visual cortex: binocular correlation and disparity selectivity. J Neurosci 8, 4531–4550 (1988). 10.1523/jneurosci.08-12-04531.1988

26 Carr, C. E. & Konishi, M. A circuit for detection of interaural time differences in the brain stem of the barn owl. J Neurosci 10, 3227 (1990). 10.1523/JNEUROSCI.10-10-03227.1990

27 Rust, N. C., Mante, V., Simoncelli, E. P. & Movshon, J. A. How MT cells analyze the motion of visual patterns. Nat Neurosci 9, 1421–1431 (2006). 10.1038/nn1786

28 Seelig, J. D. & Jayaraman, V. Neural dynamics for landmark orientation and angular path integration. Nature 521, 186–191 (2015). 10.1038/nature14446

29 Green, J. et al. A neural circuit architecture for angular integration in Drosophila. Nature 546, 101–106 (2017). 10.1038/nature22343

30 Fisher, Y. E., Lu, J., D’Alessandro, I. & Wilson, R. I. Sensorimotor experience remaps visual input to a heading-direction network. Nature 576, 121–125 (2019). 10.1038/s41586-019-1772-4

31 Lyu, C., Abbott, L. F. & Maimon, G. Building an allocentric travelling direction signal via vector computation. Nature 601, 92–97 (2022). 10.1038/s41586-021-04067-0

32 Mussells Pires, P., Zhang, L., Parache, V., Abbott, L. F. & Maimon, G. Converting an allocentric goal into an egocentric steering signal. Nature 626, 808–818 (2024). 10.1038/s41586-023-07006-3

33 Westeinde, E. A. et al. Transforming a head direction signal into a goal-oriented steering command. Nature 626, 819–826 (2024). 10.1038/s41586-024-07039-2

34 Jun, J. J. et al. Fully integrated silicon probes for high-density recording of neural activity. Nature 551, 232–236 (2017). 10.1038/nature24636

35 Steinmetz, N. A. et al. Neuropixels 2.0: A miniaturized high-density probe for stable, long-term brain recordings. Science (New York, N.Y.) 372, eabf4588 (2021). 10.1126/science.abf4588

36 Fyhn, M., Molden, S., Witter, M. P., Moser, E. I. & Moser, M. B. Spatial representation in the entorhinal cortex. Science (New York, N.Y.) 305, 1258–1264 (2004). 10.1126/science.1099901

37 Hafting, T., Fyhn, M., Molden, S., Moser, M. B. & Moser, E. I. Microstructure of a spatial map in the entorhinal cortex. Nature 436, 801–806 (2005). 10.1038/nature03721

38 Barry, C., Hayman, R., Burgess, N. & Jeffery, K. J. Experience-dependent rescaling of entorhinal grids. Nat Neurosci 10, 682–684 (2007). http://www.nature.com/neuro/journal/v10/n6/suppinfo/nn1905_S1.html

39 Stensola, H. et al. The entorhinal grid map is discretized. Nature 492, 72–78 (2012). 10.1038/nature11649

40 Waaga, T. et al. Grid-cell modules remain coordinated when neural activity is dissociated from external sensory cues. Neuron 110, 1843–1856.e1846 (2022). 10.1016/j.neuron.2022.03.011

41 Solstad, T., Moser, E. I. & Einevoll, G. T. From grid cells to place cells: a mathematical model. Hippocampus 16, 1026–1031 (2006). 10.1002/hipo.20244

42 Fuhs, M. C. & Touretzky, D. S. A spin glass model of path integration in rat medial entorhinal cortex. J Neurosci 26, 4266–4276 (2006). 10.1523/JNEUROSCI.4353-05.2006

43 McNaughton, B. L., Battaglia, F. P., Jensen, O., Moser, E. I. & Moser, M.-B. Path integration and the neural basis of the ‘cognitive map’. Nat Rev Neurosci 7, 663–678 (2006).

44 Rolls, E. T., Stringer, S. M. & Elliot, T. Entorhinal cortex grid cells can map to hippocampal place cells by competitive learning. Network 17, 447–465 (2006). 10.1080/09548980601064846

45 Si, B. & Treves, A. The role of competitive learning in the generation of DG fields from EC inputs. Cogn Neurodyn 3, 177–187 (2009). 10.1007/s11571-009-9079-z

46 de Almeida, L., Idiart, M. & Lisman, J. E. The input-output transformation of the hippocampal granule cells: from grid cells to place fields. J Neurosci 29, 7504–7512 (2009). 10.1523/jneurosci.6048-08.2009

47 Savelli, F. & Knierim, J. J. Hebbian analysis of the transformation of medial entorhinal grid-cell inputs to hippocampal place fields. J Neurophysiol 103, 3167–3183 (2010). 10.1152/jn.00932.2009

48 Fiete, I. R., Burak, Y. & Brookings, T. What grid cells convey about rat location. J Neurosci 28, 6858–6871 (2008). 10.1523/jneurosci.5684-07.2008

49 Sreenivasan, S. & Fiete, I. Grid cells generate an analog error-correcting code for singularly precise neural computation. Nat Neurosci 14, 1330–1337 (2011). 10.1038/nn.2901

50 McInnes, L. & Healy, J. UMAP: Uniform Manifold Approximation and Projection for Dimension Reduction. ArXiv abs/1802.03426 (2018).

51 Kanter, B. R. et al. A novel mechanism for the grid-to-place cell transformation revealed by transgenic depolarization of medial entorhinal cortex layer II. Neuron 93, 1480–1492 (2017). 10.1016/j.neuron.2017.03.001

52 Diehl, G. W., Hon, O. J., Leutgeb, S. & Leutgeb, J. K. Grid and nongrid cells in medial entorhinal cortex represent spatial location and environmental features with complementary coding schemes. Neuron 94, 83–92 (2017). 10.1016/j.neuron.2017.03.004

53 Butler, W. N., Hardcastle, K. & Giocomo, L. M. Remembered reward locations restructure entorhinal spatial maps. Science (New York, N.Y.) 363, 1447–1452 (2019). 10.1126/science.aav5297

54 de Almeida, L., Idiart, M. & Lisman, J. E. The single place fields of CA3 cells: a two-stage transformation from grid cells. Hippocampus 22, 200–208 (2012). 10.1002/hipo.20882

55 Guanella, A., Kiper, D. & Verschure, P. A model of grid cells based on a twisted torus topology. Int J Neural Syst 17, 231–240 (2007). 10.1142/s0129065707001093

56 Burak, Y. & Fiete, I. R. Accurate Path Integration in Continuous Attractor Network Models of Grid Cells. PLOS Computational Biology 5, e1000291 (2009). 10.1371/journal.pcbi.1000291

57 Gothard, K. M., Skaggs, W. E., Moore, K. M. & McNaughton, B. L. Binding of hippocampal CA1 neural activity to multiple reference frames in a landmark-based navigation task. J Neurosci 16, 823–835 (1996).

58 Parsa, N., Matthew, A. W. & Honi, S. Animal-to-Animal Variability in Partial Hippocampal Remapping in Repeated Environments. The Journal of Neuroscience 42, 5268 (2022). 10.1523/JNEUROSCI.3221-20.2022

59 Fenton, A. A., Wesierska, M., Kaminsky, Y. & Bures, J. Both here and there: Simultaneous expression of autonomous spatial memories in rats. Proceedings of the National Academy of Sciences 95, 11493–11498 (1998). 10.1073/pnas.95.19.11493

60 Leutgeb, J. K. et al. Progressive transformation of hippocampal neuronal representations in “morphed” environments. Neuron 48, 345–358 (2005). 10.1016/j.neuron.2005.09.007

61 Wills, T. J., Lever, C., Cacucci, F., Burgess, N. & O’Keefe, J. Attractor dynamics in the hippocampal representation of the local environment. Science (New York, N.Y.) 308, 873–876 (2005). 10.1126/science.1108905

62 Jezek, K., Henriksen, E. J., Treves, A., Moser, E. I. & Moser, M. B. Theta-paced flickering between place-cell maps in the hippocampus. Nature 478, 246–249 (2011). 10.1038/nature10439

63 Kelemen, E. & Fenton, A. A. Dynamic grouping of hippocampal neural activity during cognitive control of two spatial frames. PLoS Biol 8, e1000403 (2010). 10.1371/journal.pbio.1000403

64 Tulving, E. in Organization of memory. xiii, 423–xiii, 423 (Academic Press, 1972).

65 Squire, L. R. & Zola-Morgan, S. The medial temporal lobe memory system. Science (New York, N.Y.) 253, 1380–1386 (1991). 10.1126/science.1896849

66 Eichenbaum, H., Dudchenko, P., Wood, E., Shapiro, M. & Tanila, H. The hippocampus, memory, and place cells: is it spatial memory or a memory space? Neuron 23, 209–226 (1999).

67 Buzsáki, G. & Moser, E. I. Memory, navigation and theta rhythm in the hippocampal-entorhinal system. Nature Neuroscience 16, 130–138 (2013). 10.1038/nn.3304

68 Lee, I., Yoganarasimha, D., Rao, G. & Knierim, J. J. Comparison of population coherence of place cells in hippocampal subfields CA1 and CA3. Nature 430, 456–459 (2004). 10.1038/nature02739

69 Skaggs, W., Mcnaughton, B. & Gothard, K. An information-theoretic approach to deciphering the hippocampal code. Advances in Neural Information Processing Systems 5 (1992).

## Methods References

70 Vollan, A. Z., Gardner, R. J., Moser, M.-B. & Moser, E. I. Left–right-alternating theta sweeps in entorhinal–hippocampal maps of space. Nature 639, 995–1005 (2025). 10.1038/s41586-024-08527-1

71 Sargolini, F. et al. Conjunctive representation of position, direction, and velocity in entorhinal cortex. Science (New York, N.Y.) 312, 758–762 (2006). 10.1126/science.1125572

72 Wernle, T. et al. Integration of grid maps in merged environments. Nat Neurosci 21, 92–101 (2018). 10.1038/s41593-017-0036-6

73 Krupic, J., Bauza, M., Burton, S., Barry, C. & O’Keefe, J. Grid cell symmetry is shaped by environmental geometry. Nature 518, 232–235 (2015). 10.1038/nature14153

74 Stensola, T., Stensola, H., Moser, M. B. & Moser, E. I. Shearing-induced asymmetry in entorhinal grid cells. Nature 518, 207–212 (2015). 10.1038/nature14151

75 Boccara, C. N., Nardin, M., Stella, F., O’Neill, J. & Csicsvari, J. The entorhinal cognitive map is attracted to goals. Science (New York, N.Y.) 363, 1443 (2019). 10.1126/science.aav4837

76 Langston, R. F. et al. Development of the spatial representation system in the rat. Science (New York, N.Y.) 328, 1576–1580 (2010). 10.1126/science.1188210

77 Solstad, T., Boccara, C. N., Kropff, E., Moser, M. B. & Moser, E. I. Representation of geometric borders in the entorhinal cortex. Science (New York, N.Y.) 322, 1865–1868 (2008). 10.1126/science.1166466

78 Naber, P. A., Lopes da Silva, F. H. & Witter, M. P. Reciprocal connections between the entorhinal cortex and hippocampal fields CA1 and the subiculum are in register with the projections from CA1 to the subiculum. Hippocampus 11, 99–104 (2001). 10.1002/hipo.1028

79 Kloosterman, F., Witter, M. P. & Van Haeften, T. Topographical and laminar organization of subicular projections to the parahippocampal region of the rat. Journal of Comparative Neurology 455, 156–171 (2003). 10.1002/cne.10472

80 Ohara, S. et al. Hippocampal-medial entorhinal circuit is differently organized along the dorsoventral axis in rodents. Cell reports 42, 112001 (2023). 10.1016/j.celrep.2023.112001

